# Roads as ecological traps for giant anteaters

**DOI:** 10.1101/2021.04.02.438243

**Authors:** Michael J. Noonan, Fernando Ascensão, Débora R. Yogui, Arnaud L.J. Desbiez

## Abstract

Wildlife-vehicle collisions (WVC) represent a serious source of mortality for many species, threatening local populations’ persistence while also carrying a high economic and human safety cost. Animals may adapt their behaviour to road associated threats, but roadside resources can act as attractants, providing misleading signals about the quality of roadside habitats, ultimately acting as an ecological trap. Yet, the extent to which individuals modify their behaviour and space use to roads is largely unknown for most taxa. Using fine-scale movement data from 41 giant anteaters (*Myrmecophaga tridactyla*), tracked in the Brazilian Cerrado, we aimed to identify facets of movement behaviour that might exhibit plasticity to roads and traffic volume. Specifically, the analysis of daily and instantaneous movement speeds, home range characteristics, and crossing rates/times, allowed us to test for an effect of road proximity, traffic volume and natural linear features on movement behaviour. We found no effect of road proximity or traffic volume on space use or movement behaviour. While individuals tended to reduce their movement speed when approaching roads and crossed roads ~3 times less than would have been expected by random chance, none of the highways we monitored were impervious. The majority of tracked anteaters living near roads (<2km) crossed them, with higher crossing rates for males. Consequently, habitat near roads may function as an ecological trap where healthy individuals occupy the territories nearby or bisected by roads but eventually are road-killed given their regular crossings, leaving the territory vacant for subsequent occupation. Crucially, we found no evidence that anteaters actively searched for passage structures to cross the roads. This suggests that crossing structures alone are unlikely to mitigate WVC induced mortality. Our research reinforces the need to implement fencing, properly linked to existing passages, and minimising the amount of night-time driving to reduce the number of WVCs.

**Article Impact Statement:** Analyses of giant anteater movement show how roads act as ecological traps, reinforcing the need for fencing and reduced night-time driving.

## Introduction

Human development has been reducing the amount of habitat available to wildlife (Venter et al. 2016). Animals moving through these altered landscapes are coming into conflict with humans at rapidly increasing rates (Fahrig 2007; Macdonald 2016; Buchholtz et al. 2020). Of special concern are the impacts of roads on biodiversity (Forman et al. 2003; Van der Ree et al. 2015). In particular, wildlife-vehicle collisions (WVC) that occur while animals try to move across roads, represent a serious source of mortality for many species (D’Amico et al. 2015; González-Suárez et al. 2018; Ascensão et al. 2019a). WVC not only threaten the local population persistence (Desbiez et al. 2020), but also carry a high economic and human safety cost. In Brazil, for example, the annual cost in vehicle damage due to WVC was estimated as being between $1,171 and $1,492 USD per km of paved road, in Mato Grosso do Sul (Ascensão et al. 2021); and in São Paulo State, the total annual cost of WVC to society was estimated at $25 million USD (Abra et al. 2019). Beyond the direct impact of WVC, roads can also hinder species’ capacities to disperse and re-distribute (Clark et al. 2010; Long et al. 2010), potentially reducing gene flow and, consequently, their population viability (Riley et al. 2006; Ceia-Hasse et al. 2018).

Because of the detrimental impacts that roads have on many species (Fahrig & Rytwinski 2009), individuals living in road-bisected habitats are expected to respond by adapting their behaviour to the unique requirements of roadside environments (Ascensão et al. 2017). Problematically, however, roadside foraging opportunities can act as attractants for many species (Barrientos & Bolonio 2009; Ascensão et al. 2012), providing misleading information about the quality of roadside habitats. When species do not have the inherent capacity to learn to avoid oncoming vehicles, this attraction can act as an ecological trap (Schlaepfer et al. 2002) that may carry severe consequences, especially for *K*-selected species. Behavioural plasticity towards roads is especially important for long-lived species (Sih et al. 2011; Montgomery et al. 2020) that take longer to reach sexual maturity, and have longer interbirth intervals than short-lived species (De Magalhaes & Costa 2009), making it difficult for them to compensate for human-induced mortality. Yet, the extent to which individuals modify their movement behaviour and space use to roads is largely unknown for most taxa – despite such information being critical in both the development of species and landscape management programs and infrastructure development (D’Amico et al. 2016).

Here, we address this research gap by identifying facets of animal movement behaviour that might exhibit plasticity in response to roads. We base our study on giant anteaters (*Myrmecophaga tridactyla*), the largest extant anteater. Giant anteaters reach over two meters and weight up to 50 kg (McNab 1984), and are distributed throughout Central and South America (Gardner 2008). Anteater populations have suffered severe reductions with local and regional extirpations and are currently classified as Vulnerable to Extinction (IUCN 2014).

Moreover, WVC are a major threat to giant anteaters, as this species is commonly reported as one of the top mammals recorded in systematic roadkill surveys, reaching an annual rate of ~17 ind./100 km/year in Mato Grosso do Sul, Brazil (Ascensão et al. 2021). Such high non-natural mortality is thought to reduce the viability of populations (Desbiez et al. 2020), given their low recruitment (about one pup per year; Gaudin et al. 2018), and low densities (generally < 1 ind/km^2^) (Bertassoni et al. (in press)). Consequently, reducing WVC induced mortality is recognised a conservation priority for the species, including for the IUCN (Miranda et al. 2014). Despite this recognition, however, there is no information on whether roadside residents regularly cross highways, or if dispersing individuals make up the bulk road-killed animals, nor how anteaters respond to different types of roads and traffic volumes. Similarly, giant anteaters are known to use road passage structures (Abra et al. 2020), but there is no evidence as to whether they search for existing structures to safely cross roads, or if these are only used opportunistically.

Understanding how the movement ecology of giant anteaters responds to roads and its relationship with roadkill can thus provide valuable information for landscape and road management. Such information is even more pressing given the increasing agribusiness expansion, infrastructure development and low legal protection for this species’ habitat (Strassburg et al. 2017). We carried out the most extensive telemetry study on giant anteaters to date to fill these critical knowledge gaps. In particular, we aimed to address four over-arching questions: Q_1_) Does the movement behaviour of giant anteaters differ when living near paved roads? Q_2_) Does traffic volume influence giant anteater crossing behaviour? Q_3_) Do anteaters prefer to cross the roads via passage structures? Q_4_) Do anteaters respond to roads differently than to natural barriers?

Very little is known about the movement ecology of giant anteaters (Medri & Mourão 2005; see e.g.; Bertassoni et al. 2017), but because of the high number of giant anteaters found in road kill surveys (Ascensão et al. 2021) relative to their low population densities (Bertassoni et al. (in press)), our underlying hypothesis was that roads do not significantly deter giant anteaters. As such, we expected to observe no differences in movement behaviour with road proximity or traffic volumes and similar road crossing rates across the different types of roads and natural linear features, such as streams. Likewise, the use of road passage structures (e.g., culverts, viaducts, etc.) was expected to be opportunistic. Findings are directly applicable to developing road and landscape management plans, not only for giant anteaters but to medium-large mammals inhabiting road bisected ecosystems generally.

## Materials and methods

### Study area

The study was conducted in the state of Mato Grosso do Sul (MS), in the Cerrado biome (savannah) of Brazil (Fig.1). Three paved, two lane highways of different ages and traffic volumes cross the study area (Fig.1, Table 1). A network of unpaved dirt roads was also present, linking main roads and ranches (Fig.1). These roads had significantly lower traffic volumes (<1 car per hour) in comparison with paved roads (pers. observations). The land use bordering all roads was dominated by pasture, with sparse remnants of native forest and savanna, and some areas of eucalyptus plantation (Fig.1). Streams bordered with native riparian vegetation were common throughout the study area, mostly accompanying native savanna vegetation. Each study area had 9-10 passages (culverts, box-culverts and viaducts) near the territories of the tracked anteaters (Fig.1; Table 1). The climate throughout MS is wet from October to March and dry from April to September (Koppen’s As or Aw), with mild year-round temperatures (range 21-32°C). Average annual rainfall ranges between 1000-1500mm.

**Figure 1.**
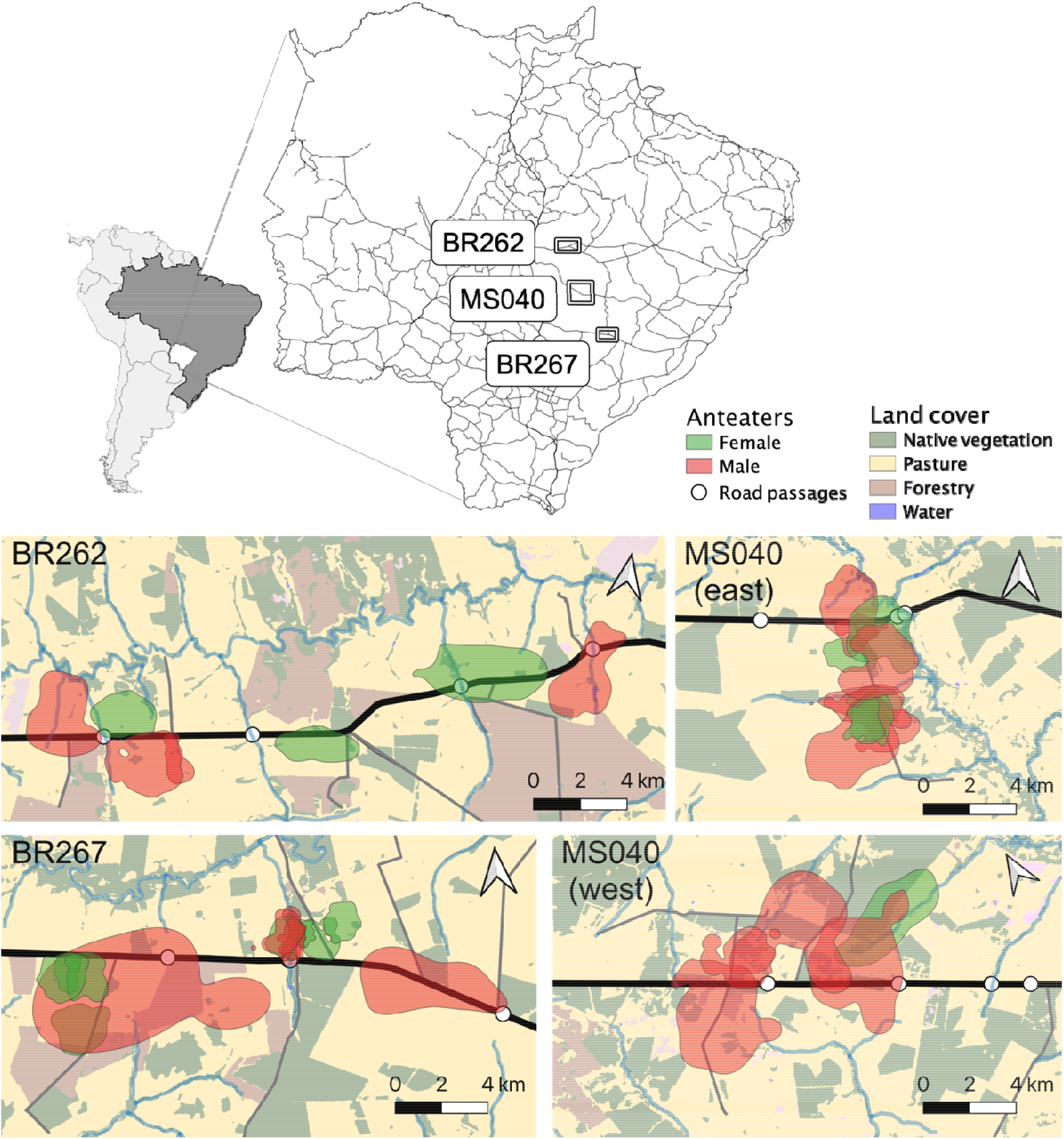
Map showing the location of the three study areas (BR262, MS040 and BR267) in Mato Grosso do Sul, Brazil (top), with state main road network (black lines). In each study area, home ranges are in red for males and green for females (see text for details); the main paved road is depicted in thick black line and other unpaved roads are in thinner black lines, and road passage location are white circles (some circles indicate more than one passage). Land cover classes were from MapBiomas (2018 version).

**Table 1.** Age and mean daily traffic volume (MDT) with the proportion of trucks and cars, for the three main roads present in the study area. Traffic volume information for BR267 and BR267 was obtained from official counts (DNIT 2020). MS040 counts was obtained using the same methodology used by governmental authorities (DNIT 2020), i.e. vehicle counting throughout a full week (7 days, 24 h per day). The ‘Passages’ column indicates the type of passage structures in the study areas BC–box-culvert, C–culvert, V–viaduct (numbers in parentheses are the width of the passages, in m).

| Study area | Road age (years) | MDT | Trucks (%) | Cars (%) | Passages |  |  |
| --- | --- | --- | --- | --- | --- | --- | --- |
|  |  |  |  |  | C | BC | V |
| MS040 | 6 | 603 | 40 | 33 | 2 (1.6m) | 6 (4.0-9.5m) | 1 (50m) |
| BR262 | 34 | 3206 | 32 | 66 | 3 (2.7-3.9m) | 7 (4.0-6.5m) | 1 (25m) |
| BR267 | 34 | 4285 | 46 | 49 | 9 (1.3-2.7m) | - | - |

### Giant anteater GPS data collection

Wild giant anteaters were captured between 2017-18, in the vicinity of the three paved highways of the study area, far from urban areas (>15km), and equipped with a GPS tracking collars. The capture team was always comprised of two veterinarians and a biologist, minimum. To capture giant anteaters, we searched in open areas for animals foraging, during colder months (May-August) when individuals are known to exhibit greater diurnal activity (Camilo◻Alves & Mourão 2006). When an adult was spotted, two members of the capture team approached the individual by foot and captured it using two long-handled-dip-nets (handle 1.5m; hoop 0.7m diameter) to restrain and immobilize it. The veterinarian was then able to safely apply an intramuscular combination anaesthetic injection of butorphanol tartrate (0.1mg/kg), detomidine hydrochloride (0.1mg/kg) and midazolam hydrochloride (0.2mg/kg) into its hind limbs (Kluyber et al. 2020). After anaesthetic induction, the front claws were first wrapped and completely immobilized using tape. Physical exams were then performed to evaluate health conditions and included measuring weight, pregnancy detection (palpation), general appearance, hydration status, mucous membrane colour, respiratory auscultation, and presence of scars or wounds (based on Kluyber et al. 2020). Only adult individuals considered in good health by the veterinaries were fitted with the GPS harness (TGW-4570-4 Iridium GPS) and VHF transmitters (MOD 400; Telonics, Mesa, Arizona). For anaesthetic reversal procedures, all individuals received a combination of three antagonists: naloxone hydrochloride (0.02mg/kg), yohimbine hydrochloride (0.125mg/kg), and Flumazenil (0.01mg/kg) (Kluyber et al. 2020). After the procedure, the animal was maintained in a wooden ventilated crate until complete recovery and was then released at its capture location. Collared anteaters were recaptured approximately one year after for harness removal and data download, but each animal was visually inspected at least once every two weeks for a general health check. The trackers took GPS fixes with 20 min intervals.

### Data analysis

#### Movement data pre-processing

Before analysis, we performed a data cleaning process in order to calibrate the GPS measurement error and filter outliers using the methods implemented in the R package ‘ctmm’ (Calabrese et al. 2016; Fleming & Calabrese 2020), see Appendix S1). For each location estimate, the GPS trackers recorded a unitless Horizontal Dilution of Precision (HDOP), value which is a measure of the accuracy of each positional fix. We converted the HDOP values into calibrated error circles by estimating an equivalent range error from 6,948 calibration data points where a tag had been left in a fixed location (Fleming et al. 2020). For each individual dataset, we then removed outliers based on error-informed distance from the median location, and the minimum speed required to explain each location’s displacement.

#### Movement metrics

For each giant anteater, we quantified key movement metrics and home range related characteristics that allowed us to test for an effect of road proximity, traffic volume and natural linear features on giant anteater movement behaviour (see Appendix S2). First, using the R package ‘ctmm’, we fitted a series of continuous-time movement models (hereafter ‘CTMM’) to the track data, using perturbative-Hybrid Residual Maximum Likelihood (pHREML; Fleming et al. 2019), and identified the best CTMM via small-sample-sized corrected Akaike’s Information Criterion (AICc).

From each CTMM we estimated both the mean ***Daily movement speed*** (in km/day) and the ***Instantaneous movement speed*** (in m/s), using continuous-time speed and distance estimation (Noonan et al. 2019). This approach is insensitive to the sampling schedule, and corrects for GPS measurement error, enabling robust comparisons across individuals.

We next estimated the ***Home-range*** for each animal, as the polygon delimited by the 95% isopleth of the Utilization Distribution using Autocorrelated Kernel Density Estimation (AKDE) (Fleming et al. 2015). AKDE home-range estimates were conditioned on the autocorrelation structure of the CTMM and we implemented the small-sample-size bias correction of Fleming & Calabrese (2017). For each individual, we quantified the land cover within their home range polygon to control for possible confounding effects related to habitat quality that could influence movement behaviour. Land cover was obtained from MapBiomas (Souza & Azevedo 2017) for 2018. We compared the proportion of cover of the two main classes occurring within home ranges, namely pasture and native vegetation. Because of a strong negative correlation between the proportion of pasture and native vegetation in an individual’s home range (Pearson correlation = −0.92), only the proportion of pasture cover was included in our analyses. We also calculated the Euclidean distance between each individual’s home-range centroid and the nearest paved road.

We further estimated the ***total number of crossings*** across the highways, unpaved roads, and streams for each giant anteater. For this, we used each animal’s tracking data and their CTMM to reconstruct the most likely path that they travelled through the landscape. We identified the total number of intersections (crossings) between the most likely path and the different linear features.

#### Movement pattern comparisons

We used the information obtained from each animal’s GPS location data to address each of our four core research questions. The R code required to reproduce these analyses, is presented in Appendix S2.

##### Q1) Does the movement behaviour of giant anteaters differ when living near paved roads?

In order to answer Q_1_, we tested for any relationship between the distance animals lived from roads and i) individual home range areas, ii) daily movement speeds, and iii) instantaneous movement speeds. We also looked at whether there were any differences in these measures across each of the three paved roads to test for an effect of traffic volume. Home-range areas and daily movement speeds were analysed using the meta-regression model implemented in the R package ‘metafor’ (Viechtbauer 2010), which allowed uncertainty in each individual estimate to be propagated into the population level estimate when making comparisons. Instantaneous speeds were analysed using a Gaussian mixed effects model that included the quadratic effects of distance to road and time of day (to control for circadian rhythms), and the nearest paved road (to control for traffic volume). We also included a random intercept for each animal and random slope for the relationship between movement speed and the distance to the road for each individual anteater. Finally, we applied a first order auto-regressive correction to the residuals to account for autocorrelation in instantaneous movement speeds. The final model structure was confirmed using likelihood ratio tests and AICc based model selection. We therefore modelled instantaneous speed as:

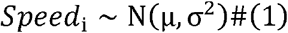

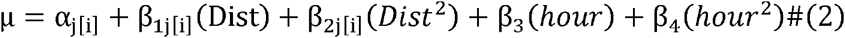

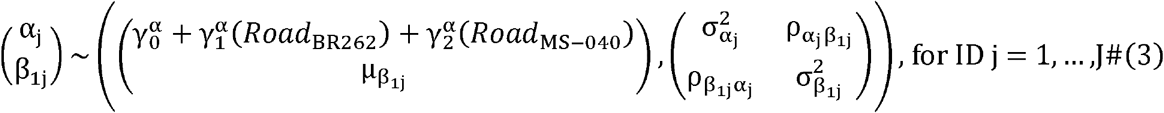

Where Eq. (1) defines the model’s stochastic component, Eq. (2) the models fixed effects structure, and Eq. (3) the model’s random effects structure. For these analyses, all animals that lived >2 km from a paved road were excluded.

##### Q2) Does traffic volume influence giant anteater crossing behaviour?

To answer Q_2_ we modelled the relationship between the number of road crossings according to the nearest highway as a proxy for traffic volume. We also included the variables related to the home range location (distance of home range centroid to nearest paved road), sampling duration, sex, and mass. The road crossing data were zero-inflated, where only 26 of the 38 animals were actually observed to have crossed a road. As a result, we analysed these data using a hurdle model (Zuur et al. 2009) that modelled individual crossing/not crossing information according to a logistic regression model, and the individual number of crossings for those animals that did cross according to a zero-truncated negative binomial generalised linear model. This formulation allowed us to distinguish between the factors governing whether giant anteaters crossed paved roads or not, and the factors driving the crossing rate for those animals that did cross, allowing us to capture both ecological processes in a single model. For both processes, we started with the following global model:

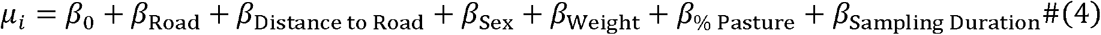

From this global model, we specified a subset of candidate models comprising of all possible combinations of fixed effects using the R package ‘MuMIn’ (Bartoń 2016), and used the AICc for model selection to identify the best-fit model for the data. As AICc has been shown to under/overfit models on small sample sizes (Brewer et al. 2016), we confirmed our selected model via block cross-validation using the methods implemented in the R package ‘DAAG’ (Maindonald et al. 2015). We also tested if the time of day for each road crossing event was related to the hourly traffic volume variation, using simple linear regression.

##### Q3) Do anteaters prefer to cross the roads via passage structures?

To address Q_3_, we identified the crossings on each highway that were within 20m (the median GPS measurement error) of a road passage. These were classified as crossings where a giant anteater could potentially have used that structure to move across the road. This allowed us to search for evidence that anteaters preferred to cross the roads through existing passages.

##### Q4) Do anteaters respond to roads differently than to natural barriers?

Finally, to answer Q_4_, we compared the number of crossings across paved roads to those across unpaved roads and streams using paired t-tests. In a complementary approach, we used simulations to generate a null model of the number of crossings across all linear features that would be expected by chance. For each giant anteater tracked, we used their CTMM to simulate 1000 movement datasets with sampling times that matched the empirical data. We then quantified the number and time of day of road and stream crossings in the simulated movement and averaged the results for each individual. This provided an estimate of the number of road crossings that would be expected by random movement within individuals’ home ranges that we could compare our empirical results against. Observed and expected crossings across paved roads, unpaved roads and streams were compared using paired t-tests

## Results

We deployed collars on 43 individuals out of a total of 45 total captures. These 43 animals were considered in good health and no signs of infection or diseases were observed during clinical exams performed upon captures and recaptures. Importantly, no differences were noted between animals living near or far from the highway. Six anteaters were found dead in the course of the study period, four of which were road-killed and two due to unknown causes. Two of the monitored animals were found dead on the road after the study and their collar was removed. They were identified through their microchip. (Appendix S1). Trackers operated for a median of 11.2 months across all tagged animals. The final GPS dataset comprised 847,683 GPS fixes collected over 12,761 individual◻days (Appendix S1).

### Movement data summary

The 38 range-resident individuals occupied stable home ranges, with an average 95% area of 6.8 km^2^ (4.6 – 9.0 km^2^, 95% CIs reported hereafter). We found that males had significantly larger home range areas than females (8.8 km^2^, 5.9 – 11.6 km^2^ versus 4.4 km^2^, 1.2 – 7.6 km^2^ respectively; *Z* = 2.01, *p* = 0.045). There was no evidence for a relationship between home-range area and body size (*Z* = −1.95, *p* = 0.052), however, we did find a negative relationship between home-range area and the proportion of pastureland that it contained (*Z* = −3.8, *p* < 0.001). We further found partial evidence that high traffic volumes seemed to inhibit some anteaters from establishing territories on both sides of the roads, but small sample sizes and substantial inter-individual variation limited the power of these analyses (Appendix S3).

### Q1) Does the movement behaviour of giant anteaters differ when living near paved roads?

When comparing giant anteaters between the three study sites, there were no differences in home-range sizes (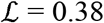; *p* = 0.83; Fig 2a), nor did we find a relationship between the distance that animals lived from a paved road and their home-range sizes (*Z* = 0.15; *p* = 0.88; Fig 2b). There were also no differences between their mean daily speeds (Z = −0.08; *p* = 0.94; Fig 2c) nor any evidence that giant anteaters’ mean daily movement speed differed when living near paved roads with differing traffic volumes (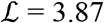; *p* = 0.14; Fig 2d).

**Figure 2.**
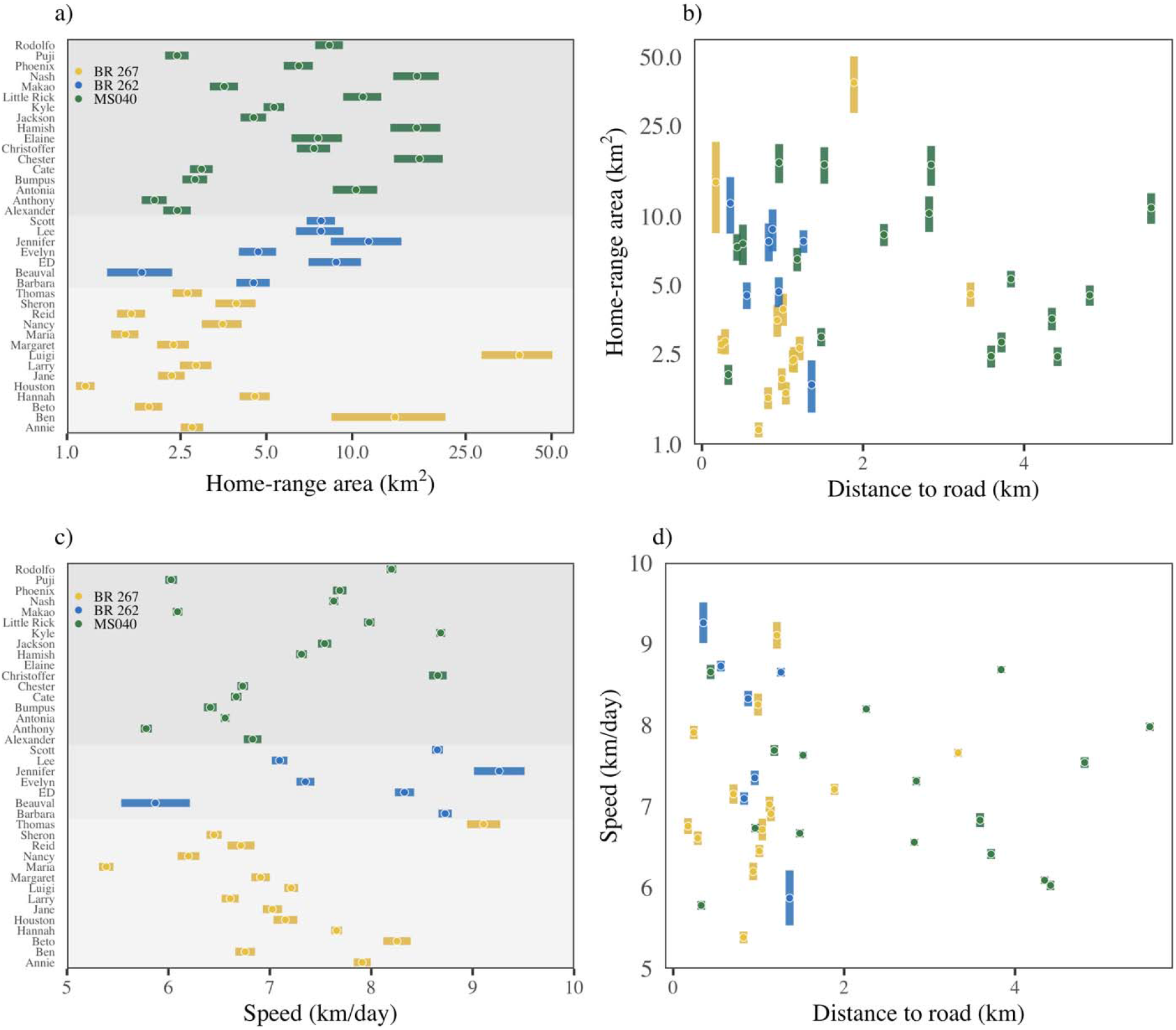
Scatter plots summarizing information for each tracked giant anteater, including the home range area (a), distance of the home range centre to the nearest paved road (b), mean daily movement speed (c) and mean daily movement speed with distance to road (d). Colours and grey shading indicate the three study areas. Circles and bars depict the estimates and the 95% confidence intervals, respectively. Note how there are no clear differences in home-range area or movement speed across sites, nor with distances to roads.

Interestingly, however, we did find evidence that individuals modified their instantaneous movement speed depending on how far they were from roads (Table 2a). This relationship was non-linear and, all else being equal, individuals tended to move slowly when close to or on roads, speed up at intermediate distances, and then slow down again as they moved further away from roads. The interclass correlation was relatively low (14.5%), suggesting this general pattern was fairly consistent across all of the anteaters we collared, but with some amount of inter-individual variation in how movement changed with distance to roads (Appendix S2).

**Table 2.** Summary of models relating A) instantaneous movement speed and B) crossing events with different predictors. **A)** parameter estimates 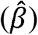, 95% confidence intervals (CI), *t*-values, and *P*-values for the best (<AICc) mixed effects regression model fit to individual instantaneous movement speeds. The model included a first order auto-regressive correlation structure with *ρ* = 0.51. **B)** parameter estimates 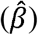, 95% confidence intervals (CI), *Z* values, and *P*-values for the best (<AICc) zero altered negative binomial model fit to the number of times each individual crossed a paved road. Significant predictors (*p*<0.05) are highlighted.

**A**
| | $\hat{\beta}$ | Lower 95% CI | Upper 95% CI | $t$-value | $P$ |
| --- | --- | --- | --- | --- | --- |
| Intercept | 1.01x10<sup>-1</sup> | 9.38 x10<sup>-2</sup> | 1.00 x10<sup>-1</sup> | 28.06 | < 0.001 |
| **Distance to Road** | **1.84x10<sup>-3</sup>** | **3.60 x10<sup>-4</sup>** | **3.31 x10<sup>-3</sup>** | **2.44** | **0.015** |
| **Distance to Road<sup>2</sup>** | **-2.28x10<sup>-4</sup>** | **-3.89 x10<sup>-4</sup>** | **-6.77 x10<sup>-5</sup>** | **-2.79** | **0.005** |
| **Time of day** | **-5.00x10<sup>-3</sup>** | **-5.13 x10<sup>-3</sup>** | **-4.88 x10<sup>-3</sup>** | **-77.06** | **< 0.001** |
| **Time of day<sup>2</sup>** | **2.09x10<sup>-4</sup>** | **2.03 x10<sup>-4</sup>** | **2.14 x10<sup>-4</sup>** | **75.52** | **< 0.001** |
| Road - BR262 | 1.05x10<sup>-2</sup> | -1.27 x10<sup>-3</sup> | 2.23 x10<sup>-2</sup> | 1.83 | 0.078 |
| Road - MS040 | 3.09x10<sup>-3</sup> | -7.78 x10<sup>-3</sup> | 1.40 x10<sup>-2</sup> | 0.58 | 0.564 |

**B**
| | $\hat{\beta}$ | Lower 95% CI | Upper 95% CI | $Z$-value | $P$ |
| --- | --- | --- | --- | --- | --- |
| *Zero hurdle model coefficients (binomial with logit link)* |  |  |  |  |  |
| **Intercept** | **1.97** | **0.06** | **3.87** | **2.03** | **0.043** |
| **Distance to road** | **-1.22** | **-2.32** | **-0.12** | **-2.17** | **0.030** |
| **Sex (male)** | **3.66** | **0.38** | **6.95** | **2.18** | **0.029** |
| Road (BR262) | 17.01 | -11339.11 | 11373.1 | 0.00 | 0.998 |
| Road (MS040) | -1.66 | -4.39 | 1.07 | -1.19 | 0.233 |
| *Count model coefficients (truncated negative binomial with log link)* |  |  |  |  |  |
| **Intercept** | **16.08** | **9.22** | **22.29** | **4.59** | **< 0.001** |
| **Distance to road** | **-2.51** | **-3.55** | **-1.48** | **-4.75** | **< 0.001** |
| **Sex (male)** | **1.33** | **0.26** | **2.40** | **2.44** | **0.015** |
| **Pasture** | **-0.13** | **-0.20** | **-0.06** | **-3.50** | **< 0.001** |

### Q2) Does traffic volume influence giant anteater crossing behaviour?

Overall, we found little evidence that traffic volume influenced giant anteater crossing behaviour. Of the 27 giant anteaters with home range centroids <2km from the nearest paved road (i.e., those individuals with home-ranges bordering or intersecting the nearest paved road), 22 crossed roads, with a median of 2 crossings per animal per day (range 1–15). Across all animals, the selected model (<AICc) predicting whether a giant anteater would cross a road or not (Table 2b) included the distance an animal lived from the nearest paved road, the animal’s sex, and the road it was crossing. Using these three terms alone, we found that a logistic regression model could predict crossings on a hold-out sample with a mean accuracy of 84% (83.8–84.3%, 95% CIs). Unsurprisingly, the closer an animal lived to a road, the more likely it was to cross. Males were ~3.7 times more likely to cross the roads than females, all else being equal. The model further suggests an overall similarity in crossing events across roads.

The selected model section relating the number of crossings per individual for those anteaters that did cross a paved road (Table 2b) included the distance to the nearest paved road, sex, and the amount of pasture contained within their home ranges. Here again, animals that lived closer to paved roads tended to cross them significantly more often, all else being equal (Fig. 3a). However, males tended to cross the roads more frequently than females (Fig. 3b), and the more pastureland that was in an individual’s home range, the less it crossed paved roads (Fig. 3c).

**Figure 3.**
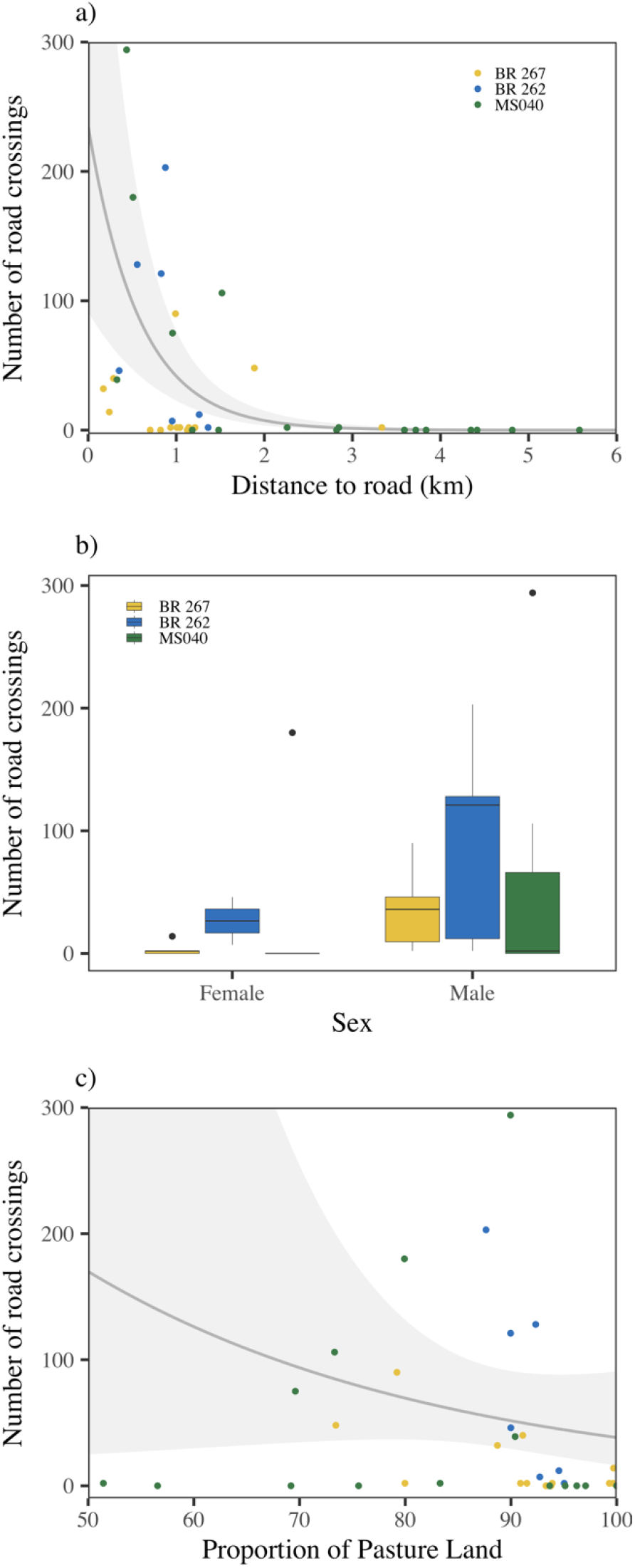
Summary of the crossing results. Panel A) Scatter plots relating the total number of road crossings that each anteater performed in relation to the distance of their home range centroid to the nearest paved road. Panel B) Boxplots depicting the number of road crossings for males and females across the three study sites. In panel c), the relationship between the number of road crossings and the proportion of pastureland in each individual’s home range is shown. In panels a) and c), the grey lines depict the fitted negative-binomial regression model and shaded area the 95% confidence intervals.

Giant anteaters tended to cross paved roads more frequently at night, when the traffic volume was lowest, but this also coincided with when they were more active (Fig. 4). We found a significant negative correlation between mean hourly road crossings and traffic volume (*F*_[1,22]_ = −17.0, *p* < 0.001, adjusted *R*^2^ = 0.41).

**Figure 4.**
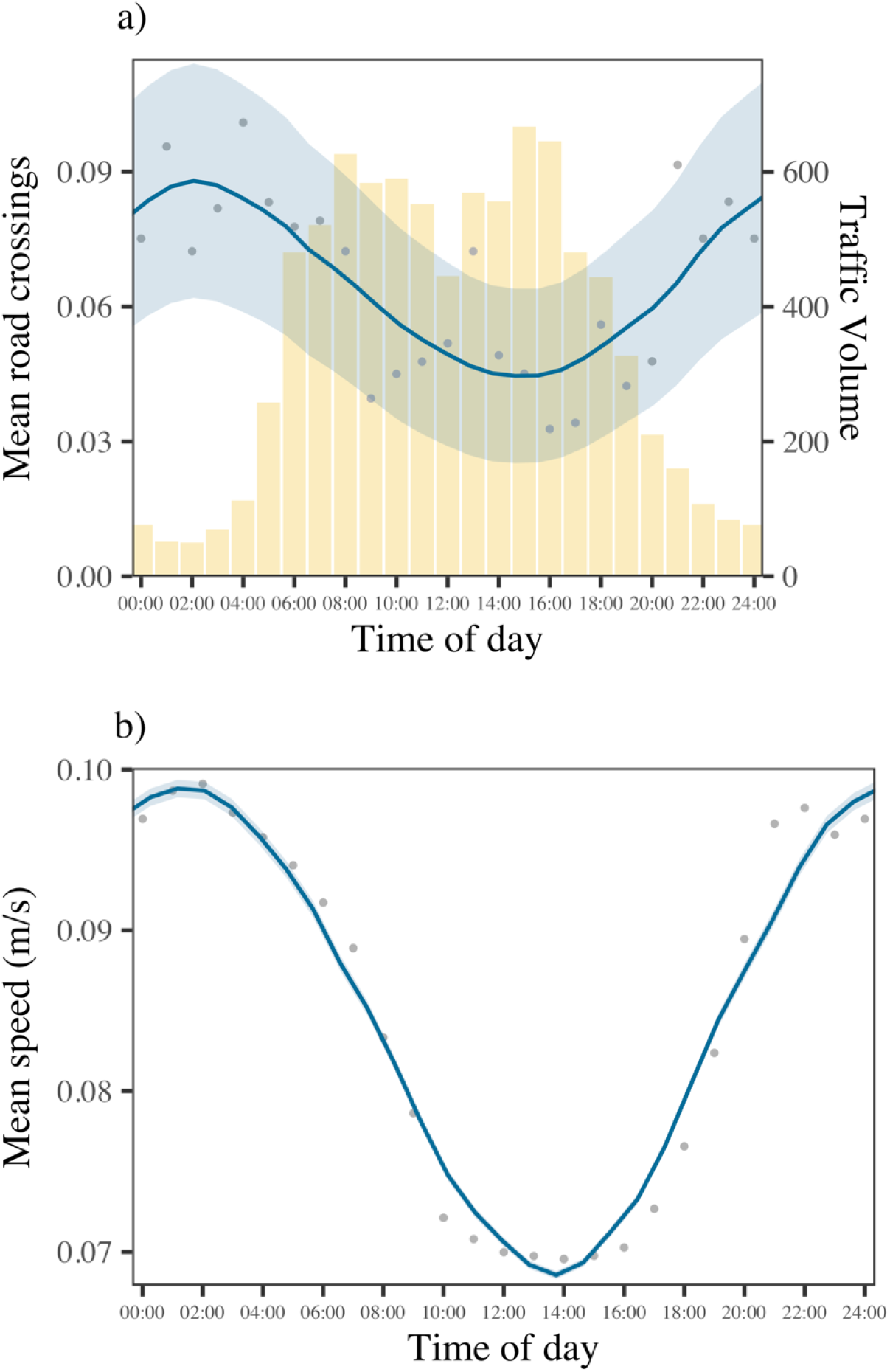
Panel A) Relation between the mean number of crossings per hour (dots, left Y-axis) and the hourly traffic volume in total number of cars (bars, right Y-axis). Panel B) mean speed per time of day over the total duration of the study period. The blue lines depict loess smoothed regression curves fit to the hourly means and the shaded area the 95% confidence intervals on hourly means. Note how most road crossings occurred at night when traffic volume was lowest and when anteaters are more active.

### Q3 Do anteaters prefer to cross the roads via passage structures?

We found no evidence that anteaters searched for road passages (culverts or viaducts) to cross the roads. The median distance of road crossings from the nearest road passage was 1720m (1710 – 1730m, 95% CIs). Only 19 crossing events (1.2%) were within 20m of a passage, thus potentially utilizing it, and all of these were from the same animal living near BR-267.

### Q4 Do anteaters respond to roads differently than to natural barriers?

We found significant differences when comparing the number of crossings of giant anteaters across paved roads to that across unpaved roads and natural linear features. Individuals crossed paved roads significantly fewer times than they crossed both unpaved roads (*t_[37]_* = −3.7, *p* < 0.001) and streams (*t_[37]_* = −3.4, *p* < 0.005; Fig. 5). This is also supported by the movement simulations results, in which the observed number of paved road crossings was ~3.3 times lower than would have been expected by random chance (*t_[37]_*=−3.6, *p* <0.001). In contrast, the observed number of dirt road crossings was only ~2.0 times lower than would have been expected by random chance (*t_[37]_*=4.5, *p* <0.001), while the number of stream crossings was not significantly different than would have been expected by random chance (*t_[37]_*=0.6, *p* =0.57).

**Figure 5.**
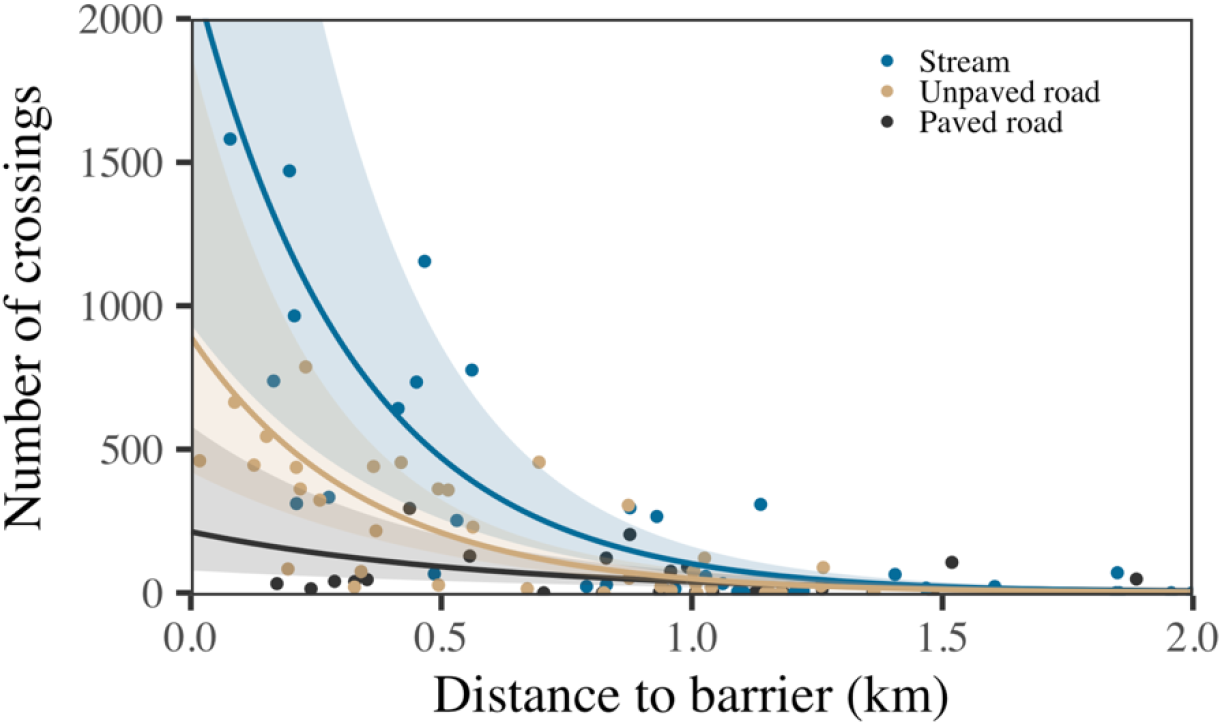
Scatterplot of the relationship between the number of crossings across barriers, and the distance the individual lived from that barrier. The solid lines depict fitted negative-binomial regression models and shaded areas the 95% confidence intervals.

## Discussion

A key step in reducing negative effects of roads on wildlife is to understand how individuals behave on and around roads of varying traffic volumes. From detailed tracking of 38 non-dispersing giant anteaters, we found that while individuals did tend to reduce their movement speed when approaching roads, they otherwise showed few signs of adapting their movement when living near paved roads of varying traffic volumes. High traffic volumes seem to inhibit some anteaters from establishing territories on both sides of the roads, and alongside this we found that individuals crossed paved roads less often than would be expected by random chance, whereas there was no difference between the number of observed stream crossings and what would have been expected by chance. In other words, this inhibition was not due to linear features functioning as home range boundaries (Riley et al. 2006), but rather that traffic may occasionally deter anteaters from crossing roads. This was also reflected in our finding that animals reduced their movement speeds as they approached paved roads, suggesting that some level of caution is taken when in the immediate vicinity of high traffic roads. Nevertheless, none of the highways we monitored were impervious to anteaters, and a high proportion (>80%) of tracked anteaters living near the paved roads (<2km) did cross them regularly. Overall, the traffic, noise, or light pollution associated with roads did not appear to affect their behaviour one way or another. In fact, it is possible to observe giant anteaters foraging at the edge of major highways, and one of the individuals tracked in this study even slept regularly in the native vegetation near the road, but he was eventually victim of a collision (Appendix S1). Worryingly, we found no evidence that anteaters actively sought out road passages for safe crossings. Although we regularly find giant anteater footprints inside road passages (Abra et al. 2020), our results suggest that few individuals actually make regular use of such passages, and those that do probably only do so opportunistically (e.g., when traveling along streams).

Collectively, these results demonstrate that giant anteaters do not change their movement behaviour and space use near paved roads, nor search for safe crossing structures, which is probably the cause of the high roadkill rates (Ascensão et al. 2021). This contradicts previous research suggesting that the barrier to movement and population redistribution was the most important road impact and emphasises the need to obtain sound empirical data on animal movement when planning and managing transportation networks (Ascensão et al. 2019b).

### Roads as Ecological Traps

When species fail to recognize and avoid suboptimal sink habitats, the result can be a decrease in population sizes and an increased extinction risk (Pulliam & Danielson 1991). To this end, no differences in clinical exams, clinical signs, or body scores were noted between animals living near or far from the paved roads, therefore eliminating the alternative hypothesis that territories near roads are suboptimal, and those anteaters that occupy such areas were likely to be more debilitated. Indeed, the best model for predicting how far an anteater lived from a road was the intercept only model (Appendix S2). Our results suggest that a large proportion of road-killed anteaters are roadside residents and not dispersing animal, As six of the animals monitored in this study eventually died from vehicle collisions (~14%; Appendix S2). The three animals that dispersed during this study (distances: 35 km, 50km and 100km), crossed roads but survived and all established themselves. Because roadside habitats apparently offer good foraging opportunities that allow giant anteaters to remain healthy, the feedback that giant anteaters receive about the quality of roadside habitat may be misguiding. This makes it unlikely that they will learn that roadsides are mortality sinks that should be avoided. Hence, habitat near roads may function as an ecological trap where healthy individuals occupy the territories nearby or bisected by roads but eventually are road-killed given their regular crossings, leaving the territory vacant for subsequent occupation.

This dynamic is a worst-case scenario for this already threatened species that is likely to negatively impact the long-term population viability (Miranda et al. 2014). Moreover, giant anteaters live at low population densities and have a low recruitment (Gaudin et al. 2018; Desbiez et al. 2020; Bertassoni et al. *in press*), making them particularly vulnerable to WVC (Miranda et al. 2014). Indeed, population viability analysis showed how giant anteaters deaths due to vehicle collisions decrease the stochastic growth rate of populations by half, making them drastically less resilient to other threats, and slows their recovery time from catastrophic events (Desbiez et al. 2020). The sex ratios of roadkill (Barragán-Ruiz et al. 2021) and our results on road crossing rates are consistent, both suggesting that males are more threatened by roadkill due to their higher crossing rates relatively to females. The fact that males are the most affected by roadkill is less critical than if it were females (Desbiez et al. 2020). This is especially so given that females care for new born pups for at least the first 6-12 months (Jerez & Halloy 2003; Desbiez et al. 2020). Yet, there may be other unforeseen downstream genetic effects resulting from the high mortality of males, which may further imperil population persistence, and further study is clearly warranted.

### Conservation Implications

As the transportation networks in South-Central America continue to expand, the impact of road, and most probably railway (Dasoler et al. 2020), induced mortality is likely going to worsen. This may severely limit population persistence throughout the giant anteater’s distribution, calling for immediate road mitigation actions. Because anteaters are mostly indifferent to roadside habitat, and do not actively seek out road passages, it is unlikely that establishing a network of crossing structures alone will provide an effective solution. Fencing, properly linked to existing passages, has been suggested as a cost-effective mitigation measure, for both conservation purposes as well for human safety, as it allows separating anteaters and other large animals from traffic, thus significantly reducing the likelihood of collisions (Clevenger et al. 2001; Jaeger & Fahrig 2004; Spanowicz et al. 2020; Ascensão et al. 2021). Another complementary approach is to manage traffic speed in some road sections (via e.g., stop signs, speeds bumps, speed limits, and/or animal crossing signs) in order to decrease the likelihood of drivers hitting animals.

The application of such management measures throughout the entire road network is unfeasible. One possible approach is to implement mitigation measures in locations with the highest WVC rates (Ascensão et al. 2021) and in road sections that clearly bisect areas of higher landscape connectivity (Grilo et al. 2011). Another approach would be to prioritize high traffic volume roads. The similarity of crossing events across the three roads suggests that the probability of giant anteaters incurring in collisions is traffic volume dependent. In fact, a systematic roadkill survey in this same study area recorded three times as many cases on high-traffic roads (BR262 and BR267) as on the lower traffic MS040 (Ascensão et al. 2021). Given that we currently lack sound estimates on local anteater abundance in the three study sites, we cannot affirm if such differences in mortality rates were due to different population sizes or different probability of being road-killed. Yet, given the similarity in land cover across the three study areas, it is reasonable to assume that the abundance of giant anteaters across them is similar. As so, the mitigation of high traffic roads should be prioritised. However, given that the anteaters’ activity peaks at night, similarly as other large mammals recurrently involved in WVC, notably tapirs (*Tapirus terrestris*), reducing the overall amount of night-time driving would most likely result in fewer collisions and consequently lower human injuries (Hobday & Hobday 2010).

## Conclusion

Coupling detailed movement information with contemporary data from systematic road surveys allowed us to disentangle the main effects of roads on giant anteaters. Roads did not affect the movement behaviour and space use of giant anteaters, nor were substantial barriers for their displacements. In turn, anteaters did not search for passages for safe crossings and readily occupied the areas near the roads. As a consequence, roadside areas are sink habitats due to the high roadkill rates, which may threaten the population persistence at the local scale. Such information is critical in the development of road and landscape management strategies, including the planning of future transportation infrastructures.

## Supporting Information

Additional information is available online in the Supporting Information section at the end of the online article. The authors are solely responsible for the content and functionality of these materials. Queries (other than absence of the material) should be directed to the corresponding author.

## Acknowledgments

We would like to thank the donors to the Anteaters & Highways Project especially the Foundation Segre as well as North American and European Zoos listed at (http://www.giantanteater.org/). We would also like to thank the owners of all the ranches that allowed us to monitor animals on their property, in particular Nhuveira, Quatro Irmãos and Santa Lourdes ranches. Thank you to M. Alves, D. Kluyber, C. Luba, A. Alves. FA was funded by Fundação para a Ciência e Tecnologia (contract CEECIND/03265/2017). We would like to thank all the volunteers that helped us on carrying out the fieldwork: AC Alves, AC Przvdzimirckí, AC Rodrigues,AG Pardo, AM Gomide, AM Leonardo, AM Silva, AN Figueiredo, B GoldFarb, B Harrower, BG de Oliveira, BLF dos Santos, BPR Leite, C Dimitrius, CA Soares, CBS Polisel, CEB Ruiz, CM Mazeto, CO Pinto, CS Bangsgaard, EN Saito, EO Pacheco, FC Jacoby, FFW de Souza, FMB da Silva, FN Paz, GD de Araújo, GF Massocato, GM Miranda, GT Dalazen, HC Soares, JGS Junior, JHM Dias, KTP Sabino, L Fromme, LC da Silva, LHGR Junior, LM Barreto, LM Gonçalves, LR Rezende, LY de Paulo, M Setti, MA da Silva, MC da Silva, MD Mattos, MEM Nascimento, MFM Paiva, MJG de Souza, MP da Silva, MP Figueira, MR Costa, MY de Paulo, P Petrazzini, PEN Suares, PGV Saracchini, PLP Gonçalves, PP Dutra, R Soares, R Yordi, SA Rodrigues, SS Sierakowski, T Giauque, T Tezlaff, TB Moreira, TSG da Silva, V Alberici, VS Dias, WC de Souza, WCM Diniz, WR Alves, Y Machado, YGG Ribeiro.

## Statement on human or animal subjects

This study was performed under License No. 53798 (Chico Mendes Institute for Biodiversity Conservation) that granted permission to capture, immobilize, and manipulate giant anteaters and collect/store biological samples. All procedures followed the Guidelines of the American Society of Mammologists for the use of wild mammals in research (Sikes et al., 2016).

## Notes

### Competing Interest Statement

The authors have declared no competing interest.

## References

1. Abra FD, Canena A da C, Garbino GST, Medici EP. 2020. Use of unfenced highway underpasses by lowland tapirs and other medium and large mammals in central-western Brazil. Perspectives in Ecology and Conservation. Available from http://www.sciencedirect.com/science/article/pii/S2530064420300651 (accessed November 17, 2020).

2. Abra FD, Granziera BM, Huijser MP, Ferraz KMPM de B, Haddad CM, Paolino RM. 2019. Pay or prevent? Human safety, costs to society and legal perspectives on animal-vehicle collisions in São Paulo state, Brazil. PLoS ONE 14. Available from https://www.ncbi.nlm.nih.gov/pmc/articles/PMC6459512/ (accessed May 22, 2020).

3. Ascensão F, Clevenger AP, Grilo C, Filipe J, Santos-Reis M. 2012. Highway verges as habitat providers for small mammals in agrosilvopastoral environments. Biodiversity and Conservation 21:3681–3697.

4. Ascensão F, D’Amico M, Barrientos R. 2019a. Validation data is needed to support modelling in Road Ecology. Biological Conservation 230:199–200. Elsevier BV.

5. Ascensão F, Lucas PS, Costa A, Bager A. 2017. The effect of roads on edge permeability and movement patterns for small mammals: a case study with Montane Akodont. Landscape Ecology 32:781–790.

6. Ascensão F, Yogui D, Alves M, Medici EP, Desbiez A. 2019b. Predicting spatiotemporal patterns of road mortality for medium-large mammals. Journal of Environmental Management 248:109320.

7. Ascensão F, Yogui DR, Alves MH, Alves AC, Abra F, Desbiez ALJ. 2021. Preventing wildlife roadkill can offset mitigation investments in short-medium term. Biological Conservation 253:108902.

8. Barragán-Ruiz CE, Paviotti-Fischer E, Rodríguez-Castro KG, Desbiez ALJ, Galetti PM. 2021. Molecular sexing of Xenarthra: a tool for genetic and ecological studies. Conservation Genetics Resources 13:41–45.

9. Barrientos R, Bolonio L. 2009. The presence of rabbits adjacent to roads increases polecat road mortality. Biodiversity and Conservation 18:405–418.

10. Bartoń K. 2016. MuMIn: Multi-Model Inference. R package version 1.15.6.

11. Bertassoni A, Bianchi R, Desbiez A. (in press). Camera trap individual identification of giant anteaters to estimate population size and viability. Journal of Wildlife Management and Wildlife Monographs.

12. Bertassoni A, Mourão G, Ribeiro RC, Cesário CS, Oliveira JP de, Bianchi R de C. 2017. Movement patterns and space use of the first giant anteater (Myrmecophaga tridactyla) monitored in São Paulo State, Brazil. Studies on Neotropical Fauna and Environment 52:68–74. Taylor & Francis.

13. Brewer MJ, Butler A, Cooksley SL. 2016. The relative performance of AIC, AICC and BIC in the presence of unobserved heterogeneity. Methods in Ecology and Evolution 7:679–692. Wiley Online Library.

14. Buchholtz EK, Stronza A, Songhurst A, McCulloch G, Fitzgerald LA. 2020. Using landscape connectivity to predict human-wildlife conflict. Biological Conservation 248:108677.

15. Calabrese JM, Fleming CH, Gurarie E. 2016. ctmm: an r package for analyzing animal relocation data as a continuous-time stochastic process. Methods in Ecology and Evolution 7:1124–1132.

16. Camilo◻Alves C de S e P, Mourão G de M. 2006. Responses of a Specialized Insectivorous Mammal (Myrmecophaga tridactyla) to Variation in Ambient Temperature 1. Biotropica: The Journal of Biology and Conservation 38:52–56. Wiley Online Library.

17. Ceia-Hasse A, Navarro LM, Borda-de-Água L, Pereira HM. 2018. Population persistence in landscapes fragmented by roads: Disentangling isolation, mortality, and the effect of dispersal. Ecological Modelling 375:45–53.

18. Clark RW, Brown WS, Stechert R, Zamudio KR. 2010. Roads, interrupted dispersal, and genetic diversity in timber rattlesnakes. Conservation Biology 24:1059–1069. Wiley Online Library.

19. Clevenger AP, Chruszcz B, Gunson KE. 2001. Highway mitigation fencing reduces wildlife-vehicle collisions. Wildlife Society Bulletin:646–653.

20. D’Amico M, Périquet S, Román J, Revilla E. 2016. Road avoidance responses determine the impact of heterogeneous road networks at a regional scale. Journal of Applied Ecology 53:181–190.

21. D’Amico M, Román J, de los Reyes L, Revilla E. 2015. Vertebrate road-kill patterns in Mediterranean habitats: Who, when and where. Biological Conservation 191:234–242.

22. Dasoler BT, Kindel A, Beduschi J, Biasotto LD, Dornas RAP, Gonçalves LO, Lombardi PM, Menger T, de Oliveira GS, Teixeira FZ. 2020. The need to consider searcher efficiency and carcass persistence in railway wildlife fatality studies. European Journal of Wildlife Research 66:81.

23. De Magalhaes JP, Costa J. 2009. A database of vertebrate longevity records and their relation to other life◻history traits. Journal of evolutionary biology 22:1770–1774. Wiley Online Library.

24. Desbiez ALJ, Bertassoni A, Traylor-Holzer K. 2020. Population viability analysis as a tool for giant anteater conservation. Perspectives in Ecology and Conservation. Available from http://www.sciencedirect.com/science/article/pii/S2530064420300213 (accessed June 16, 2020).

25. DNIT. 2020. PNCT | DNIT. Available from http://servicos.dnit.gov.br/dadospnct (accessed June 8, 2020).

26. Fahrig L. 2007. Non-optimal animal movement in human-altered landscapes. Functional Ecology 21:1003–1015.

27. Fahrig L, Rytwinski T. 2009. Effects of roads on animal abundance: an empirical review and synthesis. Ecology and society 14. JSTOR.

28. Fleming C, Calabrese JM. 2020. ctmm: Continuous-Time Movement Modeling. Available from https://CRAN.R-project.org/package=ctmm.

29. Fleming C, Fagan WF, Mueller T, Olson KA, Leimgruber P, Calabrese JM. 2015. Rigorous home range estimation with movement data: a new autocorrelated kernel density estimator. Ecology 96:1182–1188.

30. Fleming CH, Calabrese JM. 2017. A new kernel density estimator for accurate home◻range and speciesLrange area estimation. Methods in Ecology and Evolution 8:571–579. Wiley Online Library.

31. Fleming CH, Drescher-Lehman J, Noonan MJ, Akre TSB, Brown DJ, Cochrane MM, Dejid N, DeNicola V, DePerno CS, Dunlop JN. 2020. A comprehensive framework for handling location error in animal tracking data. bioRxiv. Cold Spring Harbor Laboratory.

32. Fleming CH, Noonan MJ, Medici EP, Calabrese JM. 2019. Overcoming the challenge of small effective sample sizes in home◻range estimation. Methods in Ecology and Evolution 10:1679–1689. Wiley Online Library.

33. Forman R, Sperling D, Bissonette JA, Clevenger AP, Cutshall CD, Dale VH, Fahrig L, France RL, Heanue K, Goldman CR. 2003. Road ecology: science and solutions. Island Press.

34. Gardner AL. 2008. Mammals of South America, volume 1: marsupials, xenarthrans, shrews, and bats. University of Chicago Press.

35. Gaudin TJ, Hicks P, Di Blanco Y. 2018. Myrmecophaga tridactyla (Pilosa: Myrmecophagidae). Mammalian Species 50:1–13.

36. González-Suárez M, Zanchetta Ferreira F, Grilo C. 2018. Spatial and species-level predictions of road mortality risk using trait data. Global Ecology and Biogeography. Available from http://doi.wiley.com/10.1111/geb.12769 (accessed August 21, 2018).

37. Grilo C, Ascensão F, Santos-Reis M, Bissonette JA. 2011. Do well-connected landscapes promote road-related mortality? European Journal of Wildlife Research 57:707–716.

38. Hobday AJ, Hobday AJ. 2010. Nighttime driver detection distances for Tasmanian fauna: informing speed limits to reduce roadkill. Wildlife Research 37:265–272. CSIRO PUBLISHING.

39. Jaeger JA, Fahrig L. 2004. Effects of road fencing on population persistence. Conservation Biology 18:1651–1657. Wiley Online Library.

40. Jerez S, Halloy M. 2003. El oso hormiguero, Myrmecophaga tridactyla: crecimiento e independización de una cría. Mastozoología Neotropical 10:323–330.

41. Kluyber D, Lopez RPG, Massocato G, Attias N, Desbiez ALJ. 2020. Anesthesia and surgery protocols for intra-abdominal transmitter placement in four species of wild armadillo. Journal of Zoo and Wildlife Medicine 51:514–526. BioOne.

42. Long ES, Diefenbach DR, Wallingford BD, Rosenberry CS. 2010. Influence of roads, rivers, and mountains on natal dispersal of white◻tailed deer. The Journal of Wildlife Management 74:1242–1249. Wiley Online Library.

43. Macdonald DW. 2016. Animal behaviour and its role in carnivore conservation: examples of seven deadly threats. Animal behaviour 120:197–209. Elsevier.

44. Maindonald JH, Braun WJ, Braun MWJ. 2015. Package ‘DAAG.’ Data Analysis and Graphics Data and Functions.

45. McNab BK. 1984. Physiological convergence amongst ant◻eating and termite◻eating mammals. Journal of Zoology 203:485–510. Wiley Online Library.

46. Medri ÍM, Mourão G. 2005. Home range of giant anteaters (Myrmecophaga tridactyla) in the Pantanal wetland, Brazil. Journal of Zoology 266:365–375.

47. Miranda F, Bertassoni A, Abba A. 2014. Myrmecophaga tridactyla. Available from http://dx.doi.org/10.2305/IUCN.UK.2014-1.RLTS.T14224A47441961.en (accessed February 1, 2019).

48. Montgomery RA, Macdonald DW, Hayward MW. 2020. The inducible defences of large mammals to human lethality. Functional Ecology 34:2426–2441.

49. Noonan MJ, Fleming CH, Akre TS, Drescher-lehman J, Gurarie E, Kays R, Calabrese JM. 2019. Scale-insensitive estimation of speed and distance traveled from animal tracking data. Movement Ecology 7:1–15. Movement Ecology.

50. Pulliam HR, Danielson BJ. 1991. Sources, Sinks, and Habitat Selection: A Landscape Perspective on Population Dynamics. The American Naturalist 137:S50–S66. [University of Chicago Press, American Society of Naturalists].

51. Riley SP, Pollinger JP, Sauvajot RM, York EC, Bromley C, Fuller TK, Wayne RK. 2006. FAST◻TRACK: A southern California freeway is a physical and social barrier to gene flow in carnivores. Molecular ecology 15:1733–1741. Wiley Online Library.

52. Schlaepfer MA, Runge MC, Sherman PW. 2002. Ecological and evolutionary traps. Trends in ecology & evolution 17:474–480. Elsevier.

53. Sih A, Ferrari MCO, Harris DJ. 2011. Evolution and behavioural responses to human-induced rapid environmental change. Evolutionary Applications 4:367–387.

54. Sikes RS, Animal Care and Use Committee of the American Society of Mammalogists. 2016. 2016 Guidelines of the American Society of Mammalogists for the use of wild mammals in research and education. Journal of mammalogy 97:663–688. Oxford University Press US.

55. Souza C, Azevedo T. 2017. MapBiomas general handbook. MapBiomas: São Paulo, Brazil:1–23.

56. Spanowicz AG, Teixeira FZ, Jaeger JAG. 2020. An adaptive plan for prioritizing road sections for fencing to reduce animal mortality. Conservation Biology **n/a**. Available from https://conbio.onlinelibrary.wiley.com/doi/abs/10.1111/cobi.13502 (accessed April 22, 2020).

57. Strassburg BBN et al. 2017. Moment of truth for the Cerrado hotspot. Nature Ecology & Evolution 1:0099.

58. Van der Ree R, Smith DJ, Grilo C. 2015. Handbook of road ecology. John Wiley & Sons.

59. Venter O et al. 2016. Sixteen years of change in the global terrestrial human footprint and implications for biodiversity conservation. Nature Communications 7:12558. Nature Publishing Group.

60. Viechtbauer W. 2010. Conducting meta-analyses in R with the metafor package. Journal of statistical software 36:1–48. UCLA Statistics.

61. Zuur A, Ieno EN, Walker N, Saveliev AA, Smith GM. 2009. Mixed effects models and extensions in ecology with R. Springer Science & Business Media.

