## Appendix S1 for "Roads as ecological traps for giant anteaters": Appendix_S1.html

Appendix S1 - Workflow used to analyse the giant anteater GPS data


### Appendix S1 - Workflow used to analyse the giant anteater GPS data

This document was created on March 30, 2021.

---

Our primary aim was to understand whether giant anteater (*Myrmecophaga tridactyla*) movement was constrained by roads and the relationship between movement behaviour and roadkill. In particular, we aimed to address three over-arching questions:

- Does the movement behaviour of giant anteaters differ when living near paved roads?
- Does traffic volume influence giant anteater crossing behaviour?
- Do anteaters prefer to cross the roads via passage structures?
- Do anteaters respond to roads differently than to natural barriers?

To do this, we used GPS data to quantify key movement metrics using the methods implemented in the `R` package `ctmm`. In this appendix we detail the workflow that was used on a data from a single giant anteater living near the highway ‘BR 262’. This process was repeated across all individuals and results were compiled and analysed as described in the main text, and following the workflow described in Appendix S2.

---

#### Figures depicting field protocols

In this section we include to figures depicting the giant anteater handling, and GPS harness deployment.

**Figure S1.1** Figure depicting the handling procedure in the field.

**Figure S1.2** Figure showing a giant anteater wearing one of the GPS harnesses shortly after release.

---

#### Data Import and pre-processing

Before analysis, we performed a data cleaning process in order to calibrate the GPS error and filter unreliable locations.

##### Data Import

The first step of the analysis is to import the data and convert to a telemetry object for use in `ctmm`. Here the final dataset has a column indicating which locations were considered outliers.

```
#Load in the necessary packages
library(ctmm)
library(rgeos)
library(ggplot2)

#Import the GPS dataset (Gets imported as an R objected called data)
load("Anteater_GPS_Data.Rda")

#Convert the dataset with outliers to a telemetry object for demonstration purposes
DATA_RAW <- as.telemetry(data)

#Drop the outliers
data_clean <- data[which(data$OUT == 0),]

#Convert the clean dataset to a telemetry object
DATA <- as.telemetry(data_clean)

#Summary of full dataset
summary(DATA)
```

```
##                 interval (min) period (mon) longitude  latitude
## Alexander             20.00000    11.921256 -53.75980 -21.14109
## Annie                 20.00000    11.096325 -53.49576 -21.63346
## Anthony               20.00000    14.080832 -53.75685 -21.11174
## Antonia               20.00000    11.844478 -53.91083 -21.04103
## Barbara               20.00000     7.439086 -54.03537 -20.46692
## Beauval               20.00000     4.812804 -54.09208 -20.48623
## Ben                   20.00000     2.609817 -53.46053 -21.65261
## Beto                  20.00000    12.045447 -53.50511 -21.63142
## Bumpus                20.00000    13.232697 -53.75040 -21.14285
## Cate                  20.00000    12.362887 -53.75986 -21.12229
## Chester               20.00000    13.304471 -53.93155 -21.04569
## Christoffer           20.00000     9.007152 -53.75909 -21.10599
## Delphine              20.00000    10.361044 -53.69504 -20.78791
## ED                    20.00000     6.931137 -54.14330 -20.47415
## Elaine                20.00000    13.136653 -53.75061 -21.11452
## Evelyn                20.00000    10.997062 -54.11851 -20.46768
## Gala                  20.00000     8.930011 -53.52189 -21.63222
## Hamish                20.00000    11.824547 -53.94130 -21.01902
## Hannah                20.00000    11.245092 -53.59645 -21.66039
## Houston               20.00000    10.924187 -53.50196 -21.62747
## Jackson               20.00000    14.463984 -53.74615 -21.15241
## Jane                  20.00000    11.162642 -53.48273 -21.62628
## Jennifer 262          20.01667     3.715867 -53.96972 -20.42351
## Kyle                  20.00000    14.387329 -53.75067 -21.14322
## Larry 267             20.00000     6.830496 -53.49571 -21.63331
## Lee                   20.00000    10.835320 -53.93331 -20.42216
## Little Rick           20.00000    10.232655 -53.75634 -21.15988
## Luigi                 20.00000    11.888605 -53.58243 -21.64823
## Makao                 20.00000    12.184317 -53.75183 -21.14762
## Margaret              20.00000     8.828889 -53.50742 -21.62868
## Maria                 20.00000    14.664479 -53.60082 -21.63678
## Nancy 05              20.00000     5.981558 -53.59249 -21.63860
## Nash                  20.00000    13.268255 -53.99281 -21.03864
## Phoenix 1 error       20.00000     1.562896 -53.97351 -21.02186
## Phoenix 2             20.00000    13.945722 -53.97210 -21.02095
## Puji                  20.00000    12.045415 -53.75366 -21.14839
## Reid                  20.00000    11.674806 -53.50453 -21.63001
## Rodolfo               20.00000    13.414102 -53.74969 -21.13223
## Scott                 20.00000    10.869616 -54.10078 -20.48530
## Segre                 20.00000     4.562833 -53.92258 -20.97368
## Sheron                20.00000     6.436336 -53.60102 -21.63810
## Thomas                20.00000     5.887029 -53.50656 -21.62690
## Yoki                  20.00000     5.527696 -54.54988 -20.48970
```

The individual named `Phoenix 1 error` is the same animal as `Phoenix 2`. There was a collar malfunction on the first deployment for this animal and the collar needed replacing. Only the locations from the second deployment were used. In addition, the individual named `Yoki` represents an collar that stayed in a fixed location and was only used to calibrate the measurement error (detailed below). Finally, the individuals named `Delphine`, `Gala`, and `Segre` dispersed and did not maintain stable home ranges, so these were excluded from our analyses.

##### Figure of Sampling Times

In this subsection we generate histograms of the amount of data collected over time for each of the 38 individuals included in the final analyses.

```
#Drop the three dispersers and collar malfunctions
data_clean <- data_clean[which(data_clean$ID != "Delphine"),]
data_clean <- data_clean[which(data_clean$ID != "Gala"),]
data_clean <- data_clean[which(data_clean$ID != "Segre"),]
data_clean <- data_clean[which(data_clean$ID != "Phoenix 1 error"),]
data_clean <- data_clean[which(data_clean$ID != "Yoki"),]

data_clean$timestamp <- as.POSIXct(data_clean$timestamp)

#Re-order based on location
data_clean <- data_clean[order(data_clean$Road),]

#Set up a labeller for the facet labels
data_clean$ID2 <- as.factor(paste(data_clean$Road, data_clean$ID))
anteater_names <- as.list(unique(data_clean$ID))
names(anteater_names) <- unique(data_clean$ID2)
name_labeller <- function(variable,value){
  return(anteater_names[value])
}

# Create Figure S1
ggplot(data = data_clean,
       aes(x = timestamp,
           fill = Road)) +
  geom_histogram(bins = 60) +
  scale_fill_manual(labels = c("BR 267",
                               "BR 262",
                               "MS040"), values = c("#e6c141",
                                                    "#3471bc",
                                                    "#3c7a47")) +
  facet_grid(ID2 ~ .,
             labeller = name_labeller) +
  theme_bw() + 
  theme(panel.grid.major = element_blank(),
        panel.grid.minor = element_blank(),
        axis.ticks.y = element_blank(),
        axis.title.y = element_text(size=10, family = "serif"),
        axis.title.x = element_blank(),
        axis.text.y = element_text(size=4, family = "serif"),
        axis.text.x = element_text(size=8, family = "serif"),
        strip.text.y = element_text(size = 8, family = "serif", angle = 0),
        legend.title=element_blank(),
        legend.text=element_text(size=8, family = "serif"),
        legend.background=element_blank(), 
        legend.key = element_blank(),
        legend.text.align = 0)
```

##### Error calibration

For each location estimate, the GPS trackers recorded a unitless Horizontal Dilution of Precision (HDOP) value, which is a measure of the accuracy of each positional fix. To prepare the data for error-informed analyses, we converted the HDOP values into calibrated error circles by estimating an equivalent range error from 6948 calibration data points where a tag had been left in a fixed location (Fleming et al. 2020).

```
#Visualisation of the calibration data
plot(DATA[[43]], error = FALSE)
```

```
#Estimating the UERE
UERE <- uere.fit(DATA[[43]])

UERE
```

```
## An object of class "UERE"
## [[1]]
##                   horizontal vertical     speed
## QFP [NA-speed]     12.779960 16.44517        NA
## Succeeded [speed]   9.668767 19.98038 0.6067519
## 
## [[2]]
##                   horizontal vertical speed
## QFP [NA-speed]       5312.46 5312.231     0
## Succeeded [speed]    1634.93 1634.985  1635
## 
## [[3]]
## horizontal   vertical      speed 
## 120131.874  69936.842   9350.071 
## 
## [[4]]
## horizontal   vertical      speed 
##   2.112761   1.587093   6.085857 
## 
## [[5]]
## horizontal   vertical      speed 
##  13.126280   6.820447  59.476033 
## 
## [[6]]
## horizontal   vertical      speed 
##       6948       6948       1570 
## 
## Slot "info":
## list()
```

```
#Applying the UERE to the two datasets
uere(DATA) <-UERE
uere(DATA_RAW) <-UERE
```

##### Outlier filtering

Before fitting any movement models to these data, the first step is to ensure that there are no outliers that could bias parameter estimation. For each individual dataset, we filtered out outliers based on error-informed distance from the median longitude and latitude, and the minimum speed required to explain each location’s displacement. This was done using the `outlie()` function in `ctmm`. This function calculates distances from the median longitude and latitude, and maximum speeds over each time step. It returns a `data.frame` of these estimates, as well as a plot with intervals of high speed highlighted with blue segments, and distant locations highlighted with red points. Both estimates account for telemetry error and the speed estimates account for timestamp truncation and assign each time step’s speed to the most likely offending time (based on the speed estimate of adjacent locations). The full data cleaning process is iterative. The worst locations are removed first, and then the diagnostic tools recalculated and the process is repeated until no clear outliers remain in the data. This iterative process is not shown here, but the comparison between the raw and filtered data is shown.

```
#Extract an example individual from the raw dataset
ANTEATER_RAW <- DATA_RAW[[23]]

#Checking for outliers
par(mfrow=c(1,3))

#Visualisation of the individual tracking data
plot(ANTEATER_RAW)

#Plot a check for outliers
OUTLIERS_RAW <- outlie(ANTEATER_RAW, family="serif")
plot(OUTLIERS_RAW, family="serif")
```

```
#Extract an example individual from the cleaned dataset
ANTEATER <- DATA[[23]]

#Visualisation of the individual tracking data
plot(ANTEATER)

#Plot a check for outliers
OUTLIERS_CLEAN <- outlie(ANTEATER, family="serif")
plot(OUTLIERS_CLEAN, family="serif")
```

Note how there is no longer any indication of clear outliers in the filtered data.

---

#### Fitting the movement models

The next step after importing and preparing the data was to fit the continuous-time movement models. Following the workflow described in Calabrese et al. (2016), we fit a series of continuous-time movement models to the data using perturbative-Hybrid Residual Maximum Likelihood (pHREML; (Fleming et al. 2019), and identified the best model for each individual via small-sample-sized corrected Akaike’s Information Criterion (AICc). From each CTMM, we extracted the positional autocorrelation timescale (\(\tau\_p\), in hours), which provides a measure of the home-range crossing time, and the velocity autocorrelation timescale (\(\tau\_v\), in minutes), which provides a measure of directional persistence.

```
#Generate the variogram of the cleaned data
vg.anteater <- variogram(ANTEATER)

#fit the variogram
GUESS <- variogram.fit(vg.anteater, interactive= FALSE)

#turn error on in the model
GUESS$error <- TRUE

#Fit the model
#FIT <- ctmm.select(ANTEATER,
#                   CTMM = GUESS,
#                   method = "pHREML",
#                   control=list(method="pNewton",
#                                cores = -1))

load("~/Dropbox (Personal)/UBC/Side_Projects/Arnaud_Anteaters/Scripts/Results/Fits/Fits_Jennifer 262.rda")

#Plot the variogram of movement data and the fitted model as a visual check
par(mfrow=c(1,2))
plot(vg.anteater,
     CTMM=FIT,
     family="serif",
     fraction = 0.005) 
plot(vg.anteater,
     CTMM=FIT,
     family="serif")
```

```
#Return a summary of the fitted model
summary(FIT)
```

```
## $name
## [1] "OUF anisotropic"
## 
## $DOF
##       mean       area      speed 
##   26.06152   47.75424 3148.78534 
## 
## $CI
##                                low       est      high
## area (square kilometers)  9.868991 13.396227 17.454057
## τ[position] (days)        1.559534  2.183595  3.057377
## τ[velocity] (minutes)     7.525304  7.993553  8.490937
## speed (kilometers/day)   10.717096 10.907606 11.098070
```

Here we see that for this animal the home range crossing time, \(\tau\_p\), was 2.18 days (95% CI: 1.56 - 3.06), whereas velocity autocorrelation timescale, \(\tau\_v\), was 7.99 minutes (95% CI: 7.53 - 8.49).

**Note**: The speed value returned here as part of the summary of the movement model is the mean speed and root mean square (RMS) speed of a stationary Gaussian stochastic process. The Gaussian RMS speed can be calculated very quickly from the fitted movement model’s parameter estimates, but is not proportional to the distance traveled. The approach detailed below represents the mean speed, which is conditioned off of the data and the fitted model, and is generally more accurate.

---

#### Estimating movement speeds

We estimated the mean daily movement speed (in km/day) using continuous-time speed and distance (CTSD) estimation (Noonan et al. 2019b). CTSD uses a simulation-based approach to sample from the distribution of possible trajectories that are consistent with the data and a fitted continuous-time movement model, from which the mean speed estimate and confidence intervals can be extracted. This approach is insensitive to the sampling schedule, enabling robust comparisons across individuals. To quantify individuals’ circadian rhythms, we also estimated from the CTMM the instantaneous movement speed (in m/s), along with 95% confidence interval, for each sampled location.

```
#Estimate the mean speed
SPEED <- speed(ANTEATER, FIT, cores = -1)
```

```
## 
  |                                                                            
  |                                                                      |   0%
  |                                                                            
  |========================                                              |  35%
  |                                                                            
  |=================================================                     |  70%
  |                                                                            
  |======================================================================| 100%
```

```
SPEED
```

```
##                             low      est     high
## speed (kilometers/day) 9.058979 9.218675 9.379056
```

```
#Estimate the instantaneous speeds
SPEEDS <- speeds(ANTEATER, FIT, cores = -1)

head(SPEEDS)
```

```
##          low        est      high          t           timestamp
## 1 0.01709822 0.09705145 0.2112222 1535311203 2018-08-26 19:20:03
## 2 0.01610182 0.09025291 0.1958756 1535312406 2018-08-26 19:40:06
## 3 0.01485900 0.08450199 0.1839874 1535313603 2018-08-26 20:00:03
## 4 0.01499575 0.08526497 0.1856415 1535314806 2018-08-26 20:20:06
## 5 0.01623196 0.09219791 0.2006897 1535316036 2018-08-26 20:40:36
## 6 0.01666706 0.09418263 0.2047749 1535318403 2018-08-26 21:20:03
```

Here we see that for this animal the estimated mean daily movement speed was 9.22 km/day (95% CI: 9.06 - 9.38). The instantaneous speeds are estimated for each time a GPS datapoint was collected, and conditional on the fitted movement model (units are in m/s).

---

#### Estimating home range areas

With a fitted movement model in hand, we estimated the 95% home-range areas of each anteater using Autocorrelated Kernel Density Estimation (AKDE; (Fleming et al. 2015). AKDE home-range estimates were conditioned on the autocorrelation structure of the best fit model identified above, and we implemented the small-sample-size bias correction of (Fleming and Calabrese 2017).

```
#calculate the AKDE based on the best fit model
anteater.akde <- akde(ANTEATER, FIT)

#Return the basic statistics on the HR area
summary(anteater.akde)
```

```
## $DOF
##      area bandwidth 
##  47.75424 107.76285 
## 
## $CI
##                               low     est     high
## area (square kilometers) 8.410823 11.4169 14.87518
```

```
#Create a function that scales colours between red and blue
rbPal <- colorRampPalette(c('#FF0000','#046C9A'))
#Then create a variable that scales from red to blue between the two times
ANTEATER$Col <- rbPal(nrow(ANTEATER))[as.numeric(cut(ANTEATER$t,breaks = nrow(ANTEATER)))]

#Plot the AKDE range estimate, with the relocation data, coloured by time
plot(ANTEATER,
     UD=anteater.akde,
     col.grid = NA,
     family = "serif",
     pch = 20,
     cex = 0.2,
     col.DF = "#669543",
     col = ANTEATER$Col,
     labels=FALSE)
```

Here we see that for this animal the estimated 95% home range area was 11.42 km\(^2\) (95% CI: 8.41 - 14.88). Visual diagnostics show a good correspondence between the movement data and the estimated home range area. The locations are coloured by time and there is no evidence of any range shift during the study period, confirming the asymptotic behaviour of the variogram shown above.

##### Percent Land Cover in Home Range Areas

```
#Import the land cover information
mapbiomas <- raster("~/Dropbox (Smithsonian)/Anteater Movement paper/GIS/mapbiomas-matogrossodosul-2018.tif")

#Convert the HR contours to SP objects and reproject
HR_contour <- SpatialPolygonsDataFrame.UD(anteater.akde, level = 0)
HR_contour <- spTransform(HR_contour,projection(mapbiomas))

#Extract land cover information from the raster
myLC <- raster::extract(mapbiomas, HR_contour)

class.prop <- as.data.frame(lapply(myLC, FUN = function(x) { prop.table(table(x)) * 100}))

#Rename the classes so that they are interpretable
names(class.prop)[3] <- "Class"
class.prop$LC_rec <- as.character(class.prop$Class)
class.prop$LC_rec[class.prop$Class %in% c("3", "4")]="Native forest"
class.prop$LC_rec[class.prop$Class %in% c("9")]="Forestry"
class.prop$LC_rec[class.prop$Class %in% c("12","15")]="Pasture"
class.prop$LC_rec[class.prop$Class %in% c("19")]="Agriculture"
class.prop$LC_rec[class.prop$Class %in% c("25")]="Others"
class.prop$LC_rec[class.prop$Class %in% c("33")]="Water"

#Land cover information
Land_Cover <- aggregate(Freq.2 ~ LC_rec,
                        data = class.prop,
                        FUN = "sum")
Land_Cover
```

```
##          LC_rec     Freq.2
## 1      Forestry  1.1747322
## 2 Native forest  8.1861840
## 3       Pasture 90.0184706
## 4         Water  0.6206132
```

From these analyses we can see that 90% of this animal’s home range contains pasture, while 8.2% is comprised of native forest, and the remaining 1.8% belongs to the other land cover types.

###### Percent land cover across all animals

Here, the percent native forest and pasture land in each individual’s home range area is shown.

---

#### Road Interaction Metrics

##### Road permeability

To understand what factors influenced the extent to which giant anteaters were willing to establish their home ranges on both sides of a road according to its traffic volume (Q2), we estimated the ratio of home range areas that fell on either side of the highways across the different paved roads. This was done as follows.

```
library(rgdal)

# Load the road data
BR262 <- readOGR(dsn = "~/Dropbox (Personal)/UBC/Side_Projects/Arnaud_Anteaters/Scripts/Roads/BR262")
```

```
## OGR data source with driver: ESRI Shapefile 
## Source: "/Users/michaelnoonan/Dropbox (Personal)/UBC/Side_Projects/Arnaud_Anteaters/Scripts/Roads/BR262", layer: "BR262"
## with 1 features
## It has 11 fields
## Integer64 fields read as strings:  tessellate extrude visibility drawOrder
```

```
#Reproject the roads to match the tracking data
ant_proj <- ANTEATER@info$projection
BR262 <- spTransform(BR262,ant_proj)

#Reproject the HR contour
HR_contour <- spTransform(HR_contour,ant_proj)

#Get the areas that fall on either side of the road
lpi <- gIntersection(HR_contour, BR262)
blpi <- gBuffer(lpi, width = 0.000001)
dpi <- gDifference(HR_contour, blpi)
Side_1 <- SpatialPolygons(list(Polygons(list(dpi@polygons[[1]]@Polygons[[1]]), "1")))
Side_2 <- SpatialPolygons(list(Polygons(list(dpi@polygons[[1]]@Polygons[[2]]), "2")))

#Plot the split HR and the road
plot(HR_contour)
plot(Side_1, col = "lightgreen", add = TRUE)
plot(Side_2, col = "lightblue", add = TRUE)
lines(BR262, col = "red")
```

```
#Area on each side
Side_1 <- raster::area(Side_1)
Side_2 <- raster::area(Side_2)

#Ratio
RATIO <- min(Side_1, Side_2)/max(Side_1, Side_2)

round(RATIO,3)
```

```
## [1] 0.407
```

##### Distance of home range center to nearest road

```
#Extract coordinates of home range center and carry out some reprojections
pj <- proj4::project(FIT$mu,
                     FIT@info$projection,
                     inverse = TRUE)
mu <- data.frame(lat=pj[,1],
                 lon=pj[,2])
mu <- SpatialPointsDataFrame(coords = mu,
                             data = mu,
                             proj4string=CRS("+proj=longlat +datum=WGS84 +ellps=WGS84 +towgs84=0,0,0"))
mu_proj <- spTransform(mu,CRS("+proj=utm +zone=21 +south +ellps=WGS72 +units=m +no_defs"))
```

```
## Warning in showSRID(uprojargs, format = "PROJ", multiline = "NO", prefer_proj
## = prefer_proj): Discarded datum Unknown based on WGS 72 ellipsoid in Proj4
## definition
```

```
ROAD_proj <- spTransform(BR262, CRS("+proj=utm +zone=21 +south +ellps=WGS72 +units=m +no_defs"))
```

```
## Warning in showSRID(uprojargs, format = "PROJ", multiline = "NO", prefer_proj
## = prefer_proj): Discarded datum Unknown based on WGS 72 ellipsoid in Proj4
## definition
```

```
#Calculate distance to nearest road
gDistance(mu_proj, ROAD_proj)/1000
```

```
## [1] 0.3610817
```

```
#Plot everything to make sure projections are correct
HR_contour_proj <- spTransform(HR_contour,CRS("+proj=utm +zone=21 +south +ellps=WGS72 +units=m +no_defs"))
```

```
## Warning in showSRID(uprojargs, format = "PROJ", multiline = "NO", prefer_proj
## = prefer_proj): Discarded datum Unknown based on WGS 72 ellipsoid in Proj4
## definition
```

```
plot(HR_contour_proj)
plot(ROAD_proj, add = TRUE, col = "red")
plot(mu_proj, add = TRUE, pch = 16, col = "#046C9A")
```

From this we see that the center of this animal’s home range was 0.361 km from the nearest road (BR 262).

##### Paved road Crossings

We further estimated the total number of crossings across the highways for each giant anteater. For this, we used each animal’s tracking data and their CTMM to reconstruct the most likely path that they traveled through the landscape over the course of the study period. We identified the total number of intersections (crossings) between the most likely path and the different roads.

```
#Estimate the most likely path based on the fitted movement model
mlp <- predict(ANTEATER, FIT, dt = 60, complete = TRUE)
```

```
## Warning in if (axes == "z") {: the condition has length > 1 and only the first
## element will be used
```

```
#Convert to the right format for counting road crossings
MLP <- SpatialPointsDataFrame.telemetry(mlp)
MLP <-  spTransform(MLP,ant_proj)
MLP_2 <- lapply(split(MLP, MLP$identity),
                function(x) Lines(list(Line(coordinates(x))), MLP$identity[1L]))
MLP_2 <- SpatialLines(MLP_2, proj4string = BR262@proj4string)

#How many times does it cross the paved road
BR262_crossings <- rgeos::gIntersection(MLP_2, BR262)
Num_Crossings <- length(BR262_crossings)

Num_Crossings
```

```
## [1] 46
```

```
#Plot of the most likely path and the road
plot(MLP_2, col = "NA")
lines(BR262, col = "#FF0000")
lines(MLP_2, col = "#046C9A")
title(main = paste(ANTEATER@info$identity, "  -   BR262 Crossings: ", Num_Crossings),
      family = "serif",
      font.main = 1,
      cex.main = 0.85)
```

From these calculations, we see that this animal crossed highway BR262 a total of 46 times. In addition to estimating the number of crossings, we quantified the times that each of these crossings occurred.

```
#Requires conversion to lat long for the distHaversine function
BR262_crossings_latlong <- spTransform(BR262_crossings,"+proj=longlat +datum=WGS84 +no_defs +ellps=WGS84 +towgs84=0,0,0")
```

```
## Warning in validityMethod(object): duplicate rownames are interpreted by rgeos
## as MultiPoints; use SpatialMultiPoints to define these; in future sp versions
## this warning will become an error
```

```
MLP_latlong <-  spTransform(MLP,"+proj=longlat +datum=WGS84 +no_defs +ellps=WGS84 +towgs84=0,0,0")

#Find times it crossed BR262
cross_times <- vector()
for(i in 1:nrow(BR262_crossings@coords)){
  #Find which point in the mlp is closest to the crossing location
  DISTS <- geosphere::distHaversine(BR262_crossings_latlong@coords[i,], MLP_latlong@coords)
  cross_times[i] <- MLP@data[which(DISTS == min(DISTS)),"timestamp"]
}

head(cross_times)
```

```
## [1] "2018-11-13 05:15:07.750000 UTC" "2018-11-13 04:44:14.799999 UTC"
## [3] "2018-08-28 13:53:33.599999 UTC" "2018-08-28 14:18:33.150000 UTC"
## [5] "2018-11-26 15:24:04.400000 UTC" "2018-10-08 05:07:11.262500 UTC"
```

##### Stream Crossings

We estimated the total number of crossings across local streams for each giant anteater. For this, we used each animal’s tracking data and their CTMM to reconstruct the most likely path that they traveled through the landscape over the course of the study period as above. We identified the total number of intersections (crossings) between the most likely path and local streams.

```
# Load the locations of the dirt roads
streams <- readOGR(dsn = "~/Dropbox (Personal)/UBC/Side_Projects/Arnaud_Anteaters/Scripts/Roads/Streams")
```

```
## OGR data source with driver: ESRI Shapefile 
## Source: "/Users/michaelnoonan/Dropbox (Personal)/UBC/Side_Projects/Arnaud_Anteaters/Scripts/Roads/Streams", layer: "Streams"
## with 895 features
## It has 8 fields
```

```
#Reproject the dirt roads
streams <- spTransform(streams,ant_proj)

#How many times does it cross streams
stream_crossings <- rgeos::gIntersection(MLP_2, streams)

#Total number of crossings
length(stream_crossings)
```

```
## [1] 16
```

```
#Plot of the most likely path and the local streams
# The road is shown in red for reference, not used in these calculations
plot(MLP_2, col = "NA")
lines(BR262, col = "#FF0000")
lines(streams, col = "lightblue")
lines(MLP_2, col = "#046C9A")
title(main = paste(ANTEATER@info$identity, "  -   Stream Crossings: ", length(stream_crossings)),
      family = "serif",
      font.main = 1,
      cex.main = 0.85)
```

From these calculations, we see that this animal crossed streams a total of 16 times.

##### Crossing Passages

From the track data, we classified crossings on each highway that were within 20 m (the median GPS measurement error) of a road passage as crossings where a giant anteater could potentially have used that structure to move across the road. This allowed us to search for evidence that anteaters tend to cross the roads through existing passages and/or tend to cross high traffic roads in periods of lower traffic intensity (Q3).

```
# Load in the locations of the crossing passages
passes <- readOGR(dsn = "~/Dropbox (Personal)/UBC/Side_Projects/Arnaud_Anteaters/Scripts/Roads/Passages")
```

```
## OGR data source with driver: ESRI Shapefile 
## Source: "/Users/michaelnoonan/Dropbox (Personal)/UBC/Side_Projects/Arnaud_Anteaters/Scripts/Roads/Passages", layer: "PassagesRoads"
## with 29 features
## It has 6 fields
## Integer64 fields read as strings:  ID
```

```
passes <- spTransform(passes,ant_proj)

#Empty vector to store results
pass_dists <- vector("numeric", length = length(BR262_crossings))

#Measuring distance of crossings from passages
for(i in 1:nrow(BR262_crossings@coords)){
  #Find which point in the mlp is clsoest to the crossing location
  DISTS <- raster::pointDistance(BR262_crossings@coords[i,], passes@coords, lonlat = FALSE)
  pass_dists[i] <- min(DISTS)
}

head(pass_dists)
```

```
## [1] 1146.4902  857.4086  211.2971  270.6215  294.3660  770.6589
```

```
res <- t.test(pass_dists)
res
```

```
## 
##  One Sample t-test
## 
## data:  pass_dists
## t = 16.706, df = 45, p-value < 2.2e-16
## alternative hypothesis: true mean is not equal to 0
## 95 percent confidence interval:
##  1570.207 2000.725
## sample estimates:
## mean of x 
##  1785.466
```

```
#Which crossings were within 20m of a road passage
which(pass_dists <= 20)
```

```
## integer(0)
```

```
#Plot of the most likely path, the road, and the passage structures
plot(MLP_2, col = "NA")
lines(BR262, col = "#FF0000")
lines(MLP_2, col = "#046C9A")
points(passes, pch = 16)
title(main = paste(ANTEATER@info$identity, "  -   BR262 Crossings: ", Num_Crossings),
      family = "serif",
      font.main = 1,
      cex.main = 0.85)
```

From these calculations we see that, on average, this animal crossed the road 1785.47 m from the nearest passage structure (95% CI: 1570.21 - 2000.72). For this individual, no road crossings were within 20 m of a passage structure, and the crossing that was closest to a passage structure was still 211.3m away.

---

#### Simulated Crossings

In a complementary approach, we used simulations to generate a null model of the number of crossings across all roads (paved and unpaved) and streams that would be expected by chance. For each giant anteater tracked, we used their CTMM model to simulate 100 movement datasets with sampling times that matched the empirical data. We then quantified the number and time of day of road and stream crossings in these 100 simulated datasets and averaged the results for each individual. This provided an estimate of the number of road crossings that would be expected by random movement within individuals’ home ranges that we could compare our empirical results against.

```
library(foreach)
library(doParallel)

#########################################################
# Set up the paralellisation

#Reg. multiple cores for DoParallel
nCores <- 8
registerDoParallel(nCores)
#Check that it's setup correctly
getDoParWorkers()
```

```
## [1] 8
```

```
#A character string for reprojections
LatLon <- "+proj=longlat +datum=WGS84 +no_defs +ellps=WGS84 +towgs84=0,0,0"

#Number of simulated replicates per animal
nReps <- 1000

#Run the simulations for this animal
x <- foreach(j=1:nReps) %dopar% {
  
  ###############################################
  # Simulate data from the fitted movement model
  ###############################################    
  
  SIM <- simulate(FIT, t = ANTEATER$t, complete = TRUE)
  
  #Convert to the right format/projection for identifying road crossings using rgeos
  MLP <- SpatialPointsDataFrame.telemetry(SIM)
  MLP <-  spTransform(MLP, ANTEATER@info$projection)
  MLP_2 <- lapply(split(MLP, MLP$identity),
                  function(x) Lines(list(Line(coordinates(x))), MLP$identity[1L]))
  MLP_2 <- SpatialLines(MLP_2, proj4string = BR262@proj4string)
  
  ###############################################      
  # The number of crossings for the simulated animal
  ###############################################      
  
  #How many times does it cross the road
  Sim_Road_Cross <- rgeos::gIntersection(MLP_2, BR262)
  Sim_Road_Cross_Count <- length(Sim_Road_Cross)
  
  #How many times does it cross streams
  Sim_Stream_Cross <- rgeos::gIntersection(MLP_2, streams)
  Sim_Stream_Cross_Count <- length(Sim_Stream_Cross)
  
  ###############################################      
  # Crossing times for the simulated animal
  ###############################################      
  
  #Requires conversion to lat long for the distHaversine function
  Road_crossings_latlong <- spTransform(Sim_Road_Cross,LatLon)
  Sim_Stream_Cross_latlong <- spTransform(Sim_Stream_Cross,LatLon)
  
  MLP_latlong <-  spTransform(MLP,LatLon)
  
  #Find times it crossed the road
  road_cross_times <- vector()
  for(i in 1:nrow(BR262_crossings@coords)){
    #Find which point in the mlp is closest to the crossing location
    DISTS <- geosphere::distHaversine(BR262_crossings_latlong@coords[i,], MLP_latlong@coords)
    road_cross_times[i] <- MLP@data[which(DISTS == min(DISTS)),"timestamp"]
  }
  
  #Find times it crossed the stream
  stream_cross_times <- vector()
  for(i in 1:nrow(Sim_Stream_Cross@coords)){
    #Find which point in the mlp is closest to the crossing location
    DISTS <- geosphere::distHaversine(Sim_Stream_Cross_latlong@coords[i,], MLP_latlong@coords)
    stream_cross_times[i] <- MLP@data[which(DISTS == min(DISTS)),"timestamp"]
  }
  
  #################################
  #list of results to return
  list(FIT@info$identity,
       Sim_Road_Cross_Count,
       Sim_Stream_Cross_Count,
       road_cross_times,
       stream_cross_times)
}

#Clean up results
RESULTS <- data.frame("ID" = unlist(lapply(x, function (x) x[1])),
                      "Road_Crossings" = unlist(lapply(x, function (x) x[2])),
                      "Stream_Crossings"= unlist(lapply(x, function (x) x[3])))

head(RESULTS)
```

```
##             ID Road_Crossings Stream_Crossings
## 1 Jennifer 262            184              113
## 2 Jennifer 262            244               51
## 3 Jennifer 262            205              112
## 4 Jennifer 262            172               33
## 5 Jennifer 262            156               72
## 6 Jennifer 262            163               27
```

```
#Mean road and stream crossings
mean(RESULTS$Road_Crossings); mean(RESULTS$Stream_Crossings)
```

```
## [1] 187.497
```

```
## [1] 65.239
```

From these results we can see that random movement within home ranges with data collected at the sampled times would have results in an expected 187.497 road crossings, and 65.239 stream crossings. In comparison this animal actually crossed highway BR262 a total of 46 times, and crossed streams a total of 16 times. Simulated crossing times are not shown here.

---

#### References

Calabrese JM, Fleming CH, Gurarie E. ctmm: an R package for analyzing animal relocation data as a continuous-time stochastic process. (2016). Methods in Ecology and Evolution, 7(9):1124– 1132.

Fleming, C. H., & Calabrese, J. M. (2017). A new kernel density estimator for accurate home‐range and species‐range area estimation. Methods in Ecology and Evolution, 8(5), 571-579.

Fleming, C. H., Fagan, W. F., Mueller, T., Olson, K. A., Leimgruber, P., & Calabrese, J. M. (2015). Rigorous home range estimation with movement data: a new autocorrelated kernel density estimator. Ecology, 96(5), 1182-1188.

Fleming, C. H., Noonan, M. J., Medici, E. P., & Calabrese, J. M. (2019). Overcoming the challenge of small effective sample sizes in home‐range estimation. Methods in Ecology and Evolution, 10(10), 1679-1689.

Noonan, M. J., Fleming, C. H., Akre, T. S., Drescher-Lehman, J., Gurarie, E., Harrison, A. L., … & Calabrese, J. M. (2019). Scale-insensitive estimation of speed and distance traveled from animal tracking data. Movement ecology, 7(1), 1-15.
