## Appendix S2 for "Roads as ecological traps for giant anteaters": Appendix_S2.html

Appendix S2 - Descriptive Analyses of Movement Metrics and Road Interactions


### Appendix S2 - Descriptive Analyses of Movement Metrics and Road Interactions

###### Michael J. Noonan, Fernando Ascensão, Débora R. Yogui, Arnaud L.J. Desbiez

This document was created on April 02, 2021.

---

#### Overview

In this appendix we detail the analyses that were carried out on the individual movement metrics to arrive at the results presented in the main text. We also provide Table S2.1, which provides information on road-killed animals that were included in this study.

##### Mortality of individuals tracked during the study period

**Table S2.1** Cause of death and number of individuals of giant anteaters tracked during the study

| Cause of Death | N | ID |
| --- | --- | --- |
| Roadkill after collar removal. On the road. ID through microchip | 2 | Ed, Larry |
| Roadkill with the collar. On the road. | 3 | Pequi, Ben. Christoffer |
| Roadkill with the collar. Away from the road. | 1 | Schwartz |
| Death during the study for unknown causes | 2 | Yoki, Segre |

---

##### Data Import and Pre-Analysis

The first step of the descriptive analyses was importing the dataset of movement metrics, ensuring the import was carried out correctly (e.g., numbers are numbers, factors are factors, etc.), and check for any correlations among the variables that could cause multi-collinearity issues.

```
#Load in the necessary packages
library(metafor)
library(ggplot2)
library(gridExtra)
library(MuMIn)
library(DAAG)
library(nlme)
library(lubridate)

#Import the dataset of individual movement metrics
data <- read.csv("Results/Anteater_Results_Final.csv")

#Re-order based on location
data <- data[order(data$Road),]

#Convert factor variables to factors
data$Sex <- as.factor(data$Sex)
data$Road <- as.factor(data$Road)
```

##### Pre-Analysis

As a preliminary check, we inspected the data for any correlations that would result in issues of multi-collinearity when modelling these data.

```
#Correlation between parameters used in the analyses
round(cor(data[,c("HR",  "Speed", "Dist_Rd", "Split", "Rd_Cr", "Strm_Cr", "Strm_Dist", "Forest", "Pasture", "Weight")]),2)
```

```
##              HR Speed Dist_Rd Split Rd_Cr Strm_Cr Strm_Dist Forest Pasture
## HR         1.00  0.14    0.03  0.41  0.20   -0.13     -0.01   0.33   -0.54
## Speed      0.14  1.00   -0.01  0.05  0.20    0.02     -0.14  -0.04   -0.03
## Dist_Rd    0.03 -0.01    1.00 -0.41 -0.39    0.11     -0.01  -0.07    0.06
## Split      0.41  0.05   -0.41  1.00  0.55   -0.13      0.00   0.14   -0.15
## Rd_Cr      0.20  0.20   -0.39  0.55  1.00   -0.04     -0.18   0.07   -0.13
## Strm_Cr   -0.13  0.02    0.11 -0.13 -0.04    1.00     -0.66   0.20   -0.16
## Strm_Dist -0.01 -0.14   -0.01  0.00 -0.18   -0.66      1.00  -0.25    0.24
## Forest     0.33 -0.04   -0.07  0.14  0.07    0.20     -0.25   1.00   -0.92
## Pasture   -0.54 -0.03    0.06 -0.15 -0.13   -0.16      0.24  -0.92    1.00
## Weight    -0.31  0.06   -0.23 -0.23 -0.14   -0.04      0.00  -0.28    0.23
##           Weight
## HR         -0.31
## Speed       0.06
## Dist_Rd    -0.23
## Split      -0.23
## Rd_Cr      -0.14
## Strm_Cr    -0.04
## Strm_Dist   0.00
## Forest     -0.28
## Pasture     0.23
## Weight      1.00
```

There was a strong negative correlation between the amount of natural forest and pasture in an individual’s home range (Pearson correlation: -0.92). We therefore excluded natural forest cover from all subsequent analyses. No other variables exhibited correlations that warranted special handling.

---

#### Basic summary of the movement metrics

```
#Mean HR size 
DAT <- escalc(measure = "MN",
              mi = HR/1000000,
              sdi = sqrt(HR_var/1000000),
              ni = n_area,
              data = data)

summary(rma(yi ~ 1, vi, data=DAT, method="REML"))
```

```
## 
## Random-Effects Model (k = 38; tau^2 estimator: REML)
## 
##    logLik   deviance        AIC        BIC       AICc 
## -124.3715   248.7430   252.7430   255.9648   253.0959   
## 
## tau^2 (estimated amount of total heterogeneity): 48.6253 (SE = 11.3065)
## tau (square root of estimated tau^2 value):      6.9732
## I^2 (total heterogeneity / total variability):   100.00%
## H^2 (total variability / sampling variability):  57973.13
## 
## Test for Heterogeneity:
## Q(df = 37) = 232001.2141, p-val < .0001
## 
## Model Results:
## 
## estimate      se    zval    pval   ci.lb   ci.ub 
##   6.8090  1.1313  6.0189  <.0001  4.5918  9.0262  *** 
## 
## ---
## Signif. codes:  0 '***' 0.001 '**' 0.01 '*' 0.05 '.' 0.1 ' ' 1
```

```
#Does HR size differ between the sexes?
summary(rma(yi ~ Sex, vi, data=DAT, method="REML"))
```

```
## 
## Mixed-Effects Model (k = 38; tau^2 estimator: REML)
## 
##    logLik   deviance        AIC        BIC       AICc 
## -119.5899   239.1798   245.1798   249.9304   245.9298   
## 
## tau^2 (estimated amount of residual heterogeneity):     44.9348 (SE = 10.5926)
## tau (square root of estimated tau^2 value):             6.7033
## I^2 (residual heterogeneity / unaccounted variability): 100.00%
## H^2 (unaccounted variability / sampling variability):   51595.08
## R^2 (amount of heterogeneity accounted for):            7.59%
## 
## Test for Residual Heterogeneity:
## QE(df = 36) = 210513.8747, p-val < .0001
## 
## Test of Moderators (coefficient 2):
## QM(df = 1) = 4.0382, p-val = 0.0445
## 
## Model Results:
## 
##          estimate      se    zval    pval   ci.lb   ci.ub 
## intrcpt    4.3802  1.6258  2.6941  0.0071  1.1936  7.5668  ** 
## SexMale    4.3951  2.1871  2.0095  0.0445  0.1084  8.6817   * 
## 
## ---
## Signif. codes:  0 '***' 0.001 '**' 0.01 '*' 0.05 '.' 0.1 ' ' 1
```

```
#Does HR size differ with body size?
summary(rma(yi ~ Weight, vi, data=DAT, method="REML"))
```

```
## 
## Mixed-Effects Model (k = 38; tau^2 estimator: REML)
## 
##    logLik   deviance        AIC        BIC       AICc 
## -119.7050   239.4101   245.4101   250.1606   246.1601   
## 
## tau^2 (estimated amount of residual heterogeneity):     45.2214 (SE = 10.6601)
## tau (square root of estimated tau^2 value):             6.7247
## I^2 (residual heterogeneity / unaccounted variability): 100.00%
## H^2 (unaccounted variability / sampling variability):   52045.15
## R^2 (amount of heterogeneity accounted for):            7.00%
## 
## Test for Residual Heterogeneity:
## QE(df = 36) = 203699.9539, p-val < .0001
## 
## Test of Moderators (coefficient 2):
## QM(df = 1) = 3.7831, p-val = 0.0518
## 
## Model Results:
## 
##          estimate      se     zval    pval    ci.lb    ci.ub 
## intrcpt   25.3806  9.6105   2.6409  0.0083   6.5444  44.2168  ** 
## Weight    -0.5826  0.2995  -1.9450  0.0518  -1.1697   0.0045   . 
## 
## ---
## Signif. codes:  0 '***' 0.001 '**' 0.01 '*' 0.05 '.' 0.1 ' ' 1
```

```
#Does HR size differ with amount of pasture?
summary(rma(yi ~ Pasture, vi, data=DAT, method="REML"))
```

```
## 
## Mixed-Effects Model (k = 38; tau^2 estimator: REML)
## 
##    logLik   deviance        AIC        BIC       AICc 
## -115.4291   230.8582   236.8582   241.6088   237.6082   
## 
## tau^2 (estimated amount of residual heterogeneity):     35.6513 (SE = 8.4044)
## tau (square root of estimated tau^2 value):             5.9709
## I^2 (residual heterogeneity / unaccounted variability): 100.00%
## H^2 (unaccounted variability / sampling variability):   42201.30
## R^2 (amount of heterogeneity accounted for):            26.68%
## 
## Test for Residual Heterogeneity:
## QE(df = 36) = 208313.1045, p-val < .0001
## 
## Test of Moderators (coefficient 2):
## QM(df = 1) = 14.4507, p-val = 0.0001
## 
## Model Results:
## 
##          estimate      se     zval    pval    ci.lb    ci.ub 
## intrcpt   34.1590  7.2598   4.7052  <.0001  19.9301  48.3879  *** 
## Pasture   -0.3113  0.0819  -3.8014  0.0001  -0.4719  -0.1508  *** 
## 
## ---
## Signif. codes:  0 '***' 0.001 '**' 0.01 '*' 0.05 '.' 0.1 ' ' 1
```

```
# Subset to the animals living within 2km of roads
# and fit a model predicting distance from road by anteater traits
data_Close <- data[data$Dist_Rd<2,]
FIT <- gls(Dist_Rd ~ Road + Sex + Weight,
           data = data_Close,
           na.action = na.fail,
           method = "ML")

head(dredge(FIT), 10)
```

```
## Global model call: gls(model = Dist_Rd ~ Road + Sex + Weight, data = data_Close, 
##     method = "ML", na.action = na.fail)
## ---
## Model selection table 
##   (Intrc) Road Sex    Weght df  logLik AICc delta weight
## 1  0.8951                    2 -15.212 34.9  0.00  0.569
## 3  0.8711        +           3 -15.178 37.4  2.47  0.165
## 5  0.7539          0.004360  3 -15.192 37.4  2.50  0.163
## 7  0.6956        + 0.005313  4 -15.148 40.1  5.19  0.042
## 2  0.8899    +               4 -15.200 40.2  5.29  0.040
## 6  0.6002    +     0.008318  5 -15.154 43.2  8.24  0.009
## 4  0.8688    +   +           5 -15.163 43.2  8.26  0.009
## 8  0.5654    +   + 0.008680  6 -15.113 46.4 11.50  0.002
## Models ranked by AICc(x)
```

```
INTERCEPT <- gls(Dist_Rd ~ 1,
                 data = data_Close,
                 na.action = na.fail,
                 method = "ML")
```

---

#### Question 1) Does the movement behaviour of giant anteaters differ when living near paved roads?

The first question we addressed was whether the home range size and/or daily movement distances of giant anteaters differed across the different study sites, and whether there was a relationship between these movement metrics and the distance they lived to a road. Comparisons were performed using the meta-regression model implemented in the `R` package `metafor`, which allowed uncertainty in each individual estimate to be propagated into the population level estimate when making comparisons.

```
#####################################################################
#Set up the dataset for testing differences in HR size between the three sites
DAT <- escalc(measure = "MN",
              mi = HR/1000000,
              sdi = sqrt(HR_var/1000000),
              ni = n_area,
              data = data)

#Run the meta-regression model testing differences in HR size between the three sites
Site_HR_res <- rma(yi ~ Road, vi, data=DAT, method="ML")
Site_HR_Null <- rma(yi ~ 1, vi, data=DAT, method="ML")

#likelihood ratio test
anova(Site_HR_res,
      Site_HR_Null)
```

```
## 
##         df      AIC      BIC     AICc    logLik    LRT   pval          QE 
## Full     4 262.0679 268.6182 263.2800 -127.0339               172281.1651 
## Reduced  2 258.4517 261.7269 258.7946 -127.2259 0.3838 0.8254 232001.2141 
##           tau^2     R^2 
## Full    46.8655         
## Reduced 47.3439 1.0104%
```

```
#Inspect the results
summary(rma(yi ~ Road, vi, data=DAT, method="REML"))
```

```
## 
## Mixed-Effects Model (k = 38; tau^2 estimator: REML)
## 
##    logLik   deviance        AIC        BIC       AICc 
## -118.4451   236.8902   244.8902   251.1116   246.2235   
## 
## tau^2 (estimated amount of residual heterogeneity):     50.8888 (SE = 12.1661)
## tau (square root of estimated tau^2 value):             7.1336
## I^2 (residual heterogeneity / unaccounted variability): 100.00%
## H^2 (unaccounted variability / sampling variability):   58604.68
## R^2 (amount of heterogeneity accounted for):            0.00%
## 
## Test for Residual Heterogeneity:
## QE(df = 35) = 172281.1651, p-val < .0001
## 
## Test of Moderators (coefficients 2:3):
## QM(df = 2) = 0.3552, p-val = 0.8373
## 
## Model Results:
## 
##             estimate      se    zval    pval    ci.lb   ci.ub 
## intrcpt       5.9992  1.9067  3.1464  0.0017   2.2622  9.7363  ** 
## RoadBR262     0.6836  3.3024  0.2070  0.8360  -5.7890  7.1563     
## RoadMS-040    1.5287  2.5747  0.5937  0.5527  -3.5177  6.5750     
## 
## ---
## Signif. codes:  0 '***' 0.001 '**' 0.01 '*' 0.05 '.' 0.1 ' ' 1
```

```
#Predicted home range areas of animals living in each of the three sites
predict(Site_HR_res)[c(1,15,22),]
```

```
## 
##      pred     se  ci.lb   ci.ub   cr.lb   cr.ub 
## 1  5.9990 1.8298 2.4126  9.5853 -7.8897 19.8876 
## 15 6.6828 2.5876 1.6112 11.7545 -7.6613 21.0270 
## 22 7.5279 1.6604 4.2735 10.7822 -6.2788 21.3345
```

```
#Run the meta-regression model testing differences in HR size with distance to road
Dist_HR_res <- rma(yi ~ Dist_Rd, vi, data=DAT, method="REML")

#Inspect the results
summary(Dist_HR_res)
```

```
## 
## Mixed-Effects Model (k = 38; tau^2 estimator: REML)
## 
##    logLik   deviance        AIC        BIC       AICc 
## -121.4923   242.9846   248.9846   253.7352   249.7346   
## 
## tau^2 (estimated amount of residual heterogeneity):     49.9465 (SE = 11.7738)
## tau (square root of estimated tau^2 value):             7.0673
## I^2 (residual heterogeneity / unaccounted variability): 100.00%
## H^2 (unaccounted variability / sampling variability):   57571.36
## R^2 (amount of heterogeneity accounted for):            0.00%
## 
## Test for Residual Heterogeneity:
## QE(df = 36) = 214243.8429, p-val < .0001
## 
## Test of Moderators (coefficient 2):
## QM(df = 1) = 0.0226, p-val = 0.8805
## 
## Model Results:
## 
##          estimate      se    zval    pval    ci.lb    ci.ub 
## intrcpt    6.6025  1.7892  3.6902  0.0002   3.0957  10.1092  *** 
## Dist_Rd    0.1194  0.7941  0.1504  0.8805  -1.4370   1.6759      
## 
## ---
## Signif. codes:  0 '***' 0.001 '**' 0.01 '*' 0.05 '.' 0.1 ' ' 1
```

```
#####################################################################
#Differences in movement speeds between the three sites
DAT <- data.frame(id = data$ID,
                  yi = data$Speed*86.4,
                  vi = data$Speed_var*86.4,
                  Road = data$Road,
                  Dist_Rd = data$Dist_Rd)

#Run the meta-regression model testing differences in speed between the three sites
Site_Speed_res <- rma(yi, vi, mods = ~ Road, data=DAT, method="ML")
Site_Speed_Null <- rma(yi ~ 1, vi, data=DAT, method="ML")

#likelihood ratio test
anova(Site_Speed_res,
      Site_Speed_Null)
```

```
## 
##         df      AIC      BIC     AICc   logLik    LRT   pval           QE 
## Full     4 115.1086 121.6589 116.3207 -53.5543               2770622.0248 
## Reduced  2 114.9830 118.2581 115.3258 -55.4915 3.8744 0.1441 3053327.5017 
##          tau^2     R^2 
## Full    0.9809         
## Reduced 1.0862 9.6950%
```

```
#Inspect the results
summary(rma(yi, vi, mods = ~ Road, data=DAT, method="REML"))
```

```
## 
## Mixed-Effects Model (k = 38; tau^2 estimator: REML)
## 
##   logLik  deviance       AIC       BIC      AICc 
## -50.7655  101.5310  109.5310  115.7524  110.8644   
## 
## tau^2 (estimated amount of residual heterogeneity):     1.0650 (SE = 0.2546)
## tau (square root of estimated tau^2 value):             1.0320
## I^2 (residual heterogeneity / unaccounted variability): 100.00%
## H^2 (unaccounted variability / sampling variability):   74300.81
## R^2 (amount of heterogeneity accounted for):            4.53%
## 
## Test for Residual Heterogeneity:
## QE(df = 35) = 2770622.0248, p-val < .0001
## 
## Test of Moderators (coefficients 2:3):
## QM(df = 2) = 3.7568, p-val = 0.1528
## 
## Model Results:
## 
##             estimate      se     zval    pval    ci.lb   ci.ub 
## intrcpt       7.0957  0.2758  25.7266  <.0001   6.5551  7.6363  *** 
## RoadBR262     0.8024  0.4777   1.6797  0.0930  -0.1339  1.7388    . 
## RoadMS-040   -0.0582  0.3724  -0.1562  0.8759  -0.7882  0.6718      
## 
## ---
## Signif. codes:  0 '***' 0.001 '**' 0.01 '*' 0.05 '.' 0.1 ' ' 1
```

```
#Predicted movement speeds of animals in each of the three sites
predict(Site_Speed_res)[c(1,15,22),]
```

```
## 
##      pred     se  ci.lb  ci.ub  cr.lb  cr.ub 
## 1  7.0957 0.2647 6.5769 7.6145 5.0864 9.1050 
## 15 7.8981 0.3744 7.1644 8.6319 5.8229 9.9733 
## 22 7.0375 0.2402 6.5667 7.5083 5.0401 9.0349
```

```
#Run the meta-regression model testing differences in mean speed with dist of HR center to road
Dist_Speed_res <- rma(yi, vi, mods = ~ Dist_Rd, data=DAT, method="REML")

#Inspect the results
summary(Dist_Speed_res)
```

```
## 
## Mixed-Effects Model (k = 38; tau^2 estimator: REML)
## 
##   logLik  deviance       AIC       BIC      AICc 
## -53.5408  107.0816  113.0816  117.8322  113.8316   
## 
## tau^2 (estimated amount of residual heterogeneity):     1.1463 (SE = 0.2702)
## tau (square root of estimated tau^2 value):             1.0707
## I^2 (residual heterogeneity / unaccounted variability): 100.00%
## H^2 (unaccounted variability / sampling variability):   78533.95
## R^2 (amount of heterogeneity accounted for):            0.00%
## 
## Test for Residual Heterogeneity:
## QE(df = 36) = 2913802.8763, p-val < .0001
## 
## Test of Moderators (coefficient 2):
## QM(df = 1) = 0.0066, p-val = 0.9354
## 
## Model Results:
## 
##          estimate      se     zval    pval    ci.lb   ci.ub 
## intrcpt    7.2343  0.2710  26.6904  <.0001   6.7031  7.7656  *** 
## Dist_Rd   -0.0097  0.1203  -0.0810  0.9354  -0.2455  0.2260      
## 
## ---
## Signif. codes:  0 '***' 0.001 '**' 0.01 '*' 0.05 '.' 0.1 ' ' 1
```

##### Recreate Figure 2

```
#Home range size vs. study site

Fid2_a <- 
  ggplot(data) +
  ggtitle("a)") +
  scale_y_discrete(limits=data$ID) +
  annotate("rect",
           xmin = 0, xmax = Inf,
           ymin = -Inf, ymax = 14.5, fill = "grey70",
           alpha = .15) +
  
  annotate("rect",
           xmin = 0, xmax = Inf,
           ymin = 14.5, ymax = 21.5, fill = "grey50",
           alpha = .15) +
  annotate("rect",
           xmin = 0, xmax = Inf,
           ymin = 21.5, ymax = Inf, fill = "grey20",
           alpha = .15) +
  
  theme_bw() +
  
  #BR_267
  geom_segment(aes(x = HR_min[1]/1000000, xend = HR_max[1]/1000000, y = 1, yend = 1),size = 1.5, colour = "#e6c141", alpha = 0.05) +
  geom_segment(aes(x = HR_min[2]/1000000, xend = HR_max[2]/1000000, y = 2, yend = 2),size = 1.5, colour = "#e6c141", alpha = 0.05) +
  geom_segment(aes(x = HR_min[3]/1000000, xend = HR_max[3]/1000000, y = 3, yend = 3),size = 1.5, colour = "#e6c141", alpha = 0.05) +
  geom_segment(aes(x = HR_min[4]/1000000, xend = HR_max[4]/1000000, y = 4, yend = 4),size = 1.5, colour = "#e6c141", alpha = 0.05) +
  geom_segment(aes(x = HR_min[5]/1000000, xend = HR_max[5]/1000000, y = 5, yend = 5),size = 1.5, colour = "#e6c141", alpha = 0.05) +
  geom_segment(aes(x = HR_min[6]/1000000, xend = HR_max[6]/1000000, y = 6, yend = 6),size = 1.5, colour = "#e6c141", alpha = 0.05) +
  geom_segment(aes(x = HR_min[7]/1000000, xend = HR_max[7]/1000000, y = 7, yend = 7),size = 1.5, colour = "#e6c141", alpha = 0.05) +
  geom_segment(aes(x = HR_min[8]/1000000, xend = HR_max[8]/1000000, y = 8, yend = 8),size = 1.5, colour = "#e6c141", alpha = 0.05) +
  geom_segment(aes(x = HR_min[9]/1000000, xend = HR_max[9]/1000000, y = 9, yend = 9),size = 1.5, colour = "#e6c141", alpha = 0.05) +
  geom_segment(aes(x = HR_min[10]/1000000, xend = HR_max[10]/1000000, y = 10, yend = 10),size = 1.5, colour = "#e6c141", alpha = 0.05) +
  geom_segment(aes(x = HR_min[11]/1000000, xend = HR_max[11]/1000000, y = 11, yend = 11),size = 1.5, colour = "#e6c141", alpha = 0.05) +
  geom_segment(aes(x = HR_min[12]/1000000, xend = HR_max[12]/1000000, y = 12, yend = 12),size = 1.5, colour = "#e6c141", alpha = 0.05) +
  geom_segment(aes(x = HR_min[13]/1000000, xend = HR_max[13]/1000000, y = 13, yend = 13),size = 1.5, colour = "#e6c141", alpha = 0.05) +
  geom_segment(aes(x = HR_min[14]/1000000, xend = HR_max[14]/1000000, y = 14, yend = 14),size = 1.5, colour = "#e6c141", alpha = 0.05) +
  
  #BR_262
  geom_segment(aes(x = HR_min[15]/1000000, xend = HR_max[15]/1000000, y = 15, yend = 15),size = 1.5, colour = "#3471bc", alpha = 0.05) +
  geom_segment(aes(x = HR_min[16]/1000000, xend = HR_max[16]/1000000, y = 16, yend = 16),size = 1.5, colour = "#3471bc", alpha = 0.05) +
  geom_segment(aes(x = HR_min[17]/1000000, xend = HR_max[17]/1000000, y = 17, yend = 17),size = 1.5, colour = "#3471bc", alpha = 0.05) +
  geom_segment(aes(x = HR_min[18]/1000000, xend = HR_max[18]/1000000, y = 18, yend = 18),size = 1.5, colour = "#3471bc", alpha = 0.05) +
  geom_segment(aes(x = HR_min[19]/1000000, xend = HR_max[19]/1000000, y = 19, yend = 19),size = 1.5, colour = "#3471bc", alpha = 0.05) +
  geom_segment(aes(x = HR_min[20]/1000000, xend = HR_max[20]/1000000, y = 20, yend = 20),size = 1.5, colour = "#3471bc", alpha = 0.05) +
  geom_segment(aes(x = HR_min[21]/1000000, xend = HR_max[21]/1000000, y = 21, yend = 21),size = 1.5, colour = "#3471bc", alpha = 0.05) +
  
  #MS040
  geom_segment(aes(x = HR_min[22]/1000000, xend = HR_max[22]/1000000, y = 22, yend = 22),size = 1.5, colour = "#3c7a47", alpha = 0.05) +
  geom_segment(aes(x = HR_min[23]/1000000, xend = HR_max[23]/1000000, y = 23, yend = 23),size = 1.5, colour = "#3c7a47", alpha = 0.05) +
  geom_segment(aes(x = HR_min[24]/1000000, xend = HR_max[24]/1000000, y = 24, yend = 24),size = 1.5, colour = "#3c7a47", alpha = 0.05) +
  geom_segment(aes(x = HR_min[25]/1000000, xend = HR_max[25]/1000000, y = 25, yend = 25),size = 1.5, colour = "#3c7a47", alpha = 0.05) +
  geom_segment(aes(x = HR_min[26]/1000000, xend = HR_max[26]/1000000, y = 26, yend = 26),size = 1.5, colour = "#3c7a47", alpha = 0.05) +
  geom_segment(aes(x = HR_min[27]/1000000, xend = HR_max[27]/1000000, y = 27, yend = 27),size = 1.5, colour = "#3c7a47", alpha = 0.05) +
  geom_segment(aes(x = HR_min[28]/1000000, xend = HR_max[28]/1000000, y = 28, yend = 28),size = 1.5, colour = "#3c7a47", alpha = 0.05) +
  geom_segment(aes(x = HR_min[29]/1000000, xend = HR_max[29]/1000000, y = 29, yend = 29),size = 1.5, colour = "#3c7a47", alpha = 0.05) +
  geom_segment(aes(x = HR_min[30]/1000000, xend = HR_max[30]/1000000, y = 30, yend = 30),size = 1.5, colour = "#3c7a47", alpha = 0.05) +
  geom_segment(aes(x = HR_min[31]/1000000, xend = HR_max[31]/1000000, y = 31, yend = 31),size = 1.5, colour = "#3c7a47", alpha = 0.05) +
  geom_segment(aes(x = HR_min[32]/1000000, xend = HR_max[32]/1000000, y = 32, yend = 32),size = 1.5, colour = "#3c7a47", alpha = 0.05) +
  geom_segment(aes(x = HR_min[33]/1000000, xend = HR_max[33]/1000000, y = 33, yend = 33),size = 1.5, colour = "#3c7a47", alpha = 0.05) +
  geom_segment(aes(x = HR_min[34]/1000000, xend = HR_max[34]/1000000, y = 34, yend = 34),size = 1.5, colour = "#3c7a47", alpha = 0.05) +
  geom_segment(aes(x = HR_min[35]/1000000, xend = HR_max[35]/1000000, y = 35, yend = 35),size = 1.5, colour = "#3c7a47", alpha = 0.05) +
  geom_segment(aes(x = HR_min[36]/1000000, xend = HR_max[36]/1000000, y = 36, yend = 36),size = 1.5, colour = "#3c7a47", alpha = 0.05) +
  geom_segment(aes(x = HR_min[37]/1000000, xend = HR_max[37]/1000000, y = 37, yend = 37),size = 1.5, colour = "#3c7a47", alpha = 0.05) +
  geom_segment(aes(x = HR_min[38]/1000000, xend = HR_max[38]/1000000, y = 38, yend = 38),size = 1.5, colour = "#3c7a47", alpha = 0.05) +
  geom_point(aes(x = HR/1000000, y = ID), col = "white", size = 1) +
  geom_point(aes(x = HR/1000000, y = ID, col = Road), size = 0.7) +
  
  scale_color_manual(labels = c("BR 267", "BR 262", "MS040"), values = c("#e6c141", "#3471bc", "#3c7a47")) +
  
  theme(panel.grid.major = element_blank(),
        panel.grid.minor = element_blank(),
        axis.ticks.y = element_blank(),
        axis.title.y = element_blank(),
        axis.title.x = element_text(size=10, family = "serif"),
        axis.text.y = element_text(size=5, family = "serif"),
        axis.text.x = element_text(size=8, family = "serif"),
        plot.title = element_text(hjust = -0.025, size = 10, family = "serif"),
        legend.position=c(0.08, 0.9), legend.title=element_blank(),
        legend.text=element_text(size=6, family = "serif"),legend.key.size = unit(0.2, "cm"),
        legend.key.width = unit(0.025, "cm"), legend.background=element_blank(), 
        legend.key = element_blank(), legend.text.align = 0) +
  xlab(expression(paste("Home-range area (km"^2,")"))) +
  scale_x_log10(breaks = c(1,2.5,5, 10,25, 50,100), limits = c(1,60), expand = c(0,0))


#Home range size vs. distance to road
Fid2_b <- 
  ggplot(data) +
  ggtitle("b)") +
  theme_bw() +
  
  #BR_267
  geom_segment(aes(y = HR_min[1]/1000000, yend = HR_max[1]/1000000, x = Dist_Rd[1], xend = Dist_Rd[1]),size = 1.5, colour = "#e6c141", alpha = 0.05) +
  geom_segment(aes(y = HR_min[2]/1000000, yend = HR_max[2]/1000000, x = Dist_Rd[2], xend = Dist_Rd[2]),size = 1.5, colour = "#e6c141", alpha = 0.05) +
  geom_segment(aes(y = HR_min[3]/1000000, yend = HR_max[3]/1000000, x = Dist_Rd[3], xend = Dist_Rd[3]),size = 1.5, colour = "#e6c141", alpha = 0.05) +
  geom_segment(aes(y = HR_min[4]/1000000, yend = HR_max[4]/1000000, x = Dist_Rd[4], xend = Dist_Rd[4]),size = 1.5, colour = "#e6c141", alpha = 0.05) +
  geom_segment(aes(y = HR_min[5]/1000000, yend = HR_max[5]/1000000, x = Dist_Rd[5], xend = Dist_Rd[5]),size = 1.5, colour = "#e6c141", alpha = 0.05) +
  geom_segment(aes(y = HR_min[6]/1000000, yend = HR_max[6]/1000000, x = Dist_Rd[6], xend = Dist_Rd[6]),size = 1.5, colour = "#e6c141", alpha = 0.05) +
  geom_segment(aes(y = HR_min[7]/1000000, yend = HR_max[7]/1000000, x = Dist_Rd[7], xend = Dist_Rd[7]),size = 1.5, colour = "#e6c141", alpha = 0.05) +
  geom_segment(aes(y = HR_min[8]/1000000, yend = HR_max[8]/1000000, x = Dist_Rd[8], xend = Dist_Rd[8]),size = 1.5, colour = "#e6c141", alpha = 0.05) +
  geom_segment(aes(y = HR_min[9]/1000000, yend = HR_max[9]/1000000, x = Dist_Rd[9], xend = Dist_Rd[9]),size = 1.5, colour = "#e6c141", alpha = 0.05) +
  geom_segment(aes(y = HR_min[10]/1000000, yend = HR_max[10]/1000000, x = Dist_Rd[10], xend = Dist_Rd[10]),size = 1.5, colour = "#e6c141", alpha = 0.05) +
  geom_segment(aes(y = HR_min[11]/1000000, yend = HR_max[11]/1000000, x = Dist_Rd[11], xend = Dist_Rd[11]),size = 1.5, colour = "#e6c141", alpha = 0.05) +
  geom_segment(aes(y = HR_min[12]/1000000, yend = HR_max[12]/1000000, x = Dist_Rd[12], xend = Dist_Rd[12]),size = 1.5, colour = "#e6c141", alpha = 0.05) +
  geom_segment(aes(y = HR_min[13]/1000000, yend = HR_max[13]/1000000, x = Dist_Rd[13], xend = Dist_Rd[13]),size = 1.5, colour = "#e6c141", alpha = 0.05) +
  geom_segment(aes(y = HR_min[14]/1000000, yend = HR_max[14]/1000000, x = Dist_Rd[14], xend = Dist_Rd[14]),size = 1.5, colour = "#e6c141", alpha = 0.05) +
  
  #BR_262
  geom_segment(aes(y = HR_min[15]/1000000, yend = HR_max[15]/1000000, x = Dist_Rd[15], xend = Dist_Rd[15]),size = 1.5, colour = "#3471bc", alpha = 0.05) +
  geom_segment(aes(y = HR_min[16]/1000000, yend = HR_max[16]/1000000, x = Dist_Rd[16], xend = Dist_Rd[16]),size = 1.5, colour = "#3471bc", alpha = 0.05) +
  geom_segment(aes(y = HR_min[17]/1000000, yend = HR_max[17]/1000000, x = Dist_Rd[17], xend = Dist_Rd[17]),size = 1.5, colour = "#3471bc", alpha = 0.05) +
  geom_segment(aes(y = HR_min[18]/1000000, yend = HR_max[18]/1000000, x = Dist_Rd[18], xend = Dist_Rd[18]),size = 1.5, colour = "#3471bc", alpha = 0.05) +
  geom_segment(aes(y = HR_min[19]/1000000, yend = HR_max[19]/1000000, x = Dist_Rd[19], xend = Dist_Rd[19]),size = 1.5, colour = "#3471bc", alpha = 0.05) +
  geom_segment(aes(y = HR_min[20]/1000000, yend = HR_max[20]/1000000, x = Dist_Rd[20], xend = Dist_Rd[20]),size = 1.5, colour = "#3471bc", alpha = 0.05) +
  geom_segment(aes(y = HR_min[21]/1000000, yend = HR_max[21]/1000000, x = Dist_Rd[21], xend = Dist_Rd[21]),size = 1.5, colour = "#3471bc", alpha = 0.05) +
  
  #MS040
  geom_segment(aes(y = HR_min[22]/1000000, yend = HR_max[22]/1000000, x = Dist_Rd[22], xend = Dist_Rd[22]),size = 1.5, colour = "#3c7a47", alpha = 0.05) +
  geom_segment(aes(y = HR_min[23]/1000000, yend = HR_max[23]/1000000, x = Dist_Rd[23], xend = Dist_Rd[23]),size = 1.5, colour = "#3c7a47", alpha = 0.05) +
  geom_segment(aes(y = HR_min[24]/1000000, yend = HR_max[24]/1000000, x = Dist_Rd[24], xend = Dist_Rd[24]),size = 1.5, colour = "#3c7a47", alpha = 0.05) +
  geom_segment(aes(y = HR_min[25]/1000000, yend = HR_max[25]/1000000, x = Dist_Rd[25], xend = Dist_Rd[25]),size = 1.5, colour = "#3c7a47", alpha = 0.05) +
  geom_segment(aes(y = HR_min[26]/1000000, yend = HR_max[26]/1000000, x = Dist_Rd[26], xend = Dist_Rd[26]),size = 1.5, colour = "#3c7a47", alpha = 0.05) +
  geom_segment(aes(y = HR_min[27]/1000000, yend = HR_max[27]/1000000, x = Dist_Rd[27], xend = Dist_Rd[27]),size = 1.5, colour = "#3c7a47", alpha = 0.05) +
  geom_segment(aes(y = HR_min[28]/1000000, yend = HR_max[28]/1000000, x = Dist_Rd[28], xend = Dist_Rd[28]),size = 1.5, colour = "#3c7a47", alpha = 0.05) +
  geom_segment(aes(y = HR_min[29]/1000000, yend = HR_max[29]/1000000, x = Dist_Rd[29], xend = Dist_Rd[29]),size = 1.5, colour = "#3c7a47", alpha = 0.05) +
  geom_segment(aes(y = HR_min[30]/1000000, yend = HR_max[30]/1000000, x = Dist_Rd[30], xend = Dist_Rd[30]),size = 1.5, colour = "#3c7a47", alpha = 0.05) +
  geom_segment(aes(y = HR_min[31]/1000000, yend = HR_max[31]/1000000, x = Dist_Rd[31], xend = Dist_Rd[31]),size = 1.5, colour = "#3c7a47", alpha = 0.05) +
  geom_segment(aes(y = HR_min[32]/1000000, yend = HR_max[32]/1000000, x = Dist_Rd[32], xend = Dist_Rd[32]),size = 1.5, colour = "#3c7a47", alpha = 0.05) +
  geom_segment(aes(y = HR_min[33]/1000000, yend = HR_max[33]/1000000, x = Dist_Rd[33], xend = Dist_Rd[33]),size = 1.5, colour = "#3c7a47", alpha = 0.05) +
  geom_segment(aes(y = HR_min[34]/1000000, yend = HR_max[34]/1000000, x = Dist_Rd[34], xend = Dist_Rd[34]),size = 1.5, colour = "#3c7a47", alpha = 0.05) +
  geom_segment(aes(y = HR_min[35]/1000000, yend = HR_max[35]/1000000, x = Dist_Rd[35], xend = Dist_Rd[35]),size = 1.5, colour = "#3c7a47", alpha = 0.05) +
  geom_segment(aes(y = HR_min[36]/1000000, yend = HR_max[36]/1000000, x = Dist_Rd[36], xend = Dist_Rd[36]),size = 1.5, colour = "#3c7a47", alpha = 0.05) +
  geom_segment(aes(y = HR_min[37]/1000000, yend = HR_max[37]/1000000, x = Dist_Rd[37], xend = Dist_Rd[37]),size = 1.5, colour = "#3c7a47", alpha = 0.05) +
  geom_segment(aes(y = HR_min[38]/1000000, yend = HR_max[38]/1000000, x = Dist_Rd[38], xend = Dist_Rd[38]),size = 1.5, colour = "#3c7a47", alpha = 0.05) +
  
  geom_point(aes(y = HR/1000000, x = Dist_Rd), col = "white", size = 1) +
  geom_point(aes(y = HR/1000000, x = Dist_Rd, col = Road), size = 0.7) +
  
  scale_color_manual(labels = c("BR 267", "BR 262", "MS040"), values = c("#e6c141", "#3471bc", "#3c7a47")) +
  
  theme(panel.grid.major = element_blank(),
        panel.grid.minor = element_blank(),
        axis.ticks.y = element_blank(),
        axis.title.y = element_text(size=10, family = "serif"),
        axis.title.x = element_text(size=10, family = "serif"),
        axis.text.y = element_text(size=10, family = "serif"),
        axis.text.x = element_text(size=8, family = "serif"),
        plot.title = element_text(hjust = -0.025, size = 10, family = "serif"),
        legend.position="none") +
  ylab(expression(paste("Home-range area (km"^2,")"))) +
  xlab("Distance to road (km)") +
  scale_y_log10(breaks = c(1,2.5,5, 10,25, 50,100), limits = c(1,60), expand = c(0,0))


#Movement Speed vs. study site

Fid2_c <- 
  ggplot(data) +
  ggtitle("c)") +
  scale_y_discrete(limits=data$ID) +
  annotate("rect",
           xmin = -Inf, xmax = Inf,
           ymin = -Inf, ymax = 14.5, fill = "grey70",
           alpha = .15) +
  
  annotate("rect",
           xmin = -Inf, xmax = Inf,
           ymin = 14.5, ymax = 21.5, fill = "grey50",
           alpha = .15) +
  annotate("rect",
           xmin = -Inf, xmax = Inf,
           ymin = 21.5, ymax = Inf, fill = "grey20",
           alpha = .15) +
  
  theme_bw() +
  
  #BR_267
  geom_segment(aes(x = Speed_min[1]*86.4, xend = Speed_max[1]*86.4, y = 1, yend = 1),size = 1.5, colour = "#e6c141", alpha = 0.05) +
  geom_segment(aes(x = Speed_min[2]*86.4, xend = Speed_max[2]*86.4, y = 2, yend = 2),size = 1.5, colour = "#e6c141", alpha = 0.05) +
  geom_segment(aes(x = Speed_min[3]*86.4, xend = Speed_max[3]*86.4, y = 3, yend = 3),size = 1.5, colour = "#e6c141", alpha = 0.05) +
  geom_segment(aes(x = Speed_min[4]*86.4, xend = Speed_max[4]*86.4, y = 4, yend = 4),size = 1.5, colour = "#e6c141", alpha = 0.05) +
  geom_segment(aes(x = Speed_min[5]*86.4, xend = Speed_max[5]*86.4, y = 5, yend = 5),size = 1.5, colour = "#e6c141", alpha = 0.05) +
  geom_segment(aes(x = Speed_min[6]*86.4, xend = Speed_max[6]*86.4, y = 6, yend = 6),size = 1.5, colour = "#e6c141", alpha = 0.05) +
  geom_segment(aes(x = Speed_min[7]*86.4, xend = Speed_max[7]*86.4, y = 7, yend = 7),size = 1.5, colour = "#e6c141", alpha = 0.05) +
  geom_segment(aes(x = Speed_min[8]*86.4, xend = Speed_max[8]*86.4, y = 8, yend = 8),size = 1.5, colour = "#e6c141", alpha = 0.05) +
  geom_segment(aes(x = Speed_min[9]*86.4, xend = Speed_max[9]*86.4, y = 9, yend = 9),size = 1.5, colour = "#e6c141", alpha = 0.05) +
  geom_segment(aes(x = Speed_min[10]*86.4, xend = Speed_max[10]*86.4, y = 10, yend = 10),size = 1.5, colour = "#e6c141", alpha = 0.05) +
  geom_segment(aes(x = Speed_min[11]*86.4, xend = Speed_max[11]*86.4, y = 11, yend = 11),size = 1.5, colour = "#e6c141", alpha = 0.05) +
  geom_segment(aes(x = Speed_min[12]*86.4, xend = Speed_max[12]*86.4, y = 12, yend = 12),size = 1.5, colour = "#e6c141", alpha = 0.05) +
  geom_segment(aes(x = Speed_min[13]*86.4, xend = Speed_max[13]*86.4, y = 13, yend = 13),size = 1.5, colour = "#e6c141", alpha = 0.05) +
  geom_segment(aes(x = Speed_min[14]*86.4, xend = Speed_max[14]*86.4, y = 14, yend = 14),size = 1.5, colour = "#e6c141", alpha = 0.05) +
  
  #BR_262
  geom_segment(aes(x = Speed_min[15]*86.4, xend = Speed_max[15]*86.4, y = 15, yend = 15),size = 1.5, colour = "#3471bc", alpha = 0.05) +
  geom_segment(aes(x = Speed_min[16]*86.4, xend = Speed_max[16]*86.4, y = 16, yend = 16),size = 1.5, colour = "#3471bc", alpha = 0.05) +
  geom_segment(aes(x = Speed_min[17]*86.4, xend = Speed_max[17]*86.4, y = 17, yend = 17),size = 1.5, colour = "#3471bc", alpha = 0.05) +
  geom_segment(aes(x = Speed_min[18]*86.4, xend = Speed_max[18]*86.4, y = 18, yend = 18),size = 1.5, colour = "#3471bc", alpha = 0.05) +
  geom_segment(aes(x = Speed_min[19]*86.4, xend = Speed_max[19]*86.4, y = 19, yend = 19),size = 1.5, colour = "#3471bc", alpha = 0.05) +
  geom_segment(aes(x = Speed_min[20]*86.4, xend = Speed_max[20]*86.4, y = 20, yend = 20),size = 1.5, colour = "#3471bc", alpha = 0.05) +
  geom_segment(aes(x = Speed_min[21]*86.4, xend = Speed_max[21]*86.4, y = 21, yend = 21),size = 1.5, colour = "#3471bc", alpha = 0.05) +
  
  #MS040
  geom_segment(aes(x = Speed_min[22]*86.4, xend = Speed_max[22]*86.4, y = 22, yend = 22),size = 1.5, colour = "#3c7a47", alpha = 0.05) +
  geom_segment(aes(x = Speed_min[23]*86.4, xend = Speed_max[23]*86.4, y = 23, yend = 23),size = 1.5, colour = "#3c7a47", alpha = 0.05) +
  geom_segment(aes(x = Speed_min[24]*86.4, xend = Speed_max[24]*86.4, y = 24, yend = 24),size = 1.5, colour = "#3c7a47", alpha = 0.05) +
  geom_segment(aes(x = Speed_min[25]*86.4, xend = Speed_max[25]*86.4, y = 25, yend = 25),size = 1.5, colour = "#3c7a47", alpha = 0.05) +
  geom_segment(aes(x = Speed_min[26]*86.4, xend = Speed_max[26]*86.4, y = 26, yend = 26),size = 1.5, colour = "#3c7a47", alpha = 0.05) +
  geom_segment(aes(x = Speed_min[27]*86.4, xend = Speed_max[27]*86.4, y = 27, yend = 27),size = 1.5, colour = "#3c7a47", alpha = 0.05) +
  geom_segment(aes(x = Speed_min[28]*86.4, xend = Speed_max[28]*86.4, y = 28, yend = 28),size = 1.5, colour = "#3c7a47", alpha = 0.05) +
  geom_segment(aes(x = Speed_min[29]*86.4, xend = Speed_max[29]*86.4, y = 29, yend = 29),size = 1.5, colour = "#3c7a47", alpha = 0.05) +
  geom_segment(aes(x = Speed_min[30]*86.4, xend = Speed_max[30]*86.4, y = 30, yend = 30),size = 1.5, colour = "#3c7a47", alpha = 0.05) +
  geom_segment(aes(x = Speed_min[31]*86.4, xend = Speed_max[31]*86.4, y = 31, yend = 31),size = 1.5, colour = "#3c7a47", alpha = 0.05) +
  geom_segment(aes(x = Speed_min[32]*86.4, xend = Speed_max[32]*86.4, y = 32, yend = 32),size = 1.5, colour = "#3c7a47", alpha = 0.05) +
  geom_segment(aes(x = Speed_min[33]*86.4, xend = Speed_max[33]*86.4, y = 33, yend = 33),size = 1.5, colour = "#3c7a47", alpha = 0.05) +
  geom_segment(aes(x = Speed_min[34]*86.4, xend = Speed_max[34]*86.4, y = 34, yend = 34),size = 1.5, colour = "#3c7a47", alpha = 0.05) +
  geom_segment(aes(x = Speed_min[35]*86.4, xend = Speed_max[35]*86.4, y = 35, yend = 35),size = 1.5, colour = "#3c7a47", alpha = 0.05) +
  geom_segment(aes(x = Speed_min[36]*86.4, xend = Speed_max[36]*86.4, y = 36, yend = 36),size = 1.5, colour = "#3c7a47", alpha = 0.05) +
  geom_segment(aes(x = Speed_min[37]*86.4, xend = Speed_max[37]*86.4, y = 37, yend = 37),size = 1.5, colour = "#3c7a47", alpha = 0.05) +
  geom_segment(aes(x = Speed_min[38]*86.4, xend = Speed_max[38]*86.4, y = 38, yend = 38),size = 1.5, colour = "#3c7a47", alpha = 0.05) +
  geom_point(aes(x = Speed*86.4, y = ID), col = "white", size = 1) +
  geom_point(aes(x = Speed*86.4, y = ID, col = Road), size = 0.7) +
  
  scale_color_manual(labels = c("BR 267", "BR 262", "MS040"), values = c("#e6c141", "#3471bc", "#3c7a47")) +
  
  theme(panel.grid.major = element_blank(),
        panel.grid.minor = element_blank(),
        axis.ticks.y = element_blank(),
        axis.title.y = element_blank(),
        axis.title.x = element_text(size=10, family = "serif"),
        axis.text.y = element_text(size=5, family = "serif"),
        axis.text.x = element_text(size=8, family = "serif"),
        plot.title = element_text(hjust = -0.025, size = 10, family = "serif"),
        legend.position=c(0.08, 0.9), legend.title=element_blank(),
        legend.text=element_text(size=6, family = "serif"),legend.key.size = unit(0.2, "cm"),
        legend.key.width = unit(0.025, "cm"), legend.background=element_blank(), 
        legend.key = element_blank(), legend.text.align = 0) +
  xlab("Speed (km/day)") +
  scale_x_continuous(limits = c(5,10), expand = c(0,0))


#Movement speed with distance to road
Fid2_d <- 
  ggplot(data) +
  ggtitle("d)") +
  theme_bw() +
  
  #BR_267
  geom_segment(aes(y = Speed_min[1]*86.4, yend = Speed_max[1]*86.4, x = Dist_Rd[1], xend = Dist_Rd[1]),size = 1.5, colour = "#e6c141", alpha = 0.05) +
  geom_segment(aes(y = Speed_min[2]*86.4, yend = Speed_max[2]*86.4, x = Dist_Rd[2], xend = Dist_Rd[2]),size = 1.5, colour = "#e6c141", alpha = 0.05) +
  geom_segment(aes(y = Speed_min[3]*86.4, yend = Speed_max[3]*86.4, x = Dist_Rd[3], xend = Dist_Rd[3]),size = 1.5, colour = "#e6c141", alpha = 0.05) +
  geom_segment(aes(y = Speed_min[4]*86.4, yend = Speed_max[4]*86.4, x = Dist_Rd[4], xend = Dist_Rd[4]),size = 1.5, colour = "#e6c141", alpha = 0.05) +
  geom_segment(aes(y = Speed_min[5]*86.4, yend = Speed_max[5]*86.4, x = Dist_Rd[5], xend = Dist_Rd[5]),size = 1.5, colour = "#e6c141", alpha = 0.05) +
  geom_segment(aes(y = Speed_min[6]*86.4, yend = Speed_max[6]*86.4, x = Dist_Rd[6], xend = Dist_Rd[6]),size = 1.5, colour = "#e6c141", alpha = 0.05) +
  geom_segment(aes(y = Speed_min[7]*86.4, yend = Speed_max[7]*86.4, x = Dist_Rd[7], xend = Dist_Rd[7]),size = 1.5, colour = "#e6c141", alpha = 0.05) +
  geom_segment(aes(y = Speed_min[8]*86.4, yend = Speed_max[8]*86.4, x = Dist_Rd[8], xend = Dist_Rd[8]),size = 1.5, colour = "#e6c141", alpha = 0.05) +
  geom_segment(aes(y = Speed_min[9]*86.4, yend = Speed_max[9]*86.4, x = Dist_Rd[9], xend = Dist_Rd[9]),size = 1.5, colour = "#e6c141", alpha = 0.05) +
  geom_segment(aes(y = Speed_min[10]*86.4, yend = Speed_max[10]*86.4, x = Dist_Rd[10], xend = Dist_Rd[10]),size = 1.5, colour = "#e6c141", alpha = 0.05) +
  geom_segment(aes(y = Speed_min[11]*86.4, yend = Speed_max[11]*86.4, x = Dist_Rd[11], xend = Dist_Rd[11]),size = 1.5, colour = "#e6c141", alpha = 0.05) +
  geom_segment(aes(y = Speed_min[12]*86.4, yend = Speed_max[12]*86.4, x = Dist_Rd[12], xend = Dist_Rd[12]),size = 1.5, colour = "#e6c141", alpha = 0.05) +
  geom_segment(aes(y = Speed_min[13]*86.4, yend = Speed_max[13]*86.4, x = Dist_Rd[13], xend = Dist_Rd[13]),size = 1.5, colour = "#e6c141", alpha = 0.05) +
  geom_segment(aes(y = Speed_min[14]*86.4, yend = Speed_max[14]*86.4, x = Dist_Rd[14], xend = Dist_Rd[14]),size = 1.5, colour = "#e6c141", alpha = 0.05) +
  
  #BR_262
  geom_segment(aes(y = Speed_min[15]*86.4, yend = Speed_max[15]*86.4, x = Dist_Rd[15], xend = Dist_Rd[15]),size = 1.5, colour = "#3471bc", alpha = 0.05) +
  geom_segment(aes(y = Speed_min[16]*86.4, yend = Speed_max[16]*86.4, x = Dist_Rd[16], xend = Dist_Rd[16]),size = 1.5, colour = "#3471bc", alpha = 0.05) +
  geom_segment(aes(y = Speed_min[17]*86.4, yend = Speed_max[17]*86.4, x = Dist_Rd[17], xend = Dist_Rd[17]),size = 1.5, colour = "#3471bc", alpha = 0.05) +
  geom_segment(aes(y = Speed_min[18]*86.4, yend = Speed_max[18]*86.4, x = Dist_Rd[18], xend = Dist_Rd[18]),size = 1.5, colour = "#3471bc", alpha = 0.05) +
  geom_segment(aes(y = Speed_min[19]*86.4, yend = Speed_max[19]*86.4, x = Dist_Rd[19], xend = Dist_Rd[19]),size = 1.5, colour = "#3471bc", alpha = 0.05) +
  geom_segment(aes(y = Speed_min[20]*86.4, yend = Speed_max[20]*86.4, x = Dist_Rd[20], xend = Dist_Rd[20]),size = 1.5, colour = "#3471bc", alpha = 0.05) +
  geom_segment(aes(y = Speed_min[21]*86.4, yend = Speed_max[21]*86.4, x = Dist_Rd[21], xend = Dist_Rd[21]),size = 1.5, colour = "#3471bc", alpha = 0.05) +
  
  #MS040
  geom_segment(aes(y = Speed_min[22]*86.4, yend = Speed_max[22]*86.4, x = Dist_Rd[22], xend = Dist_Rd[22]),size = 1.5, colour = "#3c7a47", alpha = 0.05) +
  geom_segment(aes(y = Speed_min[23]*86.4, yend = Speed_max[23]*86.4, x = Dist_Rd[23], xend = Dist_Rd[23]),size = 1.5, colour = "#3c7a47", alpha = 0.05) +
  geom_segment(aes(y = Speed_min[24]*86.4, yend = Speed_max[24]*86.4, x = Dist_Rd[24], xend = Dist_Rd[24]),size = 1.5, colour = "#3c7a47", alpha = 0.05) +
  geom_segment(aes(y = Speed_min[25]*86.4, yend = Speed_max[25]*86.4, x = Dist_Rd[25], xend = Dist_Rd[25]),size = 1.5, colour = "#3c7a47", alpha = 0.05) +
  geom_segment(aes(y = Speed_min[26]*86.4, yend = Speed_max[26]*86.4, x = Dist_Rd[26], xend = Dist_Rd[26]),size = 1.5, colour = "#3c7a47", alpha = 0.05) +
  geom_segment(aes(y = Speed_min[27]*86.4, yend = Speed_max[27]*86.4, x = Dist_Rd[27], xend = Dist_Rd[27]),size = 1.5, colour = "#3c7a47", alpha = 0.05) +
  geom_segment(aes(y = Speed_min[28]*86.4, yend = Speed_max[28]*86.4, x = Dist_Rd[28], xend = Dist_Rd[28]),size = 1.5, colour = "#3c7a47", alpha = 0.05) +
  geom_segment(aes(y = Speed_min[29]*86.4, yend = Speed_max[29]*86.4, x = Dist_Rd[29], xend = Dist_Rd[29]),size = 1.5, colour = "#3c7a47", alpha = 0.05) +
  geom_segment(aes(y = Speed_min[30]*86.4, yend = Speed_max[30]*86.4, x = Dist_Rd[30], xend = Dist_Rd[30]),size = 1.5, colour = "#3c7a47", alpha = 0.05) +
  geom_segment(aes(y = Speed_min[31]*86.4, yend = Speed_max[31]*86.4, x = Dist_Rd[31], xend = Dist_Rd[31]),size = 1.5, colour = "#3c7a47", alpha = 0.05) +
  geom_segment(aes(y = Speed_min[32]*86.4, yend = Speed_max[32]*86.4, x = Dist_Rd[32], xend = Dist_Rd[32]),size = 1.5, colour = "#3c7a47", alpha = 0.05) +
  geom_segment(aes(y = Speed_min[33]*86.4, yend = Speed_max[33]*86.4, x = Dist_Rd[33], xend = Dist_Rd[33]),size = 1.5, colour = "#3c7a47", alpha = 0.05) +
  geom_segment(aes(y = Speed_min[34]*86.4, yend = Speed_max[34]*86.4, x = Dist_Rd[34], xend = Dist_Rd[34]),size = 1.5, colour = "#3c7a47", alpha = 0.05) +
  geom_segment(aes(y = Speed_min[35]*86.4, yend = Speed_max[35]*86.4, x = Dist_Rd[35], xend = Dist_Rd[35]),size = 1.5, colour = "#3c7a47", alpha = 0.05) +
  geom_segment(aes(y = Speed_min[36]*86.4, yend = Speed_max[36]*86.4, x = Dist_Rd[36], xend = Dist_Rd[36]),size = 1.5, colour = "#3c7a47", alpha = 0.05) +
  geom_segment(aes(y = Speed_min[37]*86.4, yend = Speed_max[37]*86.4, x = Dist_Rd[37], xend = Dist_Rd[37]),size = 1.5, colour = "#3c7a47", alpha = 0.05) +
  geom_segment(aes(y = Speed_min[38]*86.4, yend = Speed_max[38]*86.4, x = Dist_Rd[38], xend = Dist_Rd[38]),size = 1.5, colour = "#3c7a47", alpha = 0.05) +
  
  geom_point(aes(y = Speed*86.4, x = Dist_Rd), col = "white", size = 1) +
  geom_point(aes(y = Speed*86.4, x = Dist_Rd, col = Road), size = 0.7) +
  
  scale_color_manual(labels = c("BR 267", "BR 262", "MS040"), values = c("#e6c141", "#3471bc", "#3c7a47")) +
  
  theme(panel.grid.major = element_blank(),
        panel.grid.minor = element_blank(),
        axis.ticks.y = element_blank(),
        axis.title.y = element_text(size=10, family = "serif"),
        axis.title.x = element_text(size=10, family = "serif"),
        axis.text.y = element_text(size=10, family = "serif"),
        axis.text.x = element_text(size=8, family = "serif"),
        plot.title = element_text(hjust = -0.025, size = 10, family = "serif"),
        legend.position="none") +
  xlab("Distance to road (km)") +
  ylab("Speed (km/day)") +
  scale_y_continuous(limits = c(5,10), expand = c(0,0))


FIG <- grid.arrange(Fid2_a, Fid2_b,
                    Fid2_c, Fid2_d,
                    ncol = 2)
```

##### Instantaneous Speeds

Instantaneous speeds were analysed using a Gaussian mixed effects model, with a random intercept for each animal. We also included random slopes for the relationship between movement speeds and the distance to the road for each individual anteater. Finally, we applied a first order auto-regressive correction to the residuals to account for autocorrelation in instantaneous movement speeds.


\(\mathrm{Speed}\_i= \beta\_0 + \beta\_{\mathrm{Distance~to~road}~i} + \beta\_{\mathrm{Distance~to~road}~i}^2 + \beta\_{\mathrm{Time}~i} + \beta\_{\mathrm{Time}~i}^2 + \beta\_{\mathrm{Road}~i}\)

```
#Import the dataset of instantaneous Speeds
Speeds <- read.csv("Instantaneous_Speeds.csv")
#Drop infinite speeds
Speeds <- Speeds[!is.infinite(Speeds$est),]

#Drop animals that lived > 2 km from roads
Speeds <- Speeds[which(Speeds$ID != "Alexander"),]
Speeds <- Speeds[which(Speeds$ID != "Bumpus"),]
Speeds <- Speeds[which(Speeds$ID != "Jackson"),]
Speeds <- Speeds[which(Speeds$ID != "Kyle"),]
Speeds <- Speeds[which(Speeds$ID != "Little Rick"),]
Speeds <- Speeds[which(Speeds$ID != "Puji"),]
Speeds <- Speeds[which(Speeds$ID != "Makao"),]
Speeds <- Speeds[which(Speeds$ID != "Hannah"),]
Speeds <- Speeds[which(Speeds$ID != "Yoki"),]
Speeds <- Speeds[which(Speeds$ID != "Phoenix 1 error"),]
Speeds <- Speeds[which(Speeds$ID != "Delphine"),]
Speeds <- Speeds[which(Speeds$ID != "Gala"),]

#Convert timestamps
Speeds$timestamp <- as.POSIXct(Speeds$timestamp)

#Add one-hour cut points for aggregating
Speeds$time <- cut(Speeds$timestamp, breaks = "60 min")

#Counts number of crossings per hour
SPEEDS <- aggregate(cbind(est, Dist) ~ time + ID, data = Speeds, FUN = "mean")
ROAD <- aggregate(cbind(Road) ~ time + ID, data = Speeds, FUN = "unique")
SPEEDS$Road <- as.factor(ROAD$Road)
SPEEDS$time <- as.POSIXct(SPEEDS$time)
SPEEDS$hour <- lubridate::hour(SPEEDS$time)
SPEEDS$Dist <- log(SPEEDS$Dist)

#FIT <- lme(est ~ Dist + I(Dist^2) + hour + I(hour^2) + Road,
#           random = ~ Dist|ID,
#           correlation = corAR1(form = ~ 1|ID),
#           data = SPEEDS) 

load("Speeds_AR1_Fit.Rda")
  
acf(residuals(FIT, type = "normalized"), main = "")
```

```
summary(FIT)
```

```
## Linear mixed-effects model fit by REML
##   Data: SPEEDS 
##         AIC       BIC   logLik
##   -853248.2 -853125.8 426636.1
## 
## Random effects:
##  Formula: ~Dist | ID
##  Structure: General positive-definite, Log-Cholesky parametrization
##             StdDev      Corr  
## (Intercept) 0.013623473 (Intr)
## Dist        0.003577603 -0.455
## Residual    0.032922717       
## 
## Correlation Structure: AR(1)
##  Formula: ~1 | ID 
##  Parameter estimate(s):
##       Phi 
## 0.5139856 
## Fixed effects:  est ~ Dist + I(Dist^2) + hour + I(hour^2) + Road 
##                   Value   Std.Error     DF   t-value p-value
## (Intercept)  0.10085140 0.003594504 198661  28.05711  0.0000
## Dist         0.00183677 0.000753514 198661   2.43760  0.0148
## I(Dist^2)   -0.00022820 0.000081888 198661  -2.78671  0.0053
## hour        -0.00500373 0.000064934 198661 -77.05829  0.0000
## I(hour^2)    0.00020876 0.000002764 198661  75.52137  0.0000
## RoadBR262    0.01051875 0.005734860     26   1.83418  0.0781
## RoadMS-040   0.00309344 0.005288111     26   0.58498  0.5636
##  Correlation: 
##            (Intr) Dist   I(D^2) hour   I(h^2) RBR262
## Dist       -0.321                                   
## I(Dist^2)   0.045 -0.313                            
## hour       -0.072  0.003  0.001                     
## I(hour^2)   0.065 -0.004 -0.001 -0.979              
## RoadBR262  -0.558  0.005  0.000  0.000  0.000       
## RoadMS-040 -0.601 -0.007 -0.006 -0.002  0.002  0.378
## 
## Standardized Within-Group Residuals:
##         Min          Q1         Med          Q3         Max 
## -1.84685734 -0.50083311 -0.26144699  0.09778769 15.45225445 
## 
## Number of Observations: 198694
## Number of Groups: 29
```

```
IDs <- aggregate(Road ~ ID, data = Speeds, FUN = "unique")

Preds <- list()
for(i in 1:nrow(IDs)){
  
  DISTS <- log(seq(0,6, length.out = 500))
  
  NewData <- data.frame(Dist = rep(DISTS,3),
                        hour = 0,
                        ID = IDs$ID[i],
                        Road = sort(factor(rep(levels(SPEEDS$Road),500*3))))
  
  Preds_i <- predict(FIT, newdata = NewData)
  Preds[[i]] <- data.frame(ID = IDs$ID[i],
                           Speed = Preds_i[which(sort(factor(rep(levels(SPEEDS$Road),500*3))) == IDs$Road[i])],
                           Dist = DISTS,
                           Road = IDs$Road[i])
}


Preds <- do.call(rbind, Preds)

ggplot(Preds) +
  geom_smooth(aes(x = Dist, y = Speed, col = ID)) +
  scale_color_viridis_d() +
  theme_bw() +
  theme(panel.grid.major = element_blank(),
        panel.grid.minor = element_blank(),
        axis.title.y = element_text(size=10, family = "serif"),
        axis.title.x = element_text(size=10, family = "serif"),
        axis.text.y = element_text(size=8, family = "serif"),
        axis.text.x = element_text(size=8, family = "serif"),
        plot.title = element_text(hjust = -0.025, size = 10, family = "serif"),
        legend.position="none", legend.title=element_blank(),
        legend.text=element_text(size=6, family = "serif"),legend.key.size = unit(0.2, "cm"),
        legend.key.width = unit(0.25, "cm"), legend.background=element_blank(), 
        legend.key = element_blank(), legend.text.align = 0) +
  ylab(expression(paste("Speed (m/s)"))) +
  xlab(expression(paste("Distance from road (km)"))) +
  scale_y_continuous(limits = c(0,.2), expand = c(0,0)) +
  scale_x_continuous(breaks = log(c(0.025, 0.05, .1, .2, .4, .8, 1.6, 3.2)),
                     labels = c(0.025, 0.05, .1, .2, .4, .8, 1.6, 3.2),
                     expand = c(0,0))
```

---

#### Question 2: Does traffic volume influence the movement or crossing behaviour?

The second question we addressed was focused on understanding the factors the influenced individual giant anteaters willingness to cross roads. In addition to calculating basic summary statistics,we modelled the relationship between the number of road crossings according to the nearest highway as a proxy for traffic volume. We also included the variables related to the home range location (distance of home range centroid to nearest paved road), sampling duration, and animal traits (sex and body mass). The road crossing data were zero-inflated, where only 26 of the 38 animals were actually observed to have crossed a road. As a result, we modelled these data using a hurdle model that modelled individual crossing/not crossing information according to a logistic regression model, and the individual number of crossings for those animals that did cross according to a zero-truncated negative binomial generalised linear model (GLM). This formulation allowed us to distinguish between the factors governing whether giant anteaters will cross roads or not, and the factors driving crossing rate for those animals that do cross roads. In essence, this allowed us to capture both ecological process in a single model. For both processes, we started with the following global model:


\(\mu\_i= \beta\_0 + \beta\_{\mathrm{Distance~to~road}~i} + \beta\_{\mathrm{Sex}~i} + \beta\_{\mathrm{Weight}~i} + \beta\_{\mathrm{\%~Pasture}~i} + \beta\_{\mathrm{Road}~i} + \beta\_{\mathrm{Sampling~Duration}~i}\)

From this global model, we specified a subset of candidate models comprising of all possible combinations of fixed effects using the R package ‘MuMIn’ (Bartoń 2016), and used the AICc for model selection to identify the best-fit model for the data. As AICc has been shown to under/overfit models on small sample sizes (Brewer et al. 2016 ), we confirmed our selected model via block cross-validation using the methods implemented in the R package DAAG (Maindonald and Braun 2015 ).

```
#Subset the animals that lived close to paved roads
data_subset <- data[data$Dist_Rd<2,]

#How many animals crossed the road
length(which(data_subset$Rd_Cr > 0))
```

```
## [1] 22
```

```
#Median crossings per animal
median(data_subset$Rd_Cr)
```

```
## [1] 14
```

```
#95% CIs on the median
n <- length(na.omit(data_subset$Rd_Cr))
sort(na.omit(data_subset$Rd_Cr))[round((n/2)*(1 + (1.96)/sqrt(n)),0)]
```

```
## [1] 48
```

```
sort(na.omit(data_subset$Rd_Cr))[round((n/2)*(1 - (1.96)/sqrt(n)),0)]
```

```
## [1] 2
```

```
#Histogram showing the zero inflation
hist(data$Rd_Cr, main = "Total road crossings for individual giant anteaters")
```

```
#Create a variable describing crossers vs non-crossers
data$Cross <- 0
data[which(data$Rd_Cr > 0), "Cross"] <- 1

#Model selection on the logistic part of the model
mod <- glm(Cross ~ 1 + Road + Dist_Rd + Duration + Sex + Weight + Pasture,
           family = binomial,
           data = data,
           na.action = na.fail)

#Model Selection (shows all are worth considering)
res <- dredge(mod)
head(res, 10)
```

```
## Global model call: glm(formula = Cross ~ 1 + Road + Dist_Rd + Duration + Sex + Weight + 
##     Pasture, family = binomial, data = data, na.action = na.fail)
## ---
## Model selection table 
##      (Int)  Dst_Rd      Drt      Pst Rod Sex      Wgh df  logLik AICc delta
## 64 923.500 -23.460 -0.20610 -2.93600   +   + -15.7000  8  -2.469 25.9  0.00
## 62 119.300  -3.617          -0.38800   +   +  -2.1770  7  -4.998 27.7  1.83
## 20   9.901  -1.470 -0.02370                +           4  -9.572 28.4  2.45
## 24  18.790  -1.591 -0.03178 -0.06969       +           5  -8.696 29.3  3.37
## 52  17.630  -1.696 -0.02572                +  -0.2073  5  -8.937 29.7  3.85
## 56  25.420  -1.750 -0.03546 -0.06458       +  -0.1739  6  -8.305 31.3  5.42
## 18   2.054  -1.576                         +           3 -12.549 31.8  5.90
## 58  25.360  -1.808                     +   +  -0.6277  6  -8.558 31.8  5.92
## 32  21.920  -1.500 -0.02172 -0.13710   +   +           7  -7.520 32.8  6.87
## 28   8.833  -1.407 -0.02129            +   +           6  -9.131 33.0  7.07
##    weight
## 64  0.443
## 62  0.178
## 20  0.130
## 24  0.082
## 52  0.065
## 56  0.030
## 18  0.023
## 58  0.023
## 32  0.014
## 28  0.013
## Models ranked by AICc(x)
```

```
# Cross validation to asses parameter importance
# After multiple rounds of parameter reduction, the following model was obtained
# This had the highest CV performance
Cross_mod <- glm(Cross ~ 1 + Road + Dist_Rd  + Sex,
                 family = binomial,
                 data = data,
                 na.action = na.fail)

CV_Res <- vector("numeric", 400)
for(i in 1:400){
  CV_Res[i] <- CVbinary(Cross_mod, print.details = FALSE)$acc.cv
}

#Mean cross validation rate
mean(CV_Res)
```

```
## [1] 0.8415789
```

```
t.test(CV_Res)
```

```
## 
##  One Sample t-test
## 
## data:  CV_Res
## t = 588.16, df = 399, p-value < 2.2e-16
## alternative hypothesis: true mean is not equal to 0
## 95 percent confidence interval:
##  0.8387660 0.8443919
## sample estimates:
## mean of x 
## 0.8415789
```

```
# Model selection to identify the parameters to include in
# the zero-truncated count portion of the model
mod <- pscl::hurdle(Rd_Cr ~ 1 + Road + Dist_Rd + Duration + Sex + Weight + Pasture | 1,
                    data = data,
                    na.action = na.fail,
                    dist = "negbin")

# Model selection on the count part of the model
res <- dredge(mod)
head(res, 10)
```

```
## Global model call: pscl::hurdle(formula = Rd_Cr ~ 1 + Road + Dist_Rd + Duration + 
##     Sex + Weight + Pasture | 1, data = data, na.action = na.fail, 
##     dist = "negbin")
## ---
## Model selection table 
##    cnt_(Int) cnt_Dst_Rd  cnt_Drt cnt_Pst cnt_Rod cnt_Sex  cnt_Wgh zer_(Int) df
## 22     16.08     -2.513          -0.1277               +             0.6539  6
## 14     19.89     -2.453          -0.1698       +                     0.6539  7
## 30     17.02     -2.391          -0.1439       +       +             0.6539  8
## 6      18.80     -2.565          -0.1462                             0.6539  5
## 54     17.28     -2.474          -0.1230               + -0.05341    0.6539  7
## 24     14.88     -2.551 0.001582 -0.1189               +             0.6539  7
## 46     18.00     -2.445          -0.1766       +          0.07459    0.6539  8
## 16     18.52     -2.548 0.002573 -0.1611       +                     0.6539  8
## 32     14.86     -2.530 0.003634 -0.1302       +       +             0.6539  9
## 62     15.92     -2.388          -0.1500       +       +  0.05022    0.6539  9
##      logLik  AICc delta weight
## 22 -133.017 280.7  0.00  0.236
## 14 -131.568 280.9  0.13  0.222
## 30 -130.450 281.9  1.12  0.135
## 6  -135.254 282.4  1.64  0.104
## 54 -132.851 283.4  2.69  0.061
## 24 -132.904 283.5  2.80  0.058
## 46 -131.303 283.6  2.83  0.057
## 16 -131.335 283.6  2.89  0.056
## 32 -129.872 284.2  3.43  0.043
## 62 -130.306 285.0  4.30  0.028
## Models ranked by AICc(x)
```

```
#Build the complete, selected zero-altered model
Selected_Mod <- pscl::hurdle(Rd_Cr ~ Dist_Rd + Sex + Pasture | Road + Dist_Rd + Sex,
                             data = data,
                             dist = "negbin")

summary(Selected_Mod)
```

```
## 
## Call:
## pscl::hurdle(formula = Rd_Cr ~ Dist_Rd + Sex + Pasture | Road + Dist_Rd + 
##     Sex, data = data, dist = "negbin")
## 
## Pearson residuals:
##      Min       1Q   Median       3Q      Max 
## -0.86623 -0.69306 -0.23555  0.05684  5.29722 
## 
## Count model coefficients (truncated negbin with log link):
##             Estimate Std. Error z value Pr(>|z|)    
## (Intercept) 16.07823    3.50103   4.592 4.38e-06 ***
## Dist_Rd     -2.51277    0.52903  -4.750 2.04e-06 ***
## SexMale      1.33153    0.54493   2.444  0.01455 *  
## Pasture     -0.12766    0.03644  -3.503  0.00046 ***
## Log(theta)  -0.17564    0.38368  -0.458  0.64711    
## Zero hurdle model coefficients (binomial with logit link):
##              Estimate Std. Error z value Pr(>|z|)  
## (Intercept)    1.9672     0.9709   2.026   0.0428 *
## RoadBR262     17.0075  5794.0460   0.003   0.9977  
## RoadMS-040    -1.6624     1.3938  -1.193   0.2330  
## Dist_Rd       -1.2193     0.5626  -2.167   0.0302 *
## SexMale        3.6635     1.6777   2.184   0.0290 *
## ---
## Signif. codes:  0 '***' 0.001 '**' 0.01 '*' 0.05 '.' 0.1 ' ' 1 
## 
## Theta: count = 0.8389
## Number of iterations in BFGS optimization: 22 
## Log-likelihood: -119.5 on 10 Df
```

```
par(mfrow = c(1,2))
#Figure of the estimated model coefficients
plot(y = coef(summary(Selected_Mod))$count[1:5,1],
     x = 1:5,
     axes = F,
     pch = 16,
     ylim = c(-5,24),
     xlab = "",
     ylab = expression(hat(beta)),
     cex = 0.7, mgp=c(2,1,0),
     main = "Param Est for Number Road Crossings")
text(x =  1:5, y = par("usr")[3]-0.05, srt = 40, adj = 1,
     labels = row.names(coef(summary(Selected_Mod))$count)[1:5], xpd = TRUE, cex = 0.7)
segments(x0 = 1:5, x1 = 1:5, y0 = confint(Selected_Mod)[1:5,1], y1 = confint(Selected_Mod)[1:5,2])
axis(2)
abline(h = 0, col = "grey", lty = 2)

plot(y = coef(summary(Selected_Mod))$zero[1:5,1],
     x = 1:5,
     axes = F,
     pch = 16,
     ylim = c(-3,7),
     xlab = "",
     ylab = expression(hat(beta)),
     cex = 0.7, mgp=c(2,1,0),
     main = "Param Est for Crossing or Not")
text(x =  1:5, y = par("usr")[3]-0.05, srt = 40, adj = 1,
     labels = row.names(coef(summary(Selected_Mod))$zero), xpd = TRUE, cex = 0.7)
segments(x0 = 1:5, x1 = 1:5, y0 = confint(Selected_Mod)[5:9,1], y1 = confint(Selected_Mod)[5:9,2])
axis(2)
abline(h = 0, col = "grey", lty = 2)
```

##### Recreate Figure 3

```
#Add a negative binomial curve for plotting purposes
NEW_DATA <- data.frame(Dist_Rd = seq(0, 6, 0.01))
mod1 <- MASS::glm.nb(Rd_Cr ~ Dist_Rd, data = data)
PRED <- predict(mod1, newdata = NEW_DATA, type = "link", se = T)

#Plot the confidence intervals
g1 <- 
  ggplot() +
  geom_line(aes(x=NEW_DATA$Dist, y=exp(PRED$fit - 1.96*PRED$se.fit))) +
  geom_line(aes(x=NEW_DATA$Dist, y=exp(PRED$fit + 1.96*PRED$se.fit)))

gg1 <- ggplot_build(g1)

# extract data for the loess lines from the 'data' slot
VAR <- data.frame(x = gg1$data[[1]]$x,
                  ymin = gg1$data[[1]]$y,
                  ymax = gg1$data[[2]]$y) 

Dist <- 
  ggplot() +
  ggtitle("a)") +
  geom_line(aes(y = exp(PRED$fit), x =  NEW_DATA$Dist), col = "grey70") +
  geom_ribbon(data = VAR, aes(x = x, ymin = ymin, ymax = ymax), fill="grey70", alpha = 0.2) +
  geom_point(data = data, aes(x = Dist_Rd, y = Rd_Cr, col = Road), size = 0.5) +
  scale_color_manual(labels=c("BR 267", "BR 262", "MS040"), values = c("#e6c141", "#3471bc", "#3c7a47")) +
  theme_bw() +
  theme(panel.grid.major = element_blank(),
        panel.grid.minor = element_blank(),
        axis.title.y = element_text(size=10, family = "serif"),
        axis.title.x = element_text(size=10, family = "serif"),
        axis.text.y = element_text(size=8, family = "serif"),
        axis.text.x = element_text(size=8, family = "serif"),
        plot.title = element_text(hjust = -0.025, size = 10, family = "serif"),
        legend.position=c(0.8, 0.9), legend.title=element_blank(),
        legend.text=element_text(size=6, family = "serif"),legend.key.size = unit(0.2, "cm"),
        legend.key.width = unit(0.025, "cm"), legend.background=element_blank(), 
        legend.key = element_blank(), legend.text.align = 0) +
  xlab(expression(paste("Distance to road (km)"))) +
  ylab(expression(paste("Number of road crossings"))) +
  coord_cartesian(ylim = c(-3,300), expand = c(0,0))

Sex <- 
  ggplot(data) +
  ggtitle("b)") +
  geom_boxplot(aes(x = Sex, y = Rd_Cr, fill = Road), size = 0.1, outlier.size=0.5) +
  scale_fill_manual(labels=c("BR 267", "BR 262", "MS040"), values = c("#e6c141", "#3471bc", "#3c7a47")) +
  theme_bw() +
  theme(panel.grid.major = element_blank(),
        panel.grid.minor = element_blank(),
        axis.title.y = element_text(size=10, family = "serif"),
        axis.title.x = element_text(size=10, family = "serif"),
        axis.text.y = element_text(size=8, family = "serif"),
        axis.text.x = element_text(size=8, family = "serif"),
        plot.title = element_text(hjust = -0.025, size = 10, family = "serif"),
        legend.position=c(0.15, 0.9), legend.title=element_blank(),
        legend.text=element_text(size=6, family = "serif"),legend.key.size = unit(0.2, "cm"),
        legend.key.width = unit(0.25, "cm"), legend.background=element_blank(), 
        legend.key = element_blank(), legend.text.align = 0) +
  xlab(expression(paste("Sex"))) +
  ylab(expression(paste("Number of road crossings")))


NEW_DATA2 <- data.frame(Pasture = seq(50, 100, 0.01))
mod2 <- MASS::glm.nb(Rd_Cr ~ Pasture, data = data[which(data$Rd_Cr > 0),])
PRED2 <- predict(mod2, newdata = NEW_DATA2, type = "link", se = T)

#Plot the confidence intervals
g2 <- 
  ggplot() +
  geom_line(aes(x=NEW_DATA2$Pasture, y=exp(PRED2$fit - 1.96*PRED2$se.fit))) +
  geom_line(aes(x=NEW_DATA2$Pasture, y=exp(PRED2$fit + 1.96*PRED2$se.fit)))

gg2 <- ggplot_build(g2)

# extract data for the loess lines from the 'data' slot
VAR2 <- data.frame(x = gg2$data[[1]]$x,
                  ymin = gg2$data[[1]]$y,
                  ymax = gg2$data[[2]]$y) 

Pasture <- 
  ggplot() +
  ggtitle("c)") +
  geom_line(aes(y = exp(PRED2$fit), x =  NEW_DATA2$Pasture), col = "grey70") +
  geom_ribbon(data = VAR2, aes(x = x, ymin = ymin, ymax = ymax), fill="grey70", alpha = 0.2) +
  geom_point(data = data, aes(x = Pasture, y = Rd_Cr, col = Road), size = 0.5) +
  scale_color_manual(labels=c("BR 267", "BR 262", "MS040"), values = c("#e6c141", "#3471bc", "#3c7a47")) +
  theme_bw() +
  theme(panel.grid.major = element_blank(),
        panel.grid.minor = element_blank(),
        axis.title.y = element_text(size=10, family = "serif"),
        axis.title.x = element_text(size=10, family = "serif"),
        axis.text.y = element_text(size=8, family = "serif"),
        axis.text.x = element_text(size=8, family = "serif"),
        plot.title = element_text(hjust = -0.025, size = 10, family = "serif"),
        legend.position="none", legend.title=element_blank(),
        legend.text=element_text(size=6, family = "serif"),legend.key.size = unit(0.2, "cm"),
        legend.key.width = unit(0.025, "cm"), legend.background=element_blank(), 
        legend.key = element_blank(), legend.text.align = 0) +
  xlab(expression(paste("Proportion of Pasture Land"))) +
  ylab(expression(paste("Number of road crossings"))) +
  coord_cartesian(ylim = c(-10,300), expand = c(0,10))

FIGS <- gridExtra::grid.arrange(Dist, Sex, Pasture, ncol = 1)
```

#### Crossing Time Analyses

##### Recreate Figure 4

```
#####################################################
# Pannel a) - Crossing times and traffic volume

#Import and clean up the data
traffic <- read.csv("Traffic.csv")
CROSS_TIMES <- read.csv("Crossing_Times.csv")
#Drop NAs
CROSS_TIMES <- CROSS_TIMES[!is.na(CROSS_TIMES$Crossing_Times),]
#Convert timestamps
CROSS_TIMES$Crossing_Times <- as.POSIXct(CROSS_TIMES$Crossing_Times)


#Obtain median/range of crossings per animal per day
CROSS_TIMES$day <- cut(CROSS_TIMES$Crossing_Times, breaks = "day")
CROSSES_DAY <- aggregate(Crossing_Times ~ day + ID, data = CROSS_TIMES, FUN = "length")
median(CROSSES_DAY$Crossing_Times)
```

```
## [1] 2
```

```
range(CROSSES_DAY$Crossing_Times)
```

```
## [1]  1 15
```

```
#Add one-hour cut points for aggregating
CROSS_TIMES$time <- cut(CROSS_TIMES$Crossing_Times, breaks = "hour")
#Counts number of crossings per hour
crossings <- aggregate(ID ~ time, data = CROSS_TIMES, FUN = "length")
#Generate a sequence of times to fill in with points when no crossings were observed
crossings$time <- as_datetime(crossings$time)

TIMES <- seq(from = min(crossings$time),
             to = max(crossings$time),
             by = "hour")

TIMES <- data.frame(time = TIMES,
                    crossings = 0)

#Merge the crossing events with the non-crossing events
crossings_2 <- merge(TIMES, crossings, all = TRUE)

#Change NAs to 0
crossings_2$ID[is.na(crossings_2$ID)] <- 0

#Reshuffle/restructure
crossings_2$crossings <- crossings_2$ID
crossings_2$ID <- NULL
crossings_2$hour <- hour(crossings_2$time)

#get the mean number of crossings per hour
crossings <- aggregate(crossings ~ hour, data = crossings_2, FUN = "mean")

#get the standard error of number of crossings per hour
se <- function(x) sqrt(var(x)/length(x))
crossings_se <- aggregate(crossings ~ hour, data = crossings_2, FUN = "se")
crossings$se <- crossings_se$crossings

#calculate 95% CIs
crossings$max <- crossings$crossings + 1.96*crossings$se
crossings$min <- crossings$crossings - 1.96*crossings$se

#########################################################
#Statistical test on crossings vs traffic volume
mod <- lm(crossings$crossings ~ traffic$Cars[-25])
summary(mod)
```

```
## 
## Call:
## lm(formula = crossings$crossings ~ traffic$Cars[-25])
## 
## Residuals:
##       Min        1Q    Median        3Q       Max 
## -0.025954 -0.008567 -0.003730  0.012963  0.024049 
## 
## Coefficients:
##                     Estimate Std. Error t value Pr(>|t|)    
## (Intercept)        8.635e-02  6.156e-03  14.026 1.88e-12 ***
## traffic$Cars[-25] -5.310e-05  1.287e-05  -4.127 0.000442 ***
## ---
## Signif. codes:  0 '***' 0.001 '**' 0.01 '*' 0.05 '.' 0.1 ' ' 1
## 
## Residual standard error: 0.01573 on 22 degrees of freedom
## Multiple R-squared:  0.4364, Adjusted R-squared:  0.4107 
## F-statistic: 17.03 on 1 and 22 DF,  p-value: 0.0004424
```

```
#########################################################

crossings2 <- crossings
crossings2$hour <- crossings2$hour - 24

crossings3 <- crossings
crossings3$hour <- crossings3$hour + 24

crossings <- rbind(crossings2, crossings, crossings3) 

g1 <- ggplot(data=crossings) +
  stat_smooth(aes(x=hour, y=max), colour="steelblue2", span = 0.3) +
  stat_smooth(aes(x=hour, y=min), colour="steelblue2", span = 0.3)
gg1 <- ggplot_build(g1)

# extract data for the loess lines from the 'data' slot
CIs <- data.frame(x = gg1$data[[1]]$x,
                  ymin = gg1$data[[1]]$y,
                  ymax = gg1$data[[2]]$y) 

#Axis labels
LABELS <- seq(from=as.POSIXct("2020-1-1 0:00"), to=as.POSIXct("2020-1-1 24:00"), by="2 hours")
LABELS <- format(strptime(LABELS,"%Y-%m-%d %H:%M:%S"),'%H:%M')
LABELS[13] <- "24:00"

#Scale the traffic data
traffic$Cars_2 <- (traffic$Cars/max(traffic$Cars))*0.1

#Plot the figure
crossings_pannel <- 
  ggplot(data=crossings) +
  ggtitle("a)") +
  geom_bar(data = traffic, aes(x=Hour, y=Cars_2), stat="identity", alpha = 0.5, fill = "#f8da78") + 
  geom_point(aes(x=hour, y=crossings), col = "grey70", size = 0.35) +
  stat_smooth(aes(x=hour, y=crossings), se=FALSE, lwd = 0.5, col = "#046C9A", span = 0.3) +
  geom_ribbon(data = CIs, aes(x = x, ymin = ymin, ymax = ymax), fill="#046C9A", alpha = 0.2) +
  theme_bw() +
  theme(panel.grid.major = element_blank(),
        panel.grid.minor = element_blank(),
        axis.title.y = element_text(size=10, family = "serif"),
        axis.title.x = element_text(size=10, family = "serif"),
        axis.text.y = element_text(size=8, family = "serif"),
        axis.text.x = element_text(size=5, family = "serif"),
        plot.title = element_text(hjust = -0.025, size = 10, family = "serif"),
        legend.position=c(0.15, 0.9), legend.title=element_blank(),
        legend.text=element_text(size=6, family = "serif"),legend.key.size = unit(0.2, "cm"),
        legend.key.width = unit(0.25, "cm"), legend.background=element_blank(), 
        legend.key = element_blank(), legend.text.align = 0) +
  ylab(expression(paste("Mean road crossings"))) +
  xlab(expression(paste("Time of day"))) +
  scale_x_continuous(breaks = seq(0,24,2), labels = LABELS, expand = c(0, 0.05)) +
  scale_y_continuous(expand = c(0, 0.0001),
                     sec.axis = sec_axis(~. * 6666.667, name = "Traffic Volume")) +
  coord_cartesian(xlim = c(-.3, 24.3), ylim = c(0,0.115))


#####################################################
# Pannel b) - circadian activity rhythm

#Import the dataset of instantaneous Speeds
Speeds <- read.csv("Instantaneous_Speeds.csv")
#Drop infinite speeds
Speeds <- Speeds[!is.infinite(Speeds$est),]
#Convert timestamps
Speeds$timestamp <- as.POSIXct(Speeds$timestamp)

#Add one-hour cut points for aggregating
Speeds$time <- cut(Speeds$timestamp, breaks = "hour")
Speeds$hour <- hour(Speeds$time)

#Counts number of crossings per hour
SPEEDS <- aggregate(est ~ hour, data = Speeds, FUN = "mean")

#get the standard error of number of crossings per hour
se <- function(x) sqrt(var(x)/length(x))
SPEEDS_se <- aggregate(est ~ hour, data = Speeds, FUN = "se")
SPEEDS$se <- SPEEDS_se$est

#calculate 95% CIs
SPEEDS$max <- SPEEDS$est + 1.96*SPEEDS$se
SPEEDS$min <- SPEEDS$est - 1.96*SPEEDS$se

#Make the data span 3 "days" so the 24hr plot starts and ends at the right spots
SPEEDS2 <- SPEEDS
SPEEDS2$hour <- SPEEDS2$hour - 24
SPEEDS3 <- SPEEDS
SPEEDS3$hour <- SPEEDS3$hour + 24

SPEEDS <- rbind(SPEEDS2, SPEEDS, SPEEDS3) 

g1 <- ggplot(data=SPEEDS) +
  stat_smooth(aes(x=hour, y=max), colour="steelblue2", span = 0.3) +
  stat_smooth(aes(x=hour, y=min), colour="steelblue2", span = 0.3)
gg1 <- ggplot_build(g1)

# extract data for the loess lines from the 'data' slot
CIs <- data.frame(x = gg1$data[[1]]$x,
                  ymin = gg1$data[[1]]$y,
                  ymax = gg1$data[[2]]$y) 

#Axis labels
LABELS <- seq(from=as.POSIXct("2020-1-1 0:00"), to=as.POSIXct("2020-1-1 24:00"), by="2 hours")
LABELS <- format(strptime(LABELS,"%Y-%m-%d %H:%M:%S"),'%H:%M')
LABELS[13] <- "24:00"

#Plot the figure
Activity_pannel <- 
  ggplot(data=SPEEDS) +
  ggtitle("b)") +
  geom_point(aes(x=hour, y=est), col = "grey70", size = 0.35) +
  stat_smooth(aes(x=hour, y=est), se=FALSE, lwd = 0.5, col = "#046C9A", span = 0.3) +
  geom_ribbon(data = CIs, aes(x = x, ymin = ymin, ymax = ymax), fill="#046C9A", alpha = 0.2) +
  theme_bw() +
  theme(panel.grid.major = element_blank(),
        panel.grid.minor = element_blank(),
        axis.title.y = element_text(size=10, family = "serif"),
        axis.title.x = element_text(size=10, family = "serif"),
        axis.text.y = element_text(size=8, family = "serif"),
        axis.text.x = element_text(size=5, family = "serif"),
        plot.title = element_text(hjust = -0.025, size = 10, family = "serif"),
        legend.position=c(0.15, 0.9), legend.title=element_blank(),
        legend.text=element_text(size=6, family = "serif"),legend.key.size = unit(0.2, "cm"),
        legend.key.width = unit(0.25, "cm"), legend.background=element_blank(), 
        legend.key = element_blank(), legend.text.align = 0) +
  ylab(expression(paste("Mean speed (m/s)"))) +
  xlab(expression(paste("Time of day"))) +
  scale_x_continuous(breaks = seq(0,24,2), labels = LABELS, expand = c(0, 0.05)) +
  scale_y_continuous(expand = c(0, 0.0001)) +
  coord_cartesian(xlim = c(-.3, 24.3), ylim = c(0.068,0.1))


gridExtra::grid.arrange(crossings_pannel, Activity_pannel,ncol = 1)
```

---

#### Question 3: Do anteaters prefer to cross the roads through existing passages?

We were also interested in understanding whether giant anteaters preferred crossing roads through existing passage structures. To test for this we identified the number of crossings that were within the median error of a passage structure (20.27m).

```
#Load in the dataset of distances of crossings from passage structures
Passages <- read.csv("Passage_Distances.csv")

#Drop NAs
Passages <- Passages[!is.na(Passages$Passage_Distance),]

#Which crossings are within the median error of a passage structure
Passages[which(Passages$Passage_Distance <= 20.27),]
```

```
##       X   ID Passage_Distance
## 571 571 Beto        16.062860
## 572 572 Beto        11.121835
## 573 573 Beto         7.847453
## 574 574 Beto         7.281797
## 575 575 Beto         7.414816
## 576 576 Beto         8.666198
## 577 577 Beto        10.567850
## 578 578 Beto        11.965687
## 579 579 Beto        12.075959
## 580 580 Beto        10.984135
## 581 581 Beto         7.962624
## 582 582 Beto         8.583376
## 583 583 Beto        12.069507
## 584 584 Beto         9.479235
## 585 585 Beto         8.690240
## 586 586 Beto         9.566406
## 587 587 Beto         8.355456
## 588 588 Beto         9.060778
## 589 589 Beto        14.487649
```

```
length(which(Passages$Passage_Distance <= 20.27))
```

```
## [1] 19
```

```
#Median distance of crossings from passage structures
median(Passages$Passage_Distance)
```

```
## [1] 1733.273
```

```
#95% CIs on the median
n <- length(na.omit(Passages$Passage_Distance))
sort(na.omit(Passages$Passage_Distance))[round((n/2)*(1 + (1.96)/sqrt(n)),0)]
```

```
## [1] 1747.542
```

```
sort(na.omit(Passages$Passage_Distance))[round((n/2)*(1 - (1.96)/sqrt(n)),0)]
```

```
## [1] 1717.141
```

From this we can see that the median distance of road crossings from the nearest road passage was 1733.3m (95% CI: 1717.1 - 1747.5m).

---

#### Question 4: Do anteaters respond to roads differently than to natural barriers?

```
#Import the simulated crossings
SIMS <- read.csv("~/Dropbox (Personal)/UBC/Side_Projects/Arnaud_Anteaters/Scripts/Results/Simulated_Crossings_1000.csv")
SIMS <- aggregate(cbind(Num_Crossings, Stream_Crossings, Dirt_Crossings) ~ ID, data = SIMS, FUN = "mean")
data <- merge(data, SIMS, by = "ID", all.x = TRUE)

###########
# Paved road crossings vs simulated paved road crossings
t.test(data$Rd_Cr, data$Num_Crossings, paired = TRUE)
```

```
## 
##  Paired t-test
## 
## data:  data$Rd_Cr and data$Num_Crossings
## t = -3.805, df = 37, p-value = 0.0005153
## alternative hypothesis: true difference in means is not equal to 0
## 95 percent confidence interval:
##  -135.5943  -41.3636
## sample estimates:
## mean of the differences 
##               -88.47895
```

```
mean(data$Num_Crossings)/mean(data$Rd_Cr)
```

```
## [1] 3.313971
```

```
###########
# Dirt road crossings vs paved crossings
t.test(data$Rd_Cr, data$Drt_Cr, paired = TRUE)
```

```
## 
##  Paired t-test
## 
## data:  data$Rd_Cr and data$Drt_Cr
## t = -3.6702, df = 36, p-value = 0.0007797
## alternative hypothesis: true difference in means is not equal to 0
## 95 percent confidence interval:
##  -209.01219  -60.23105
## sample estimates:
## mean of the differences 
##               -134.6216
```

```
# Dirt roadcrossings vs simulated dirt road crossings
t.test(data$Dirt_Crossings, data$Drt_Cr, paired = TRUE)
```

```
## 
##  Paired t-test
## 
## data:  data$Dirt_Crossings and data$Drt_Cr
## t = 4.4646, df = 36, p-value = 7.612e-05
## alternative hypothesis: true difference in means is not equal to 0
## 95 percent confidence interval:
##   95.97067 255.73582
## sample estimates:
## mean of the differences 
##                175.8532
```

```
mean(na.omit(data$Dirt_Crossings))/mean(na.omit(data$Drt_Cr))
```

```
## [1] 2.048984
```

```
###########
# Stream crossings vs paved crossings
t.test(data$Rd_Cr, data$Strm_Cr, paired = TRUE)
```

```
## 
##  Paired t-test
## 
## data:  data$Rd_Cr and data$Strm_Cr
## t = -3.2737, df = 37, p-value = 0.002306
## alternative hypothesis: true difference in means is not equal to 0
## 95 percent confidence interval:
##  -375.46262  -88.37949
## sample estimates:
## mean of the differences 
##               -231.9211
```

```
# Stream crossings vs simulated stream crossings
t.test(data$Stream_Crossings, data$Strm_Cr, paired = TRUE)
```

```
## 
##  Paired t-test
## 
## data:  data$Stream_Crossings and data$Strm_Cr
## t = 0.56811, df = 37, p-value = 0.5734
## alternative hypothesis: true difference in means is not equal to 0
## 95 percent confidence interval:
##  -52.42163  93.27110
## sample estimates:
## mean of the differences 
##                20.42474
```

##### Recreate Figure 5

```
#Neg Binomial GLMs for plotting purposes
mod1 <- MASS::glm.nb(Rd_Cr ~ Dist_Rd, data = data)
mod2 <- MASS::glm.nb(Strm_Cr ~ Strm_Dist, data = data)
mod3 <- MASS::glm.nb(Drt_Cr ~ Drt_Dist, data = data)

#New datasets for plotting purposes
NEW_DATA1 <- data.frame(Dist_Rd = seq(0, 6, 0.01))
NEW_DATA2 <- data.frame(Strm_Dist = seq(0, 6, 0.01))
NEW_DATA3 <- data.frame(Drt_Dist = seq(0, 6, 0.01))

PRED <- predict(mod1, newdata = NEW_DATA1, type = "link", se = T)
PRED2 <- predict(mod2, newdata = NEW_DATA2, type = "link", se = T)
PRED3 <- predict(mod3, newdata = NEW_DATA3, type = "link", se = T)

#Paved road CIs
g1 <- 
  ggplot() +
  geom_line(aes(x=NEW_DATA1$Dist_Rd, y=exp(PRED$fit - 1.96*PRED$se.fit))) +
  geom_line(aes(x=NEW_DATA1$Dist_Rd, y=exp(PRED$fit + 1.96*PRED$se.fit)))

gg1 <- ggplot_build(g1)

# extract data for the loess lines from the 'data' slot
VAR <- data.frame(x = gg1$data[[1]]$x,
                  ymin = gg1$data[[1]]$y,
                  ymax = gg1$data[[2]]$y) 

#Stream CIs
g1 <- 
  ggplot() +
  geom_line(aes(x=NEW_DATA2$Strm_Dist, y=exp(PRED2$fit - 1.96*PRED2$se.fit))) +
  geom_line(aes(x=NEW_DATA2$Strm_Dist, y=exp(PRED2$fit + 1.96*PRED2$se.fit)))

gg1 <- ggplot_build(g1)

# extract data for the loess lines from the 'data' slot
Stream_VAR <- data.frame(x = gg1$data[[1]]$x,
                         ymin = gg1$data[[1]]$y,
                         ymax = gg1$data[[2]]$y) 

#Dirt road CIs
g1 <- 
  ggplot() +
  geom_line(aes(x=NEW_DATA3$Drt_Dist, y=exp(PRED3$fit - 1.96*PRED3$se.fit))) +
  geom_line(aes(x=NEW_DATA3$Drt_Dist, y=exp(PRED3$fit + 1.96*PRED3$se.fit)))

gg1 <- ggplot_build(g1)

# extract data for the loess lines from the 'data' slot
Dirt_VAR <- data.frame(x = gg1$data[[1]]$x,
                       ymin = gg1$data[[1]]$y,
                       ymax = gg1$data[[2]]$y) 

ggplot() +
  geom_point(data = data, aes(x = Dist_Rd, y = Rd_Cr, col = Road), size = 0.4) +
  geom_point(data = data, aes(x = Dist_Rd, y = Rd_Cr), col = "grey20", size = 0.5) +
  geom_point(data = data, aes(x = Strm_Dist, y = Strm_Cr), col = "#046C9A", size = 0.5) + 
  geom_point(data = data, aes(x = Drt_Dist, y = Drt_Cr), col = "#cdaa7d", size = 0.5) + 
  
  geom_line(aes(y = exp(PRED$fit), x =  NEW_DATA1$Dist_Rd), col = "grey20") +
  geom_ribbon(data = VAR, aes(x = x, ymin = ymin, ymax = ymax), fill="grey20", alpha = 0.2) +
  
  geom_line(aes(y = exp(PRED2$fit), x =  NEW_DATA2$Strm_Dist), col = "#046C9A") +
  geom_ribbon(data = Stream_VAR, aes(x = x, ymin = ymin, ymax = ymax), fill="#046C9A", alpha = 0.2) +
  
  geom_line(aes(y = exp(PRED3$fit), x =  NEW_DATA3$Drt_Dist), col = "#cdaa7d") +
  geom_ribbon(data = Dirt_VAR, aes(x = x, ymin = ymin, ymax = ymax), fill="#cdaa7d", alpha = 0.2) +
  
  scale_color_manual(values = c( "#046C9A", "#cdaa7d", "grey20"),
                     labels = c("Stream", "Unpaved road", "Paved road")) +
  
  theme_bw() +
  theme(panel.grid.major = element_blank(),
        panel.grid.minor = element_blank(),
        axis.title.y = element_text(size=10, family = "serif"),
        axis.title.x = element_text(size=10, family = "serif"),
        axis.text.y = element_text(size=8, family = "serif"),
        axis.text.x = element_text(size=8, family = "serif"),
        plot.title = element_text(hjust = -0.025, size = 10, family = "serif"),
        legend.position=c(0.8, 0.9), legend.title=element_blank(),
        legend.text=element_text(size=8, family = "serif"),legend.key.size = unit(0.2, "cm"),
        legend.key.width = unit(0.025, "cm"), legend.background=element_blank(), 
        legend.key = element_blank(), legend.text.align = 0) +
  xlab(expression(paste("Distance to barrier (km)"))) +
  ylab(expression(paste("Number of crossings"))) +
  coord_cartesian(ylim = c(-10,2000), xlim = c(0,2), expand = c(0,0.2))
```

#### Session Info

```
sessionInfo()
```

```
## R version 4.0.2 (2020-06-22)
## Platform: x86_64-apple-darwin17.0 (64-bit)
## Running under: macOS  10.16
## 
## Matrix products: default
## BLAS:   /Library/Frameworks/R.framework/Versions/4.0/Resources/lib/libRblas.dylib
## LAPACK: /Library/Frameworks/R.framework/Versions/4.0/Resources/lib/libRlapack.dylib
## 
## locale:
## [1] en_US.UTF-8/en_US.UTF-8/en_US.UTF-8/C/en_US.UTF-8/en_US.UTF-8
## 
## attached base packages:
## [1] stats     graphics  grDevices utils     datasets  methods   base     
## 
## other attached packages:
## [1] lubridate_1.7.10 nlme_3.1-152     DAAG_1.24        lattice_0.20-41 
## [5] MuMIn_1.43.17    gridExtra_2.3    ggplot2_3.3.3    metafor_2.4-0   
## [9] Matrix_1.3-2    
## 
## loaded via a namespace (and not attached):
##  [1] tidyselect_1.1.0    xfun_0.22           bslib_0.2.4        
##  [4] purrr_0.3.4         splines_4.0.2       colorspace_2.0-0   
##  [7] vctrs_0.3.6         generics_0.1.0      viridisLite_0.3.0  
## [10] htmltools_0.5.1.1   stats4_4.0.2        mgcv_1.8-34        
## [13] yaml_2.2.1          utf8_1.2.1          rlang_0.4.10       
## [16] jquerylib_0.1.3     pillar_1.5.1        glue_1.4.2         
## [19] withr_2.4.1         DBI_1.1.1           RColorBrewer_1.1-2 
## [22] jpeg_0.1-8.1        lifecycle_1.0.0     stringr_1.4.0      
## [25] munsell_0.5.0       gtable_0.3.0        evaluate_0.14      
## [28] labeling_0.4.2      latticeExtra_0.6-29 knitr_1.31         
## [31] pscl_1.5.5          fansi_0.4.2         highr_0.8          
## [34] Rcpp_1.0.6          scales_1.1.1        jsonlite_1.7.2     
## [37] farver_2.1.0        png_0.1-7           digest_0.6.27      
## [40] stringi_1.5.3       dplyr_1.0.5         grid_4.0.2         
## [43] tools_4.0.2         magrittr_2.0.1      sass_0.3.1         
## [46] tibble_3.1.0        crayon_1.4.1        pkgconfig_2.0.3    
## [49] MASS_7.3-53.1       ellipsis_0.3.1      assertthat_0.2.1   
## [52] rmarkdown_2.7       R6_2.5.0            compiler_4.0.2
```
