## Appendix S3 for "Roads as ecological traps for giant anteaters": Appendix_S3.html

Appendix S3 - Home range establishment on roads of different traffic volumes


### Appendix S3 - Home range establishment on roads of different traffic volumes

This document was created on March 30, 2021.

---

#### Home range establishment on roads of various traffic volumes

We were also interested in understanding how permeability might change between sites. To test for this, we calculated the ratio between the area that fell on either side of the nearest highway. This ratio ranges between zero, when home ranges are not bisected by roads and 1 when a road splits the home range area in half with an equal area on either side of the road.

Figures depicting these relationships are shown below.

We then modelled the relationship between the ratio of home range areas that fell on either side of the highways across the different paved roads (and therefore traffic volume).

```
#Load in the necessary packages
library(nlme)

#Import the dataset of individual movement metrics
data <- read.csv("Results/Anteater_Results_Final.csv")

#Re-order based on location
data <- data[order(data$Road),]

#Convert factor variables to factors
data$Sex <- as.factor(data$Sex)
data$Road <- as.factor(data$Road)

#Road specific sample sizes
table(data = data[which(data$Dist_Rd < 2),"Road"])
```

```
## data
## BR_267  BR262 MS-040 
##     13      7      7
```

```
boxplot(Split ~ Road, data[which(data$Dist_Rd < 2),],
        xlab = "Road")
```

```
#Are animals establishing HRs on roads differently?
mod <- glm(Split ~ Road, data = data[which(data$Dist_Rd < 2),],
           family = binomial)

summary(mod)
```

```
## 
## Call:
## glm(formula = Split ~ Road, family = binomial, data = data[which(data$Dist_Rd < 
##     2), ])
## 
## Deviance Residuals: 
##     Min       1Q   Median       3Q      Max  
## -0.7255  -0.4519  -0.3132   0.2066   2.0283  
## 
## Coefficients:
##             Estimate Std. Error z value Pr(>|z|)  
## (Intercept)  -2.2162     0.9316  -2.379   0.0174 *
## RoadBR262     0.6212     1.3735   0.452   0.6511  
## RoadMS-040    1.0157     1.2927   0.786   0.4320  
## ---
## Signif. codes:  0 '***' 0.001 '**' 0.01 '*' 0.05 '.' 0.1 ' ' 1
## 
## (Dispersion parameter for binomial family taken to be 1)
## 
##     Null deviance: 8.8888  on 26  degrees of freedom
## Residual deviance: 8.2453  on 24  degrees of freedom
## AIC: 19.393
## 
## Number of Fisher Scoring iterations: 5
```

```
plot(coef(summary(mod))[,1],
     axes = F,
     pch = 16,
     ylim = c(-5,4),
     xlab = "",
     ylab = expression(beta),
     cex = 0.7)
text(x =  1:3, y = par("usr")[3]-0.05, srt = 40, adj = 1,
     labels = row.names(coef(summary(mod))), xpd = TRUE, cex = 0.7)
segments(x0 = 1:4, x1 = 1:4, y0 = confint(mod)[,1], y1 = confint(mod)[,2])
axis(2)
abline(h = 0, col = "grey", lty = 2)
```

The mean daily traffic volume acroos these roads were: MS-040 - 603 vehicles/day; BR-262 - 3206 vehicles/day; BR-267 - 4285 vehicles/day. From our results we see that individuals were more likely to establish home ranges on the lower traffic MS-040 than the highest traffic BR-267. Similarly, individuals were more likely to establish home ranges on the BR-262 than BR-267. None of these relationships were significant however. It does appear if two potential outliers from BR 267 might be influencing the results. Repeating the analyses without these outliers yields the following.

```
#Outlier removal for BR 267
data <- data[-which(data$Road == "BR_267" & data$Split == 0.958850287),]
data <- data[-which(data$Road == "BR_267" & data$Split == 0.158780748),]

boxplot(Split ~ Road, data[which(data$Dist_Rd < 2),],
        xlab = "Road")
```

```
#Are animals establishing HRs on roads differently?
mod <- glm(Split ~ Road, data = data[which(data$Dist_Rd < 2),],
           family = binomial)

summary(mod)
```

```
## 
## Call:
## glm(formula = Split ~ Road, family = binomial, data = data[which(data$Dist_Rd < 
##     2), ])
## 
## Deviance Residuals: 
##     Min       1Q   Median       3Q      Max  
## -0.7255  -0.1714  -0.1064   0.2249   0.5623  
## 
## Coefficients:
##             Estimate Std. Error z value Pr(>|z|)  
## (Intercept)   -4.214      2.516  -1.675    0.094 .
## RoadBR262      2.619      2.711   0.966    0.334  
## RoadMS-040     3.013      2.671   1.128    0.259  
## ---
## Signif. codes:  0 '***' 0.001 '**' 0.01 '*' 0.05 '.' 0.1 ' ' 1
## 
## (Dispersion parameter for binomial family taken to be 1)
## 
##     Null deviance: 5.3719  on 24  degrees of freedom
## Residual deviance: 2.7830  on 22  degrees of freedom
## AIC: 12.594
## 
## Number of Fisher Scoring iterations: 7
```

Outliers do not seem to be influencing this relationship, although the p values did move closer to significance. To check explicitly for an effect of traffic volume, we also modelled the relationship between the ratio of home range areas that fell on either side of the highways and the mean daily traffic volume.

```
data$Traffic <- 0
data[which(data$Road == "BR262"),"Traffic"] <- 3206
data[which(data$Road == "BR_267"),"Traffic"] <- 4285
data[which(data$Road == "MS-040"),"Traffic"] <- 603


#Are animals establishing HRs on roads differently?
mod <- glm(Split ~ Traffic, data = data[which(data$Dist_Rd < 2),],
           family = binomial)

summary(mod)
```

```
## 
## Call:
## glm(formula = Split ~ Traffic, family = binomial, data = data[which(data$Dist_Rd < 
##     2), ])
## 
## Deviance Residuals: 
##     Min       1Q   Median       3Q      Max  
## -0.7685  -0.3265  -0.1885   0.1484   0.8684  
## 
## Coefficients:
##               Estimate Std. Error z value Pr(>|z|)
## (Intercept) -0.7676852  1.0322621  -0.744    0.457
## Traffic     -0.0004988  0.0004009  -1.244    0.213
## 
## (Dispersion parameter for binomial family taken to be 1)
## 
##     Null deviance: 5.3719  on 24  degrees of freedom
## Residual deviance: 3.7264  on 23  degrees of freedom
## AIC: 10.561
## 
## Number of Fisher Scoring iterations: 5
```

```
boxplot(Split ~ Traffic, data[which(data$Dist_Rd < 2),],
        xlab = "Mean Daily Traffic Volume")
```

Again, although there is a negative relationship between traffic volume and the ratio of home range areas that fell on either side of the highways, this effect was non-significant, likely due to small sample sizes and substantial inter-individual variation

#### Session Info

```
sessionInfo()
```

```
## R version 4.0.2 (2020-06-22)
## Platform: x86_64-apple-darwin17.0 (64-bit)
## Running under: macOS  10.16
## 
## Matrix products: default
## BLAS:   /Library/Frameworks/R.framework/Versions/4.0/Resources/lib/libRblas.dylib
## LAPACK: /Library/Frameworks/R.framework/Versions/4.0/Resources/lib/libRlapack.dylib
## 
## locale:
## [1] en_US.UTF-8/en_US.UTF-8/en_US.UTF-8/C/en_US.UTF-8/en_US.UTF-8
## 
## attached base packages:
## [1] stats     graphics  grDevices utils     datasets  methods   base     
## 
## other attached packages:
##  [1] nlme_3.1-152     plyr_1.8.6       rgeos_0.5-5      raster_3.4-5    
##  [5] maptools_1.0-2   rgdal_1.5-23     sp_1.4-5         geosphere_1.5-10
##  [9] lubridate_1.7.10 proj4_1.0-10.1   ctmm_0.6.1      
## 
## loaded via a namespace (and not attached):
##  [1] Rcpp_1.0.6        knitr_1.31        magrittr_2.0.1    MASS_7.3-53.1    
##  [5] lattice_0.20-41   R6_2.5.0          rlang_0.4.10      highr_0.8        
##  [9] stringr_1.4.0     tools_4.0.2       grid_4.0.2        xfun_0.22        
## [13] jquerylib_0.1.3   htmltools_0.5.1.1 yaml_2.2.1        digest_0.6.27    
## [17] sass_0.3.1        codetools_0.2-18  evaluate_0.14     rmarkdown_2.7    
## [21] stringi_1.5.3     compiler_4.0.2    bslib_0.2.4       generics_0.1.0   
## [25] jsonlite_1.7.2    foreign_0.8-81
```
